# Towards Useful and Private Synthetic Omics: Community Benchmarking of Generative Models for Transcriptomics Data

**DOI:** 10.64898/2026.03.02.707794

**Authors:** Hakime Öztürk, Tejumade Afonja, Joonas Jälkö, Ruta Binkyte, Pablo Rodriguez-Mier, Sebastian Lobentanzer, Andrew Wicks, Jules Kreuer, Sofiane Ouaari, Nico Pfeifer, Shane Menzies, Sikha Pentyala, Daniil Filienko, Steven Golob, Patrick McKeever, Jineta Banerjee, Luca Foschini, Martine De Cock, Julio Saez-Rodriguez, Mario Fritz, Oliver Stegle, Antti Honkela

## Abstract

**Background:** The synthesis of anonymized data derived from real-world cohorts offers a promising strategy for regulatory-compliant and privacy-preserving biological data sharing, potentially facilitating model development that can improve predictive performance. However, the extent to which generative models can preserve biological signals while remaining resilient to adversarial privacy attacks in high-dimensional omics contexts remains underexplored. To address this gap, the CAMDA 2025 Health Privacy Challenge launched a community-driven effort to systematically benchmark synthetic and privacy-preserving data generation for bulk RNA-seq cohorts.

**Results:** Building on this initiative, we systematically benchmarked 11 generative methods across two cancer cohorts (∼1,000 and ∼5,000 patients) over 978 landmark genes. Methods were evaluated across complementary axes of distributional fidelity, downstream utility, biological plausibility and empirical privacy risk, with emphasis on trade-offs between vulnerability to membership inference attacks (MIA) and other evaluation dimensions. Expressive deep generative models achieved strong predictive utility and differential expression recovery, but were often more vulnerable to membership inference risk. Differentially private methods improved resistance to attacks at the cost of reduced utility, while simpler statistical approaches offered competitive utility with moderate privacy risk and fast training.

**Conclusions:** Synthetic bulk RNA-seq quality is inherently multi-dimensional and shaped by trade-offs between utility, biological preservation and privacy. Our results indicate that differences in model architecture drive distinct trade-offs across these axes, suggesting that model choice should align with dataset characteristics, intended downstream use and privacy requirements. Privacy risk should also be assessed using multiple complementary attack methods and, where possible, formal differential privacy protection.

## Background

The unprecedented availability of health data facilitates understanding and predicting disease and patient trajectories. Machine and deep learning-based methods learn latent representations from large amounts of datasets, potentially improving generalization to unknown and rare cases. However, the improved performance of these methods often depends on large parameter spaces that are prone to memorisation of data points, posing challenges for patient privacy. While on one hand, the benefit of sharing and utilizing large scale health care data promises increased utility, it is also crucial to ensure patient privacy is not disclosed. Provable privacy preservation can be implemented through differential privacy (DP)^1, 2^. Federated learning (FL)^3^ and synthetic data generation^4, 5, 6^ can provide some protection alone, or be combined with DP.

Synthetic data generation enables privacy preservation through generating data points that approximate the distribution of the real data, although synthetic data can still be vulnerable to privacy attacks^7,8^. This not only enables safer data sharing but also provides an opportunity in data augmentation and building fair models ^9–11^. The utility of synthetic data generation for Electronic Health Records^12,13,14,15^, genomics^16^ and gene expression^17,18^ has been investigated through different evaluation metrics. The “Hide and Seek Privacy Challenge”, organized as a part of NeurIPS 2020, provided a platform for teams to generate synthetic longitudinal intensive care data and to launch privacy attacks on the generated data to compete against each other ^19^. The results of the competition highlighted the complexities of balancing utility and privacy in synthetic biological datasets, emphasizing the need for developing robust privacy-preserving generative methods and for reliable evaluation metrics.

Although recent studies have been examining trade-offs among utility, fidelity, and privacy metrics ^17, 20,21–23^, the way these metrics interact across diverse generative-model families remains underexplored, particularly for gene-expression datasets, which are high dimensional and exhibit biologically structured patterns, including gene-gene co-expression dependencies and condition-specific differential expression. Bulk RNA-seq is highly relevant in this context because it is widely available across cohorts, supports different downstream analyses, and is commonly represented as tabular format, lowering the barrier to study utility-privacy trade-offs under realistic data-sharing constraints. However, it remains unclear whether improvements in global distributional fidelity translate into downstream task utility, whether biological signal preservation aligns with either of these objectives, and how privacy risks evolve along these axes. This gap complicates the interpretation of benchmark results and the design of synthetic data pipelines for biomedical research.

The Health Privacy Challenge (https://benchmarks.elsa-ai.eu/?ch=4), organized in conjunction with the Critical Assessment of Massive Data Analysis (CAMDA) 2025, was designed to collectively explore privacy-preserving synthetic data generation in the context of gene expression datasets. The first track of the challenge focused on private synthetic bulk RNA-seq generation for two The Cancer Genome Atlas (TCGA) datasets, where “blue” teams generated biologically meaningful and private synthetic datasets, and “red” teams investigated the vulnerabilities of these generated datasets at the level of membership inference attack (MIA)^24^. In MIA, an adversary aims to determine the presence of an individual’s data in the training set of a generative model. Although membership inference attacks can pose a privacy risk in regulated or stigmatized sensitive datasets, they often reveal limited information and are commonly used to evaluate vulnerability to more serious attacks such as attribute inference and data reconstruction ^25^. The challenge setting therefore provides a controlled environment in which utility, fidelity, biological signal preservation, and privacy risk can be evaluated under comparable conditions. In this work, we build on the first track’s “blue” team task of the Health Privacy Challenge 2025 which addresses private synthetic bulk RNA-seq data generation and received the majority of submissions (Supplementary Materials). Extending prior evaluations focused primarily on differentially private models^17^, we benchmark a broader range of generative approaches and assess their performance across complementary dimensions. We harmonize community submissions with suggested baseline and post-challenge methods to benchmark models for private synthetic bulk gene-expression generation. Rather than emphasizing model ranking or competitive outcomes, we analyze how different classes of metrics such as utility, global fidelity, biological preservation, and privacy relate to one another, where they diverge, and how sensitive their conclusions are to model architecture, dataset characteristics, threat models and differential privacy constraints. This enables explicit quantification of trade-offs and provides actionable guidance for model selection in realistic cohort settings. Overall, this study, (i) provides a summary of the current landscape of approaches, (ii) establishes a reference point for future research and community challenges, and (iii) highlights the need for multi-dimensional evaluation practices for synthetic transcriptomic datasets.

## Results

We generated synthetic bulk RNA-seq data based on two patient-level cohorts, TCGA-BRCA and TCGA-COMBINED, using 8 distinct generative architectures across different privacy settings (11 methods in total), and assessed the quality of the generated synthetic datasets across four key dimensions: distributional fidelity, downstream utility biological plausibility, and privacy (Fig.1). The complete list of all metrics considered in this benchmarking study is provided in Supplementary Table S1.

**Figure 1.**
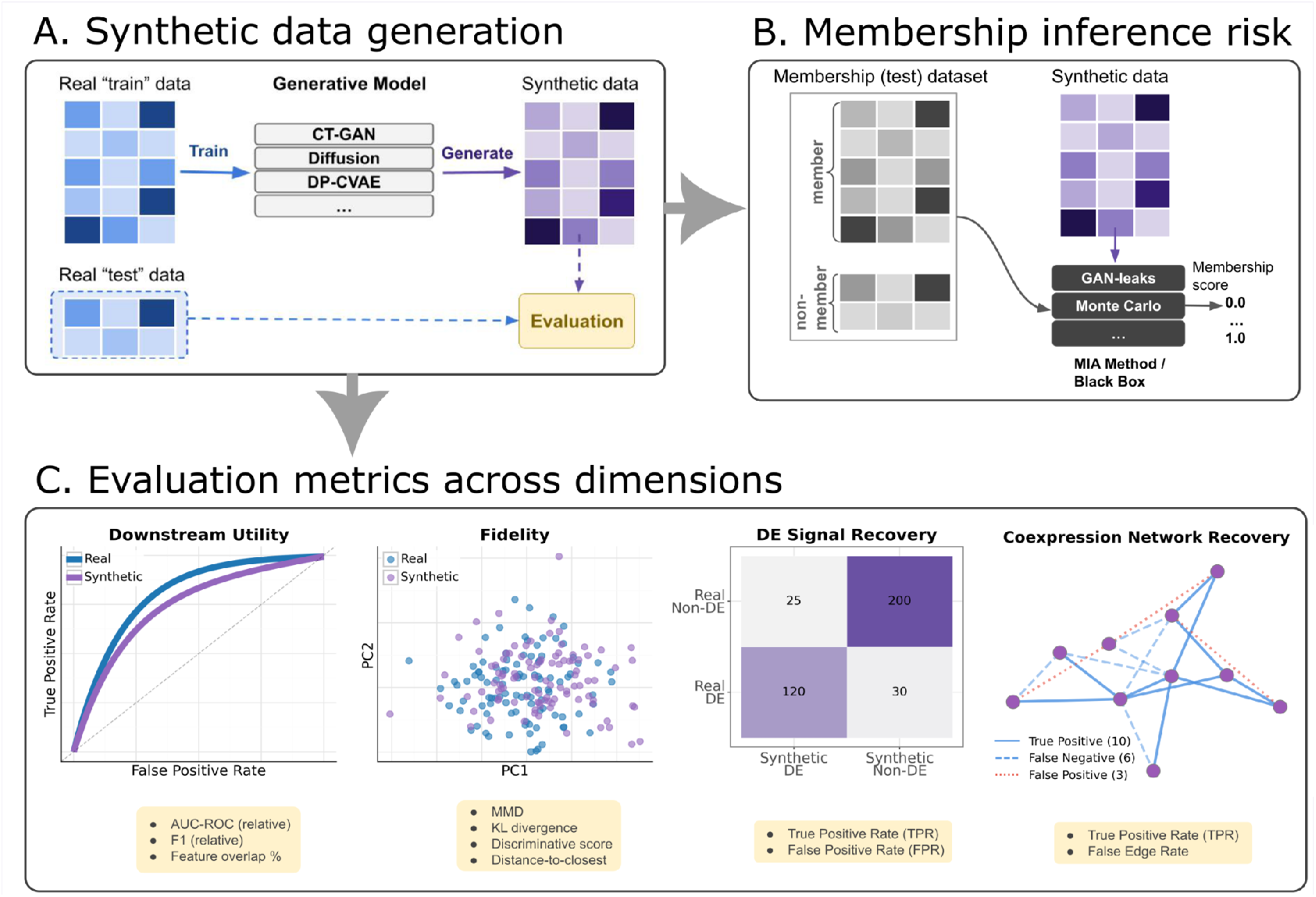
Benchmarking framework for evaluating generative models on synthetic bulk RNA-seq data generation. **(A) Synthetic data generation workflow**. Real bulk RNA-seq data are partitioned into training and test sets. Generative models (including differential privacy methods, denoted by DP) are trained exclusively on the training data to generate synthetic samples. The test set is held out for evaluation purposes and not used during model training or data generation. **(B) Privacy risk assessment via membership inference attacks (MIA)**. Real data is labelled as members (samples present in the training set) and non-members (held-out (test) samples) to generate a membership (query) test dataset. Both membership test dataset and synthetic data are input to black-box MIA methods (e.g. GAN-leaks ^26^ etc.) to predict membership scores, which quantify the risk of identifying whether specific samples were used in model training. MIA performance is quantified by AUC-ROC and true positive rate (TPR) fixed at specific false positive rate (FPR) value (i.e. TPR@FPR=0.1) metrics to assess potential disclosure of training samples. **(C) Evaluation metrics across dimensions**. Models are evaluated across complementary dimensions: (i) Distributional Fidelity: statistical similarity between real and synthetic distributions assessed via maximum mean discrepancy (MMD), Kullback-Leibler (KL) divergence, distance-to-closest real record, and discriminator-based metrics; (ii) Downstream Utility: preservation of predictive performance measured by relative AUC-ROC, F1 score, and overlap of important features when training classifiers on synthetic versus real data; (iii) Biological Plausibility: recovery of biologically meaningful patterns including differential expression (DE) at controlled false positive rate (FPR ≤ 0.05) (true positives, false negatives, false positives etc.), and co-expression network edge recovery (true positives in blue, false negatives in dashed blue, false positives in dotted red).

### Distributional fidelity and separability of synthetic bulk RNA-seq across models

Fidelity measures how closely the synthetic data reproduce the global and local structure of the real data. We evaluated distributional fidelity using complementary global (Maximum Mean Discrepancy (MMD) and Kullback-Leibler (KL) divergence), local (Distance-to-closest record) and discriminative (classifier-based) metrics, computed against both the real train and the real held-out test data (Methods). For interpretability, KL, MMD, and discriminative score metrics were inverted such that higher values indicate closer alignment between real and synthetic distributions. We report inverted MMD (test) and inverted KL (test) to emphasize generalization of synthetic data to held-out real data, as divergence metrics computed against the (real) training data were strongly correlated with their test-based counterparts, indicating partial redundancy (Supplementary Figures S1-S2). Figure 2 summarizes these fidelity metrics across generative models.

**Figure 2.**
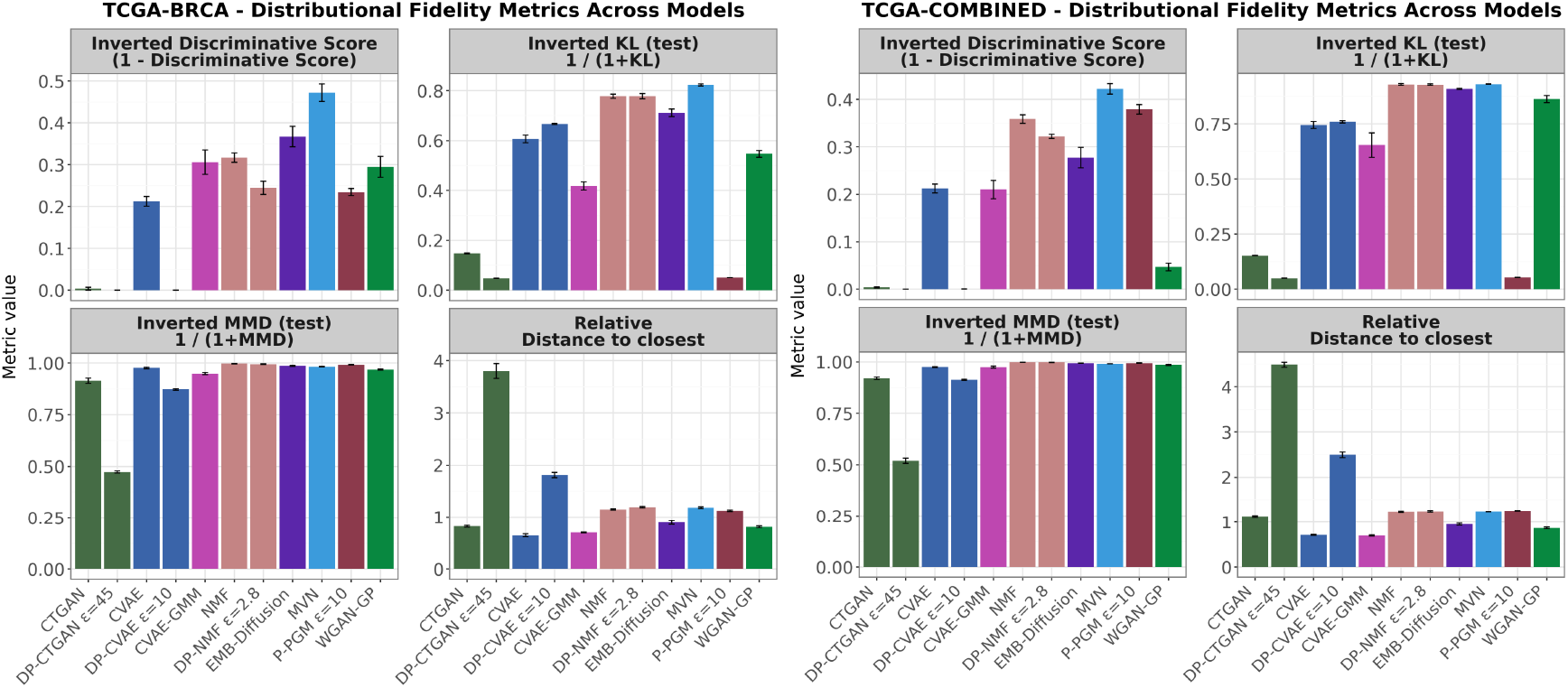
Fidelity metrics for BRCA and COMBINED datasets. Four metrics are shown in separate facets: MMD (test) and KL divergence (test) are computed against the real test dataset and visualized after inversion (e.g., 1/(1+KL)), while discriminative scores are computed against the training dataset and visualized as 1 - discriminative score, so that higher values indicate more difficult separation between synthetic and real (i.e. higher fidelity). Distance-to-closest record is reported relative to the real data baseline (values >1 indicate synthetic samples are farther from the training set than real test samples). Differentially private (DP) and non-DP variants of the same model share the same color. Each bar represents the mean value of the metric across five folds, with error bars showing the standard deviation.

Although most methods exhibit similar inverted MMD scores, differences emerge in KL divergence metric. Statistical methods, including the Multivariate Normal (MVN) and Non-negative Matrix Factorization (NMF) variants, achieve comparatively higher inverted KL divergence, followed by the Embedded Diffusion model on both datasets. Differentially Private (DP) P-PGM (ϵ = 10 ) displays a surprisingly low inverted KL divergence despite having a relatively high inverted MMD score. This indicates that, although synthetic data matches multivariate structure of real samples under RBF kernel (high inverted MMD), it might fail to reproduce gene-wise marginal distributions (low inverted KL). As a result, P-PGM generates samples that appear realistic in overall multivariate structure but deviate from real data at the level of individual genes. This discrepancy is consistent in both datasets and might contribute to reduced performance in downstream evaluations that depend on gene-wise distributions such as co-expression and differential expression recovery, which are examined in the subsequent sections.

Classifier-based discriminative performance (F1) and relative distance-to-closest mostly follow similar ordering across models. Deep generative DP-models, including DP-CTGAN, and DP-CVAE, tend to generate samples farther from any individual real record while simultaneously being more easily separable from real data by a classifier (indicated by low inverted discriminative score, implying low indistinguishability). MVN achieves the highest inverted discriminative score across both datasets, indicating that its synthetic samples are particularly difficult to distinguish from real data. MVN also lies in the high-fidelity region of both MMD and KL, suggesting strong global distributional alignment despite its simple parametric form. Among deep generative models without formal privacy guarantee, the Embedded Diffusion model attains the highest fidelity levels. Complete fidelity metric scores for each method across both datasets can be found in Supplementary Tables S2-S3.

### Downstream predictive utility of synthetic bulk RNA-seq datasets

Utility refers to the extent to which a synthetic dataset can replicate the performance of real data in a downstream task, as well as how well it preserves the critical features underlying predictive decisions. We assessed this using the “train-on-synthetic, test-on-real” (TSTR) scheme, where the models trained on synthetic data were evaluated on real test data compared to the models trained on real data tested on the held-out real data (TRTR).

In our study, the downstream tasks are molecular subtype prediction for the TCGA-BRCA dataset and cancer type prediction for the TCGA-COMBINED dataset. Performance is summarized using AUROC, F1, and AU-PR, reported as relative performance (TSTR/TRTR), with ratios closer to one indicating higher utility. We additionally report the feature overlap (%), using the top predictive genes from each class, aggregated globally (Methods). Utility metric values for each method across both datasets can be found in Supplementary Tables S2-S3.

Figure 3 shows that AUROC, AU-PR, and F1 relative scores exhibit highly consistent patterns across datasets. CTGAN variants, together with DP-CVAE, NMF variants, display the lowest relative scores across most metrics, indicating substantial loss of downstream predictive signal when trained on synthetic data. In contrast, MVN and CVAE, CVAE-GMM and Embedded Diffusion achieve substantially higher relative scores across most metrics in both datasets. These same models also exhibit the highest feature overlap proportions, indicating better preservation of the genes that are relevant for the downstream classification task. We also note that P-PGM consistently outperforms other formally DP models and achieves relative scores that are comparable to, or slightly lower than, those of non-DP models. However, its proportion of overlapping important features is substantially lower than that of non-DP models across both datasets.

**Figure 3.**
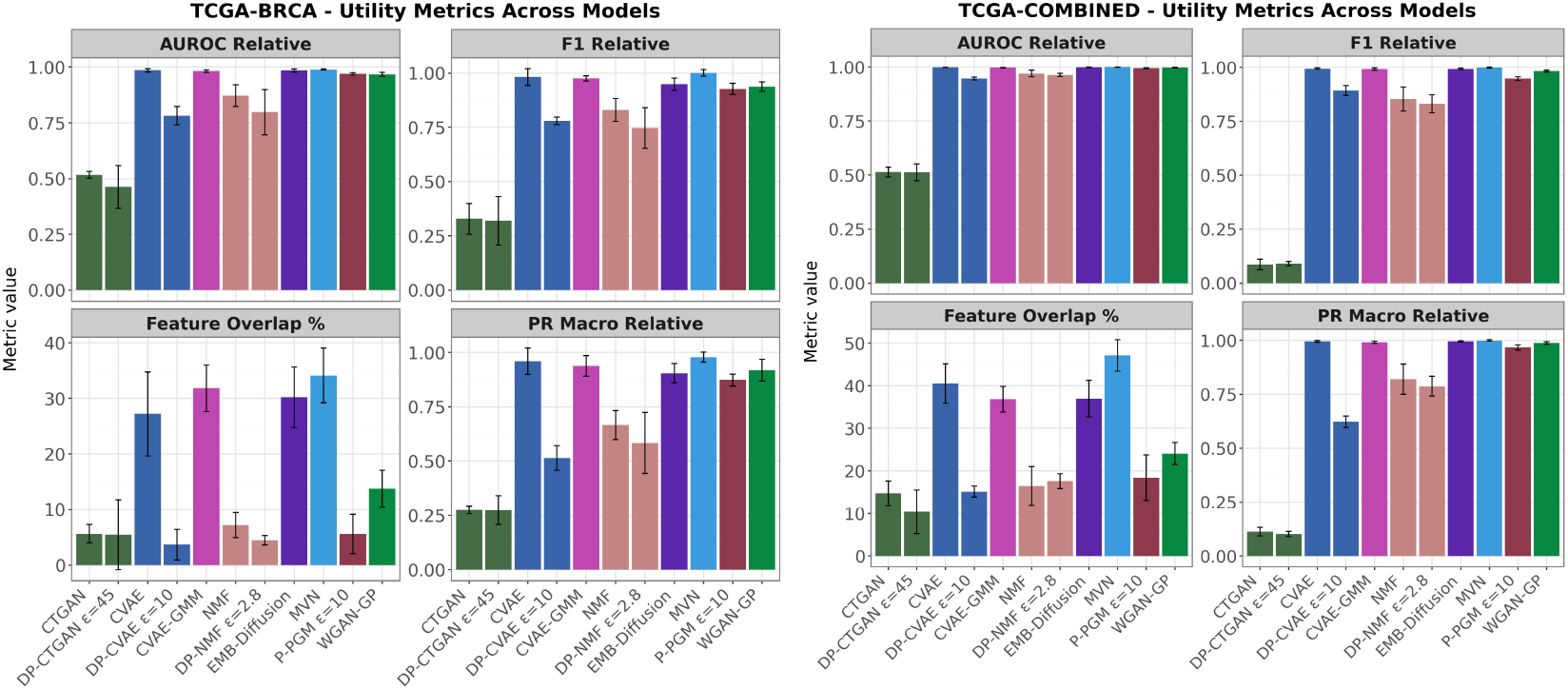
Utility metrics for BRCA and COMBINED datasets. Four metrics are shown, each in a separate facet: Relative AUROC, relative F1, relative precision-recall (macro), and important feature overlap proportion (%). For the relative metrics, values closer to 1 indicate better utility, i.e. the performance of the synthetic downstream prediction getting closer to real. A higher feature overlap proportion indicates a stronger match between the predictive features identified by synthetic downstream models and those identified by real downstream models. Differentially private (DP) and non-DP variants of the same generative model are represented with the same color for consistency. Each bar represents the mean value of the metric across five folds, with error bars showing the standard deviation.

### Differential expression and co-expression recovery across synthetic RNA-seq datasets

We investigated how well the synthetic data captures gene expression dependencies via co-expression and differential expression recovery patterns.

#### Differential expression (DE)

We evaluated DE recovery using paired comparisons between groups (e.g., molecular subtypes for TCGA-BRCA and cancer types for TGCA-COMBINED; see Methods). Figure 4 shows DE recovery under stringent criteria (effect size ΔVST > 0.5 and false positive rate (FPR) ≤ 0.05) averaged across comparisons and folds, illustrated as true positive rate (TPR) against FPR in a ROC-style format. Up- and down-regulated genes are assessed separately to capture directional differences. Supplementary Figures S3-S4 show TPR performance for each pairwise comparison in the BRCA and COMBINED datasets.

**Figure 4:**
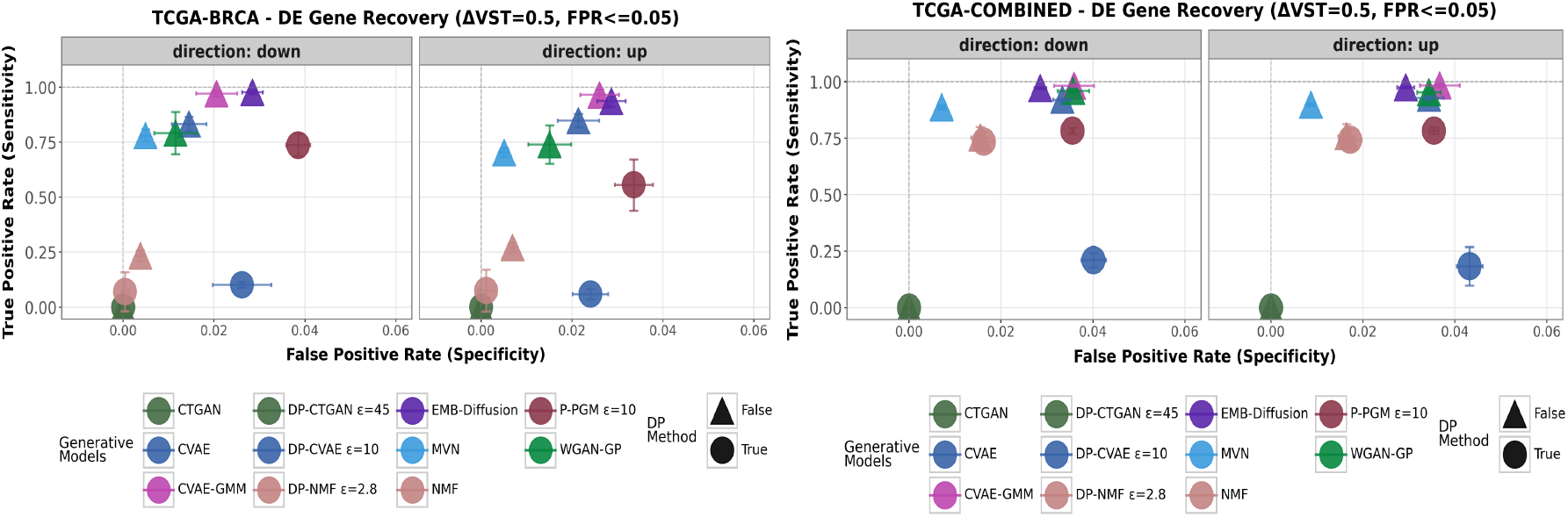
Differential Expression (DE) recovery analysis for BRCA and COMBINED datasets. Each colored dot represents the mean metric values for a generative model across five folds, with error bars indicating the corresponding standard deviation along each axis. Differentially private (DP) and non-DP versions of the same model share the same color but are represented with different shapes. The y-axis shows the true positive rate (TPR) and the x-axis shows the false positive rate (FPR), forming a ROC-style curve where better models appear toward the top-left corner. Points The analysis is valid for effect size threshold set to 0.5 and FPR ≤ 0.05.

Across both datasets, up- and down-regulation patterns are broadly similar, with model rankings largely consistent between directions. Deep generative models CVAE-GMM and Embedded Diffusion achieve the highest DE recovery, followed closely by CVAE and WGAN-GP. Among statistical models, MVN attains the highest TPR and slightly lower FPR than deep models. P-PGM, while outperforming other DP-constrained models in both datasets, remains less successful than non-DP models and shows direction-specific behaviour in the BRCA dataset, with lower TPR for up-regulated genes than for down-regulated genes.

Moving from the smaller BRCA dataset to larger COMBINED dataset, FPR is consistently elevated across most models. NMF variants, which exhibit poor DE TPR in the BRCA dataset, improve substantially in the COMBINED dataset.

In our ablation experiments with effect-size thresholds of 0.0 and 0.5 for BRCA, and 0.0, 0.5, and 1.0 for COMBINED (Supplementary Figures S4-S5), a consistent trend emerges across models: FPR decreases as the effect size threshold increases in both datasets. In TCGA-BRCA, CVAE variants that introduce spurious correlations and P-PGM affected by DP-induced noise only pass the FPR cutoff once the DE signal is sufficiently strong (effect size ≥ 0.5).

Overall, consistent with their high downstream utility, expressive deep generative models achieve strong DE recovery under stringent FPR control, while simpler parametric models such as MVN also perform well. DP constraints generally reduce DE recovery TPR, particularly for subtle DE signals, as reflected in lower performance on BRCA (molecular subtypes) compared to COMBINED (cancer types).

#### Gene co-expression

We assessed co-expression recovery by comparing gene-gene correlation networks between real and synthetic data under correlation thresholds implemented in hCoCena package, with cut-offs of 0.3 and 0.5 to investigate model robustness and biological fidelity (Methods). Higher true positive rates indicate stronger preservation of biological co-expression patterns, whereas false edge rates reflect spurious relationships introduced by the generative model, computed as the number of spurious edges in the synthetic network, normalized by the number of true edges in the real network (Fig. 5).

**Figure 5:**
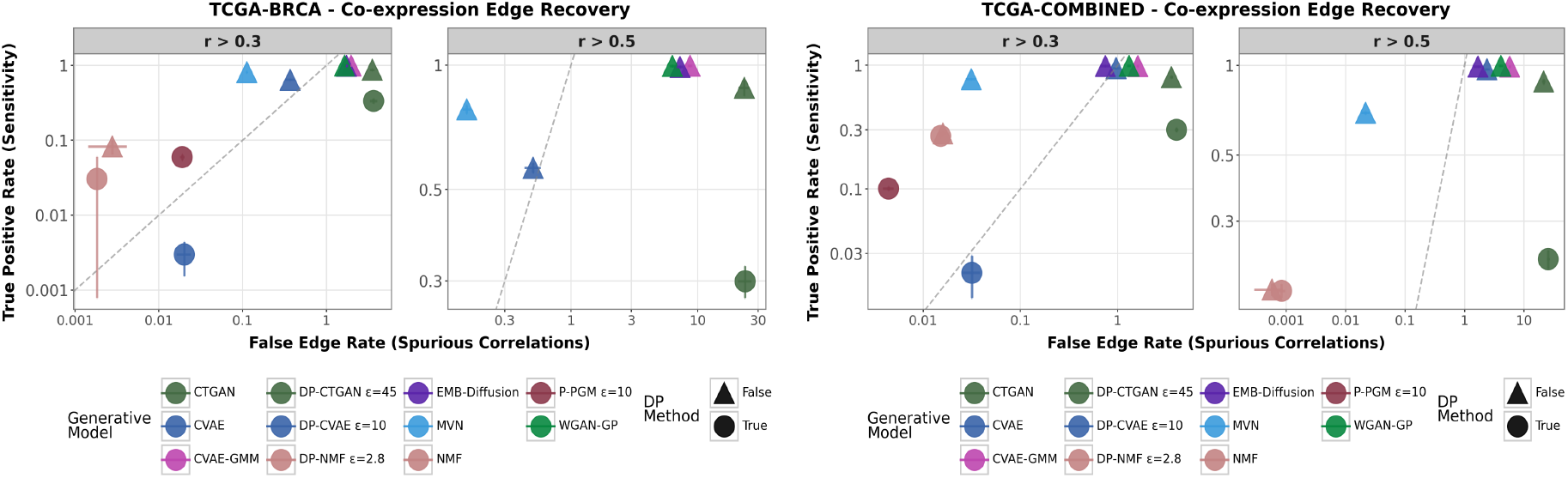
Co-expression recovery analysis for the BRCA and COMBINED datasets. Each facet corresponds to the correlation threshold applied to gene-gene pairs. Each colored point represents the mean metric values for a generative model across five folds, with error bars indicating the corresponding standard deviation along each axis. Differentially private (DP) and non-DP variants share the same color but are distinguished by shape. The y-axis shows the true positive rate (fraction of real edges recovered in the synthetic network), and the x-axis shows the false edge rate (spurious edges in the synthetic network, normalized by the number of true edges in the real network). The gray dashed line indicates a reference linear trend.

In the BRCA dataset, increasing the correlation threshold from 0.3 to 0.5 challenges several models. P-PGM, DP-CVAE, and NMF variants fail to reliably re-construct the real co-expression network at the stricter cutoff, reflecting in their disappearance from the plot. Deep generative models such as CVAE-GMM, Embedded Diffusion and WGAN-GP maintain relatively high TPR but at the cost of introducing substantially more spurious edges under a stringent threshold. In contrast, MVN consistently preserves a very low false edge rate across cutoffs, although it exhibits a notable drop in TPR as the threshold becomes more stringent.

A broadly similar pattern is also observed in the COMBINED dataset. When the cutoff increases from 0.3 to 0.5, P-PGM, and DP-CVAE again fail to reliably recover the real co-expression network. CVAE-GMM, Embedded Diffusion and WGAN-GP consistently exhibit high TPR while still introducing a high amount of spurious edges. Notably, however, their false edge rate at cutoff 0.5 is substantially lower in COMBINED than BRCA. This might reflect the increased stability of correlation estimates in greater sample size. In addition, higher heterogeneity in COMBINED, which spans multiple cancer types, might induce stronger and structured co-variance patterns that are easier to recover.

These results illustrate a trade-off: deep generative models can produce dense networks that preserve many true edges but also generate many spurious interactions, whereas simpler models like MVN favor sparse networks with fewer false edges but may lose some true connections under stringent thresholds. This balance is further influenced by dataset characteristics.

### Privacy risk under membership inference attacks

We evaluated the privacy vulnerability of the synthetic datasets using multiple black-box membership inference attacks (MIAs) that only require access to the released synthetic data (Methods). These include distance-, density estimation- and classifier confidence-based attacks. For TCGA-COMBINED, we additionally evaluated reference-based attacks that utilize a reference dataset with similar distribution. Attack performance was reported using a threshold-independent classification metric (i.e. AUC-ROC) and an operational metric (TPR at fixed FPR levels of 0.1). While distance-based attacks generally show aligned classification and operational metrics, other attack types display either distinct clustering or no consistent pattern. Complete performances and correlation analyses across all attack methods are provided in Supplementary Figures S7-S12.

Figure 6 demonstrates two representative attacks: confidence-based random forest (RF) and GAN-leaks, where the latter is evaluated with and without auxiliary dataset. Across attacks and datasets, three distinct privacy regimes emerge: random guessing level, moderate and elevated vulnerability.

**Figure 6.**
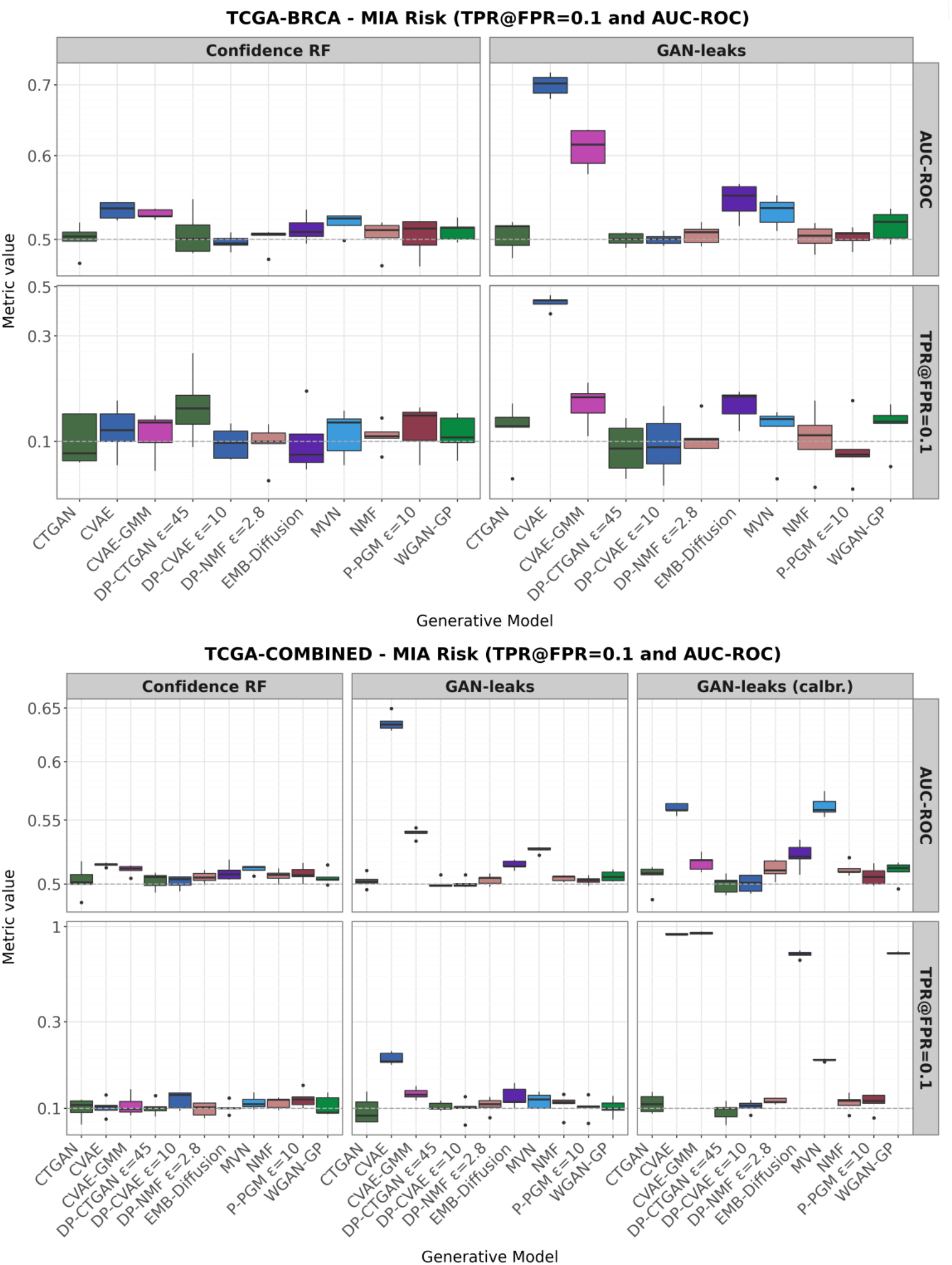
Membership inference attack (MIA) performance on the BRCA and COMBINED datasets. We report two complementary privacy risk metrics: area under the ROC curve (AUC-ROC) and true positive rate at a fixed false positive rate of 0.1 (TPR@FPR=0.1). Facets correspond to the evaluated metric and the MIA algorithm. For the BRCA dataset, results are shown for GAN-leaks and confidence-based Random Forest attacks; for the COMBINED dataset, we additionally include the calibrated GAN-leaks variant. Each boxplot summarizes performance across five cross-validation folds. Generative models are color-coded, with differentially private (DP) and non-DP variants of the same model sharing the same color.

#### Random-guessing level attack performance

Formally differentially private (DP) methods, including DP-CVAE (ϵ = 10 ), DP-NMF (ϵ = 2.8 ), and P-PGM (ϵ = 10) consistently operate close to random-guessing level (e.g. TPR@FPR=0.1 ∼ 0.1). This pattern holds across attack variants, and is particularly stable for DP-CVAE and P-PGM under the calibrated GAN-leaks attack in the COMBINED dataset, which highlights the importance of DP guarantees in adversarial settings that include auxiliary reference data. It’s also important to note that DP is applied at different stages in these models, for instance, gradients (DP-CVAE), factor matrices (DP-NMF) and probabilistic parameters (P-PGM), which might influence how privacy constraints interact with attack algorithms. Nonetheless, these results confirm that explicit DP constraints demonstrate effective empirical mitigation of MIA risk.

Among non-DP methods, NMF also exhibits lower vulnerability across settings. Given its constrained factorization structure and comparatively moderate utility, this suggests that limited model expressiveness may inherently restrict the amount of membership-relevant signal encoded in the synthetic data.

A cautionary case is CTGAN and DP-CTGAN, which generally exhibit near random-guessing level membership inference risk across both datasets. However, these methods simultaneously demonstrate poor downstream utility, DE and co-expression recovery. Therefore, their apparent resistance to MIA may primarily reflect limited learning of the underlying data structure rather than privacy protection.

#### Moderate level attack performance

MVN, Embedded Diffusion and WGAN-GP occupy intermediate vulnerability regime across most attack-metric settings. However, under the calibrated GAN-leaks, which utilizes an auxiliary dataset, WGAN-GP and Embedded Diffusion exhibit substantially higher TPR@FPR values than MVN. Despite its parametric simplicity, MVN’s explicit modeling of global co-variance structure may encode sufficient distributional information to enable membership inference above random-guessing level. We can suggest that moderate vulnerability is not restricted to deep neural networks, but may also arise from faithful modeling of high-dimensional structure.

#### Elevated attack performance

CVAE and CVAE-GMM exhibit systematically elevated vulnerability across both datasets, with CVAE showing the highest risk across most attack-metric combinations. An interesting contrast is observed between CVAE and CVAE-GMM. Under GAN-leaks, which emphasizes proximity between synthetic and real samples, reconstruction- or posterior-sampling-based models are particularly susceptible. This effect is further amplified under the calibrated GAN-leaks attack in the COMBINED dataset, where auxiliary reference data increases attack power. Interestingly, CVAE-GMM, despite re-using real data to fit Gaussian mixtures, is not consistently more vulnerable than CVAE. This might indicate that data re-use might not immediately result in a higher MIA risk, as the GMM may regularize generation by sampling from cluster-level latent distributions rather than directly reflecting individual posterior encodings.

#### Distance-to-closest as a privacy proxy

Distance-to-closest real record is commonly used as a partial proxy for memorization and privacy risk in synthetic data, under the assumption that synthetic samples lying unusually close to training instances may indicate overfitting and increased membership inference vulnerability. We validated this by correlating distance-to-closest with two MIA metrics (TPR@FPR=0.1, AUC-ROC) across two attack methods in BRCA (4 comparisons per dataset) and three attack methods in COMBINED dataset (6 comparisons per dataset). When a statistically significant Spearman correlation is observed between two metrics, a trend line is shown with a shaded confidence interval, and significance is indicated by the corresponding False Discovery Rate (FDR)-adjusted p-value (i.e. q-value).

For distance-based attacks, we observe strong negative correlations between attack success and distance-to-closest metrics. In the BRCA dataset GAN-leaks shows strong association with both attack metrics (e.g. TPR@FPR, *ρ* = −0.81, *q* = 0.01). In the COMBINED dataset, GAN-leaks variants display a similar pattern with different attack and metric combinations. In the larger dataset, the uncalibrated GAN-leaks variant is negatively associated with AUC-ROC metric only (*ρ* = −0.82, *q* < 0.01) and the calibrated GAN-leaks is negatively associated with TPR@FPR (*ρ* = −0.85, *q* <0.01), consistent with these attacks’ reliance on local density estimation around training points (Fig. 7). In contrast, the classifier confidence-based RF attack doesn’t exhibit any strong correlation with distance-to-closest metric across both datasets and metrics.

**Figure 7.**
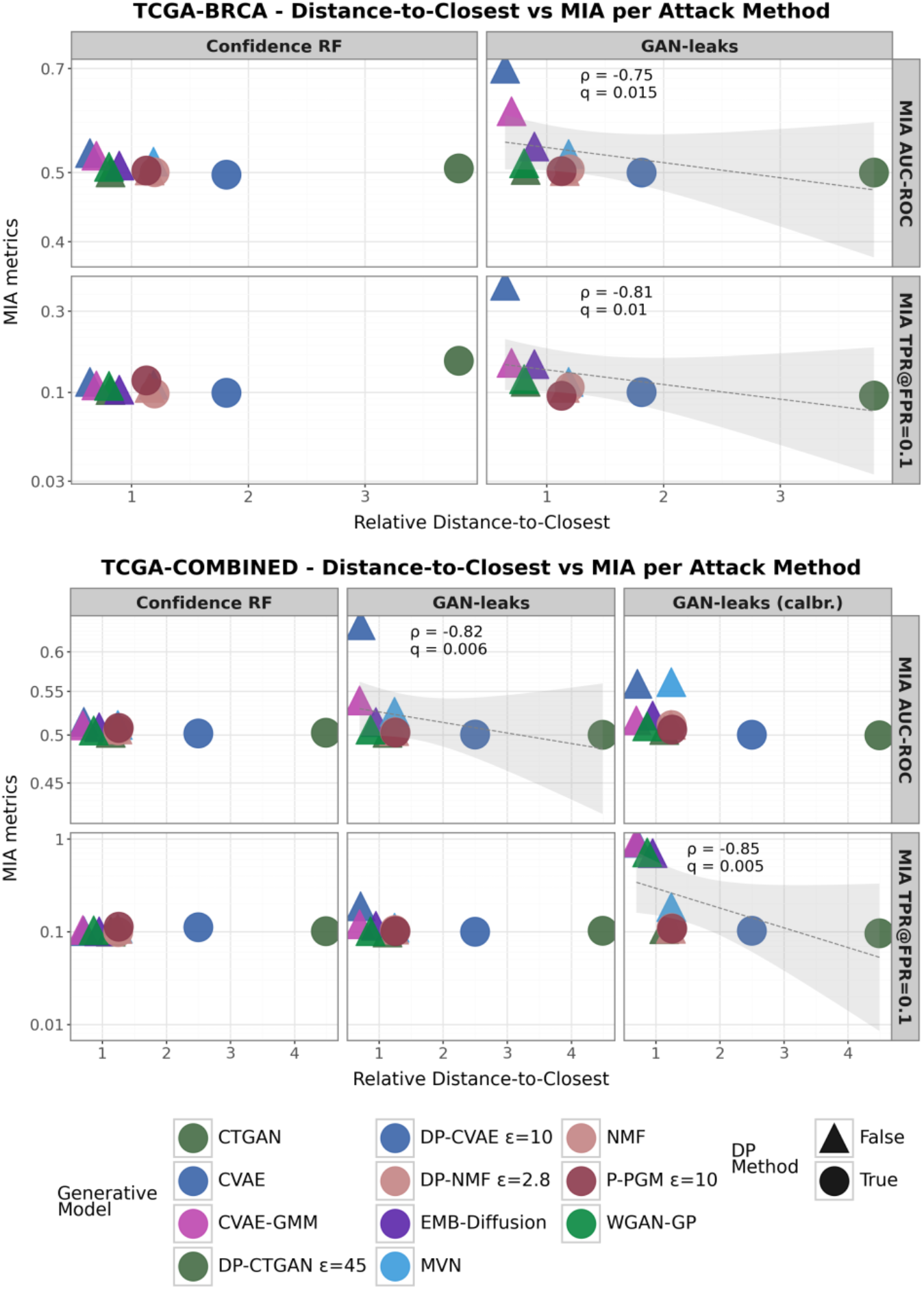
Distance-to-closest as a partial privacy proxy. On the y-axis we report two privacy risk metrics: area under the receiver operating characteristic curve (AUC-ROC) and true positive rate at a fixed false positive rate of 0.1 (TPR@FPR=0.1) and on the x-axis we report relative distance-to-closest record values, reported relative to the real data baseline (values >1 indicate synthetic samples are farther from the training set than real test samples). Differentially private (DP) and non-DP variants of the same model share the same color, but are represented with different shapes.

At the model level, DP methods and statistical approaches exhibit higher relative distance-to-closest values compared to non-DP deep-learning models. As discussed previously, these methods also show lower vulnerability under distance-based attacks. Conversely, deep learning models producing samples closer to real records are more susceptible to such attacks. This demonstrates how distance-to-closest systematically relates to both model architecture and privacy protection mechanisms.

Overall, these results indicate that distance-to-closest can serve as a useful proxy for privacy risk under certain attack types, but its predictive value is method-dependent. Recent work has also highlighted limitations of distance-to-closest alone for assessing privacy risk, noting that proximity alone may not reliably indicate meaningful privacy risk ^27^.

### Trade-offs between Evaluation Metrics

Here we investigate trade-offs between evaluation axes and how they interact. We focus on three key relationships: (i) membership inference risk against other evaluation axes, examining whether privacy-preserving methods sacrifice performance (Fig. 8); (ii) differential expression recovery against other axes, assessing whether biological plausibility aligns with other quality metrics (Fig. 9); and (iii) distributional fidelity against downstream utility, examining whether statistical similarity translates to downstream predictive performance (Fig.10).

**Figure 8.**
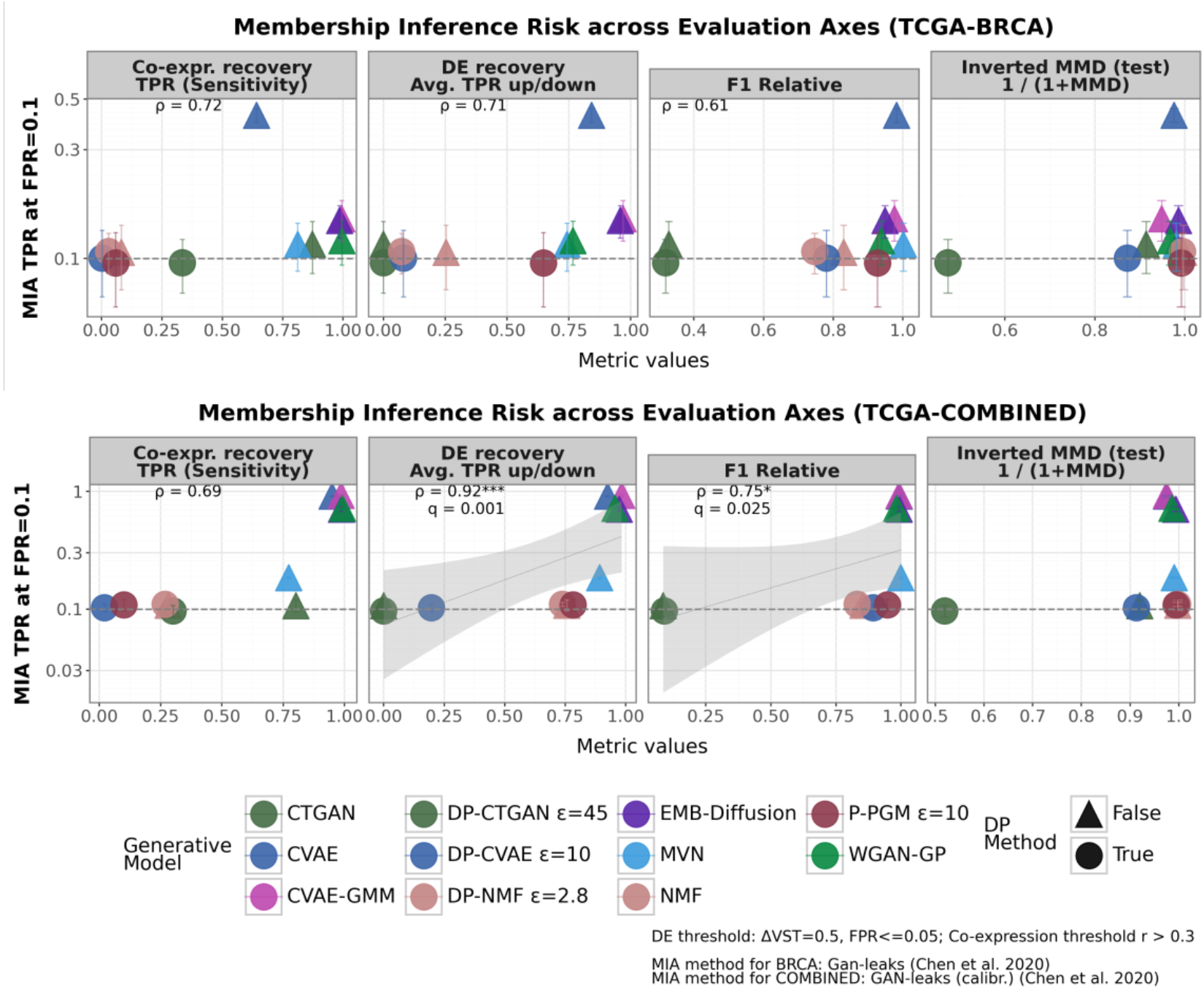
Membership inference risk across evaluation axes. Membership inference attack (MIA) risk is quantified as the true positive rate at a fixed false positive rate of 0.1 (TPR@FPR = 0.1). Each panel shows the relationship between MIA risk and a representative metric from another evaluation axis. Higher values indicate better synthetic data quality across evaluation axes.

**Figure 9.**
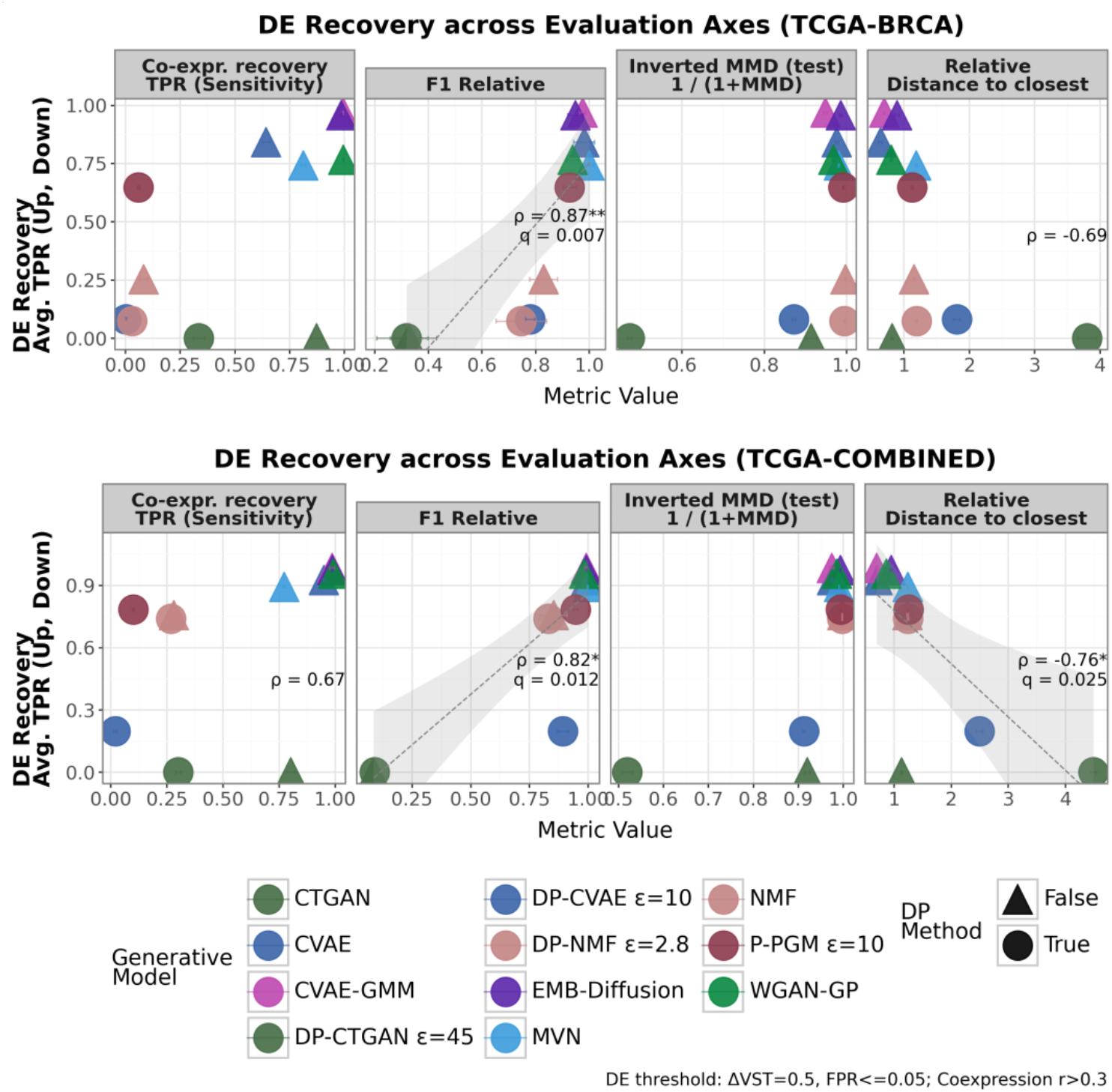
DE recovery across other evaluation axes. DE preservation is represented as the average TPR of the up- and down-regulated recovery. Each panel shows the relationship between DE recovery and a representative metric from another evaluation axis. Higher values indicate better synthetic data quality across evaluation axes except for relative distance-to-closest (i.e. values > 1 indicate synthetic samples are farther from the training set than real test samples).

For the BRCA dataset, membership inference risk is evaluated using Gan-leaks, while for the COMBINED dataset we report results using GAN-leaks calibrated. We analyze correlations separately for each dataset given differences in attack calibration and data properties (n=11 methods, Spearman correlation). Correlations presented in the following trade-off analyses are extracted from a comprehensive correlation matrix across all (raw) evaluation metrics (10 selected metrics, 45 pairwise comparisons; full matrix in Supplementary Figures S13-S14), with global FDR correction (α = 0.05) applied across all comparisons. For some of the metrics, such as MMD and co-expression false edge rate, we use their inverted versions (i.e. inverted MMD, co-expression specificity) for better readability, however, correlations are always computed on raw values. When a statistically significant Spearman correlation is observed between two metrics, a trend line is shown with a shaded confidence interval (0.95), and significance is indicated by the corresponding FDR-adjusted p-value (i.e. q-value).

#### Membership inference risk across evaluation axes

Figure 8 summarizes the relationship between membership inference risk, quantified using membership inference attack (MIA) true positive rate (TPR) at a fixed false positive rate (FPR) of 0.1 (TPR@FPR=0.1), and representative metrics from other evaluation axes: relative F1 (utility), inverted MMD (fidelity), DE recovery TPR (biological plausibility) and co-expression recovery precision (biological plausibility). Across these metrics higher scores indicate better synthetic data quality, whereas higher TPR@FPR=0.1 corresponds to increased privacy risk.

Across datasets, membership inference risk exhibits a dataset-and method-dependent relationship with utility and biological preservation, whereas no consistent association is observed with the fidelity metric MMD. In the smaller BRCA dataset, correlations between membership inference risk and other evaluation axes are generally positive (*ρ* > 0.6, for co-expression recovery, DE recovery, and F1 relative), but none reach significance after multiple-testing correction (*q* < 0.05), likely due to limited method sample size (n=11). Under Gan-leaks, DP models remain close to random-guessing level attack performance (TPR∼0.1), whereas non-DP deep models with high-utility, including CVAE variants and Embedded Diffusion, exhibit elevated membership inference risk, consistent with the observed correlation trends.

In contrast, for the COMBINED dataset under GAN-leaks calibrated, we observe significant positive associations between membership inference risk and both relative utility (*ρ =* 0.75, *q* < 0.05) and DE recovery (*ρ* = 0.92, *q* < 0.05), with co-expression recovery showing a substantial but non-significant positive correlation (*ρ* > 0.6). At the model level: deep-learning-based methods with high utility and DE recovery, including CVAE variants, WGAN-GP, and Embedded Diffusion, exhibit elevated MIA risk (TPR>0.5). MVN achieves competitive utility performance with a moderate privacy risk (TPR<0.25). The DP-model P-PGM remains close to random guessing performance (TPR∼0.1), while achieving moderate to high relative utility and DE preservation but very limited co-expression recovery.

Comparing across datasets, DP approaches such as DP-NMF and P-PGM exhibit higher DE recovery in the larger COMBINED dataset (TPR > 0.7) than BRCA, while maintaining random-guessing level attack performance (TPR ∼ 0.1) in both datasets. This observation is consistent with the prior literature suggesting that the utility loss caused by DP mechanisms decreases when the size of the data increases ^28, 29^. Notably, co-expression recovery for these models do not improve with dataset size, suggesting that while individual gene-level signals benefit from more data, network-level relationships remain difficult to capture under DP-constraints.

Together, these results suggest that the privacy-utility trade-off is present, but not universal. While deep-learning-based non-DP methods that achieve high downstream utility and biological plausibility frequently exhibit elevated MIA risk, certain approaches, such MVN and P-PGM, demonstrate that competitive utility and biological preservation can be achieved with moderate to low privacy risk. The absence of significant correlations in the smaller BRCA dataset emphasizes the importance of dataset-specific factors influencing these relationships.

#### Differential expression (DE) recovery across evaluation axes

Figure 9 examines the relationship between DE recovery and other evaluation axes. DE recovery is quantified as a true positive rate at an effect size threshold of Δ VST = 0.5 and FPR ≤ 0.05, with higher values indicating better preservation of biologically meaningful expression differences. We correlate DE TPR with representative metrics from other axes: co-expression recovery sensitivity (biological plausibility), relative F1 (utility), inverted MMD (fidelity) and relative distance-to-closest (fidelity). Except for relative distance-to-closest, higher values indicate better synthetic data quality in all other metrics.

Since DE recovery is highly consistent across both up- and downregulated genes (Supplementary Figures S13-S14, Spearman *ρ* ≤ 0.98, *q* < 0.001), subsequent analyses report DE recovery based on average TPR across both directions.

Across both datasets, DE recovery is strongly positively associated with downstream utility (*ρ* = 0.87, *q* < 0.01 in BRCA, *ρ* = 0.82, *q* < 0.05 in COMBINED), indicating that synthetic data preserving key DE genes retain the predictive signals required for the downstream task. In contrast, DE recovery shows no association with the fidelity metric MMD, while distance-to-closest exhibits a strong negative correlation in the larger COMBINED dataset (*ρ* = −0.76, *p* < 0.05), and a moderate non-significant negative correlation in the BRCA dataset. Co-expression sensitivity demonstrates a modest positive correlation with DE recovery in the COMBINED dataset, but not in the BRCA, highlighting that preserving individual DE signals does not necessarily guarantee recovery of network-level gene relationships.

Notably, both downstream tasks (i.e. molecular subtype and cancer type prediction) are primarily driven by between-class mean expression differences. Consequently, synthetic datasets that more faithfully recover DE genes retain the dominant discriminative signals required for classification, resulting in higher relative F1. These findings suggest that, for the evaluated tasks, DE preservation is predictive of downstream utility but largely independent of fidelity metrics, highlighting the need to evaluate multiple axes when benchmarking synthetic RNA-seq data.

#### Fidelity across evaluation axes

Figure 10 examines the relationship between the distributional fidelity measured by MMD (visualized as inverted), and representative metrics from utility (relative F1), biological plausibility (co-expression recovery specificity and sensitivity) and fidelity (inverted discriminative score). Co-expression specificity is defined as an inverted metric obtained through, 1/(1 + co-expression false-edge rate), where higher values indicate fewer spurious edges introduced. Inverted discriminative score reflects indistinguishability from real data under logistic regression classifier.

**Figure 10.**
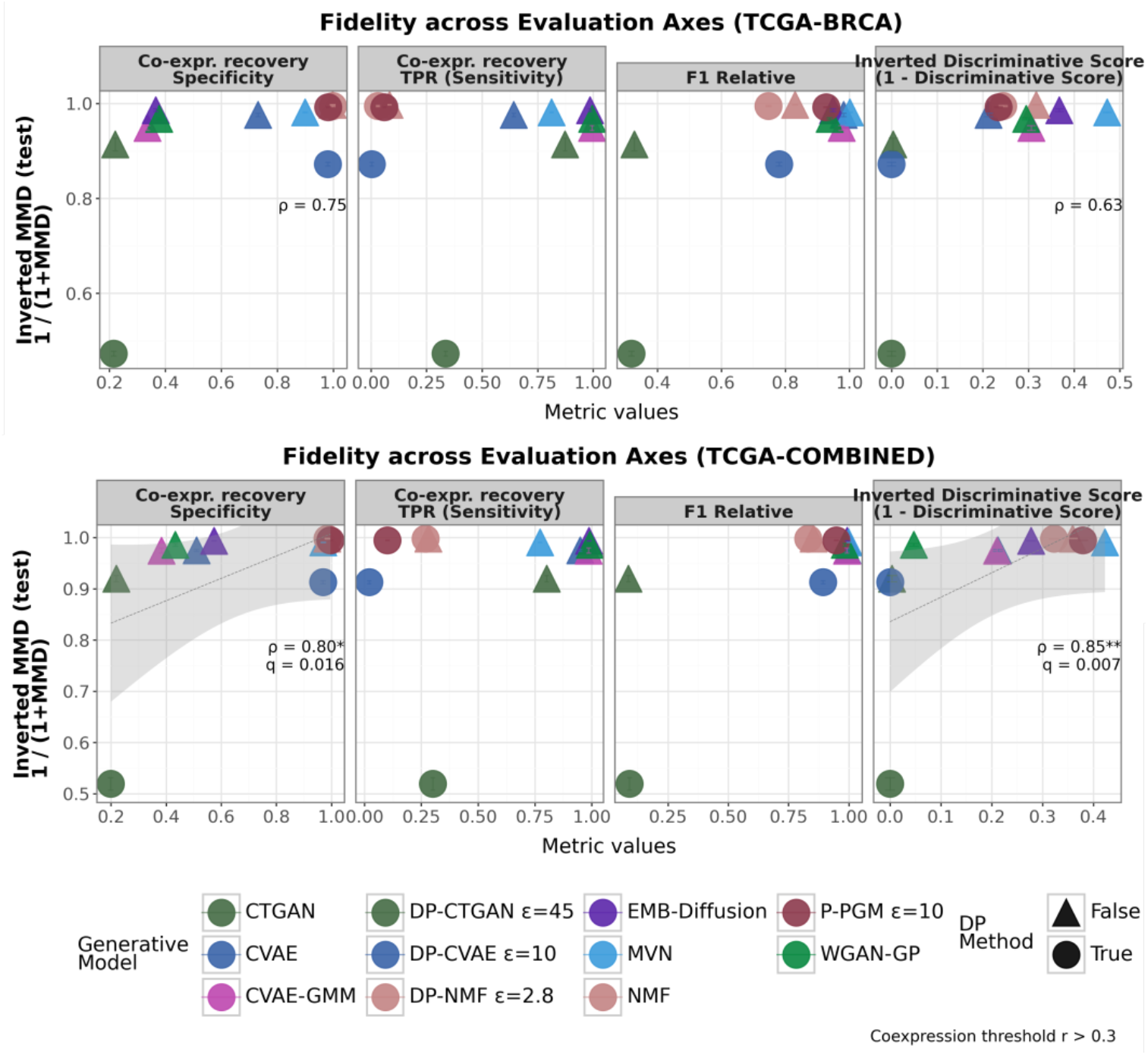
Inverted MMD (test) across evaluation axes. Each panel shows the relationship between inverted MMD (test) and a representative metric from another evaluation axis. Higher values indicate better synthetic data quality across all evaluation axes.

In the COMBINED dataset, MMD shows a strong positive correlation with co-expression specificity (*ρ* = 0.8, *q* < 0.05), and similarly a strong positive association with discriminative score (*ρ* = 0.85, *q* < 0.01). These results indicate that methods with lower MMD (better global distributional alignment) tend to introduce fewer spurious co-expression edges and are harder to distinguish from real data. In the smaller BRCA dataset, the same trends are observed but do not reach statistical significance.

At the model level: statistical approaches such as NMF and MVN, and DP model P-PGM occupy the region of low MMD and high indistinguishability, and also demonstrate higher co-expression specificity. We should however note that these methods (Fig. 10) also exhibit consistently low co-expression sensitivity, highlighting their sparse co-expression network structure. In contrast, more expressive deep generative models tend to achieve higher utility and DE recovery but exhibit elevated MMD and reduced co-expression specificity, while maintaining high co-expression sensitivity.

## Discussion

This study provides a systematic, multi-dimensional evaluation of synthetic bulk RNA-seq generators, emphasizing the tension between distributional realism, downstream utility, biological plausibility and privacy.

### Evaluation metrics are complementary, not interchangeable

Across two cancer datasets with distinct sizes and biological heterogeneity, and spanning diverse generative model families with total 11 approaches, our results indicate that no single evaluation metric provides a sufficient assessment of synthetic RNA-seq data quality. Instead, fidelity, utility and biological plausibility capture complementary, and often non-overlapping properties of generative model performance.

Utility metrics are strongly and consistently correlated with each other, reflecting a coherent notion of predictive signal preservation. Fidelity metrics (MMD, KL divergence) and discriminative score cluster together in correlation matrix, but are largely independent of utility, suggesting that global distributional similarity does not guarantee predictive performance for downstream tasks (Supplementary Figures S1-S2). These patterns indicate that fidelity and utility represent distinct evaluation axes rather than interchangeable measures.

Biological plausibility metrics further separate into distinct dimensions: DE recovery (both up- and down-regulated) aligns with downstream utility, consistent with the idea that preserving strong transcriptional signals contributes to predictive performance. In contrast, co-expression recovery metrics are largely uncorrelated with utility and DE recovery, indicating that differential expression and network structure capture different biological properties. Importantly, the observed association between DE recovery and downstream performance should be interpreted as task-dependent rather than universal. The evaluated downstream prediction tasks rely heavily on marginal mean shifts across genes; therefore, DE recovery serves as a strong proxy for predictive signal retention in this context. However, for downstream tasks that may depend more critically on gene regulation, pathway-level activity, or network topology, preservation of co-expression structure may have a stronger influence on utility than DE recovery alone.

Taken together, these findings demonstrate that evaluation outcomes are jointly shaped by generative model architecture, dataset characteristics, metric selection, and the downstream task for which the synthetic data are intended. Performance is therefore not an intrinsic property of a model alone, but emerges from the interaction between model assumptions and task-specific signal structure. Accordingly, effective benchmarking of synthetic RNA-seq generators requires a multi-dimensional evaluation framework that explicitly incorporates biological preservation alongside distributional fidelity and downstream utility. Limiting assessment to fidelity or predictive performance alone risks overlooking biologically meaningful distortions that may not immediately manifest in classifier-based metrics.

### Membership inference risk is model- and attack-dependent

The privacy evaluation reveals a clear trade-off between model expressiveness and membership risk. Models that closely approximate the underlying data distribution, particularly deep expressive generative models, tend to exhibit elevated vulnerability, whereas formally DP-constrained methods help suppress detectable membership signals. At the same time, low empirical vulnerability can also arise from limited structural learning, as underfitted generative models can exhibit reduced membership risk simply because they fail to capture meaningful data signals. Therefore, privacy risk must be interpreted in conjunction with utility and fidelity metrics to avoid mistaking underfitting as robustness.

Differences across attack algorithms, especially when auxiliary reference data are introduced, further demonstrate that measuring vulnerability depends not only on the generative model but also on the adversarial setting.

Finally, the results also show that non-DP models are not uniformly risky. Simpler statistical approaches can occupy intermediate or low-risk regimes, suggesting that they can serve as meaningful privacy-utility baselines when benchmarking more expressive architectures.

### Architecture-level patterns and failure modes across generative models

Across model families, performance differences are primarily driven by architectural inductive biases and regularization strategies, and how each model interacts with high-dimensional continuous data rather than model class alone. Variational models (CVAE and CVAE-GMM) showed strong utility and recovery of biologically meaningful signals. In particular, CVAE-GMM outperformed the standard CVAE in the smaller BRCA dataset, by coupling representation learning with density modeling via a mixture prior. The Embedded Diffusion model achieved performance comparable to CVAE-GMM across utility, biological and privacy metrics, suggesting that diffusion architectures may offer a favorable balance between expressivity and regularization.

Adversarial models showed more variable outcomes: WGAN-GP achieved a strong performance in several evaluation axes, whereas CTGAN and DP-CTGAN often underperformed, reflecting architectural assumptions tailored to mixed tabular data (e.g. feature-wise transformations and PAC-based discrimination) that might interact poorly with high dimensional continuous VST-transformed RNA-seq data. DP-CTGAN further incurred performance penalties due to privacy constraints, likely the cost of per-feature preprocessing in high dimensional settings.

Among formal DP methods, P-PGM achieved the strongest overall DP-compliant performance, however with reduced utility and biological plausibility relative to non-DP deep neural methods. This may be because the model explicitly encodes only 1-way and 2-way marginals; correlations between genes not included in these marginals are under-represented in the synthetic data, reducing biological plausibility for analyses that rely on gene-gene interactions. MVN, a simpler structure method, performed consistently well across evaluation axes, and maintained moderate privacy risk, illustrating that less expressive but well-regularized architectures can deliver robust performance.

Overall, these results show that privacy, fidelity, utility and biological plausibility are governed by architectural inductive biases: highly expressive models with regularization favor fidelity at the cost of privacy, whereas structured or regularized models trade coverage for robustness. A per-model comparison overview across selected evaluation metrics are provided in Supplementary Figure S15.

### Practical implications for users

Our benchmarking suggests that model choice should be guided by the trade-offs between utility, biological preservation and privacy. For downstream analyses requiring accurate differential expression or co-expression: CVAE-GMM and Embedded Diffusion provide strong performance, while WGAN-GP could also be useful if properly tuned. For privacy concerns, P-PGM can reduce membership inference risk but at the cost of reduced fidelity and co-expression network recovery. MVN can surprisingly offer strong utility with moderate risk and fast training, so could be a good baseline to evaluate against before training complex deep generative models. Overall, the downstream task choice, preprocessing, dataset size and heterogeneity strongly affects outcomes, therefore users should align these factors when selecting a generative model.

### Challenges and Limitations

The scope of this benchmark is limited to bulk RNA-seq data derived from cancer-relevant (TCGA) gene expression datasets with comparatively modest sample sizes (∼1,000 and ∼5,000), and a curated set of 978 landmark genes. Therefore, the observed performance of the generative models reflects their behaviour within this reduced feature space and should not be directly extrapolated to complete transcriptome setting, where higher dimensionality, sparsity, and complex gene-gene interactions may pose additional challenges. Moreover, all datasets were pre-processed using variance-stabilizing normalization (VST), which limits the generalizability of the results to models trained on raw count data or alternative normalization schemes.

The dataset size constrains the range of applicable models, as some architectures require larger cohorts for stable optimization or are unsuitable in high-dimensional settings. The relatively small number of evaluated methods (n=11) further restricts the comparative scope, particularly when examining cross-metric correlations under limited statistical power. Accordingly, the findings should be viewed as cohort-specific and method-dependent, inclusion of additional approaches may refine and modify the comparative conclusions.

Although we incorporate a broad range of fidelity, utility, biological and privacy metrics, the evaluation is necessarily incomplete. Biological preservation is assessed only through differential expression and co-expression recovery under fixed conditions, including effect size threshold, false-positive cutoff, and correlation thresholds. Alternative thresholds or biological analyses may result in different relative rankings of the generative models. Therefore, all conclusions should be interpreted as conditional on the selected metrics and evaluation settings rather than as universal statements about model performance.

Another limitation of our benchmark is that formal differential privacy (DP) mechanisms are applied heterogeneously across models, complicating their direct comparison. Although privacy budgets (*ϵ*) are reported, identical *ϵ* values do not guarantee equivalent privacy-utility trade-offs across different architectures or training schemes. Our ablation analysis of DP-CVAE across different privacy budgets (*ϵ* = 1, 2.8, 7, 10, 20, 45) illustrates the following point: while the downstream utility degrades as *ϵ* decreases, no consistent monotonic improvement in privacy protection is observed across both datasets (Supplementary Figures S16-S17). Similarly, models with stricter privacy budgets, such as P-PGM (*ϵ* = 7, 10), achieves higher relative AUC-PR utility scores compared to DP-CVAE (*ϵ* = 20, 45), and models with matched *ϵ* values (DP-NMF and DP-CVAE at *ϵ* = 2.8) exhibit different utility and attack resistance. These results emphasise that *ϵ* alone is insufficient to predict empirical privacy risk, as models may be to varying degrees more private than the budget suggests.

Finally, our privacy evaluation focuses on black-box membership inference attacks of low complexity. While these attacks can capture memorization-related vulnerabilities that are realistic in many applied settings, they do not represent worst-case adversaries. More advanced attacks such as likelihood-ratio based (e.g. LiRA ^30^), were not explored and could reveal additional privacy risks.

### Community Engagement Lessons

This benchmarking study builds on the CAMDA 2025 Health Privacy Challenge, yielding several lessons relevant to future community benchmarks in synthetic transcriptomics. First, leaderboard-style ranking is not fitted to multi-objective evaluation. No method performed consistently well across fidelity, utility, biological preservation and privacy risk, or across datasets. Encouraging accompanying extended abstracts helped surface both strengths and failure modes beyond single-score comparisons. Future benchmark efforts might consider providing example visualizations, such as heatmaps, or trade-off plots, to help participants interpret multi-metric performance and understand trade-offs better.

Second, evaluating only a single, optimized model snapshot can mask important sensitivities. Participant feedback showed that privacy and utility often vary with architectural and hyperparameter choices, suggesting that apparent model performance may reflect tuning rather than intrinsic behaviour. Benchmark platforms should encourage transparent reporting of hyperparameters, privacy budgets, and architectural variants, and allow multiple submissions to enable robust comparison across model configurations.

Finally, fragmented tooling hindered biological evaluation in participants’ extended abstracts. Analyses such as co-expression and differential expression recovery were underutilized, likely because R-based tools required specific libraries or dependencies. Future benchmarks should ensure smooth cross-language integration, for example via containerized pipelines, so participants can run all metrics without skipping analyses due to language or dependency barriers.

Overall, these observations highlight the importance of emphasizing trade-offs over rankings, encouraging exploration of model variants rather than single submissions, and providing tightly integrated, cross-domain evaluation pipelines in future benchmarks.

### Future Direction

Building on our joint effort with DECIDER ^31^ and the DREAM challenges ^32^, we are currently assessing the generalizability of the methods evaluated in this benchmark by generating synthetic ovarian cancer data and validating its quality on the ovarian cancer patients in the DECIDER cohort. This replication will be part of the OpenEBench platform (https://openebench.bsc.es/), explicitly improved to support benchmarking. Following steps should be testing the models across even larger and more heterogeneous datasets, including single-cell RNA-seq, which was also one of the tracks of the CAMDA 2025 Health Privacy Challenge, as well as multi-omics and non-cancer electronic health records. In parallel, development of generative models for data synthesis could benefit from multi-objective optimization, explicitly balancing utility, biological plausibility and privacy rather than treating these as post-hoc evaluations.

Synthetic data utility and privacy may vary across demographic or clinical subpopulations, but our analyses are reported only at the dataset-level. Prior work has shown disparate impacts of differential privacy ^27^ and membership inference attacks ^33^. Future benchmarking should assess synthetic data generation under sensitive attribute stratification (e.g. race, gender) to enable fairness-aware evaluation and more realistic assessment.

Finally, privacy and biological plausibility evaluation could be expanded. Privacy assessment should extend beyond membership inference to include re-identification, linkage, attribute inference, or model inversion attacks ^34, 35, 36,37^. Differential privacy mechanisms should be systematically studied to disentangle the effects of privacy accounting, noise injection strategies and model architecture on empirical trade-offs. Biological plausibility evaluations could further be extended to incorporate gene-set enrichment and functional pathway enrichment analyses. Collectively, these directions can enhance the robustness, fairness, and interoperability of synthetic data benchmarking in transcriptomics.

## Conclusions

In this work, we benchmarked generative models for private and biologically meaningful synthetic data generation, building on the community competition, CAMDA 2025 Health Privacy Challenge. Our results demonstrate that, within transcriptomics context, the quality of synthetic data must be evaluated across multiple complementary axes, as no single metric can capture biological signal recovery and utility. These findings highlight the importance of multi-dimensional benchmarking frameworks for synthetic data generation, as evaluation outcomes are jointly shaped by generative model architecture, dataset characteristics, metric selection, and the downstream task for which the synthetic data are intended. Consequently, this evaluation framework serves as a guidance for transcriptomics and broader healthcare applications and emphasises that task-specific benchmarking should become a standard practice before data sharing.

## Methods

### Challenge aim and design

The Health Privacy Challenge CAMDA 2025 (https://benchmarks.elsa-ai.eu/?ch=4) aimed to benchmark generative models for private and biologically plausible synthetic transcriptomics data generation. The challenge was established with two tracks: Track 1 focused on bulk RNA-seq data and Track 2 focused on single-cell RNA-seq data. Track 1 ran in two complementary tasks that are referred to as Blue Team versus Red Team, sharing the bulk RNA-seq dataset but diverging in goals.

- The Blue Teams generated synthetic bulk RNA-seq datasets that balance the utility and privacy trade-off.
- The Red Teams designed methods to launch a membership inference attack (MIA) on synthetic datasets to determine whether the target sample is included in the training dataset that generated the synthetic dataset.

In this benchmarking study, we only cover Track 1 Blue Team baseline methods and participant submissions, and only use the baseline methods for membership inference attack-based privacy risk assessment and do not include Red Team participant submissions. All baseline generative and MIA methods, and evaluation metrics are available for reproduction at https://github.com/PMBio/Health-Privacy-Challenge following the challenge format (See Supplementary Material for details).

### Datasets and Pre-processing

We re-distributed the open access The Cancer Genome Atlas (TCGA) ^38^ bulk gene-expression data (STAR counts) obtained from the Genomic Data Commons (GDC) portal (https://portal.gdc.cancer.gov) using the TCGABiolinks R package ^39^. We provided two pre-processed datasets, that we refer to as TCGA-BRCA and TCGA COMBINED.

#### TCGA-BRCA

The dataset contains 1094 (donor) samples with known subtypes from breast cancer (TCGA-BRCA) cohort, belonging to either LumA, LumB, Basal, Her2, or Normal.

#### TCGA COMBINED

The dataset consists of 5222 primary tumour (donor) samples containing 12 cancer types spanning 10 tissue types: Breast (TCGA-BRCA), Colorectal (TCGA-COAD), Esophagus (TCGA-ESCA), Kidney (TCGA-KIRC, TCGA-KIRP), Liver (TCGA-LIHC), Lung (TCGA-LUSC, TCGA-LUAD), Ovarian (TCGA-OV), Pancreatic (TCGA-PAAD), Prostate (TCGA-PRAD), and Skin (TCGA-SKCM). The raw count data for both datasets is pre-processed similar to ^17^ :

- Each donor contributed a single sample.
- For TCGA-BRCA, the samples without known molecular (sub)types are filtered.
- Low count genes are filtered: genes with counts ≥ 1 in at least 10% of samples are kept.
- Remaining genes are normalized using the DeSeq2 variance-stabilizing transformation (VST) transformation with blind=FALSE ^40^.
- Analysis was restricted to the LINCS L1000 landmark gene set comprising 978 genes ^41^ that are reported to capture most transcriptional variance and enable the inference of the remaining genes in transcriptome (https://clue.io/command?q=/gene-space%20lm, accessed on Nov 1, 2024).

TCGA COMBINED dataset was subsequently divided into two stratified sets, comprising 4398 and 824 donors/samples. The smaller set (824) was reserved for launching membership inference attack algorithms that require an external dataset. Consequently, only the larger dataset was employed to generate and evaluate synthetic data.

All generative models were trained using VST-transformed real expression data and generated synthetic datasets directly in the VST-transformed expression space, restricted to the 978 LINCS L1000 landmark genes.

### Experiment Design

TCGA bulk RNA-seq datasets, denoted as *D*, were used to generate synthetic datasets using a generative model *f*(*D*). Each dataset was split into a stratified five-fold cross validation setting using a fixed random seed (42) to enable comparisons across models. For each fold *i* ∈ {1,2,3,4,5}, a shape-matched synthetic dataset *D*_*SYN*_^(*i*)^ was generated from the model *f*(*D*) trained on the corresponding training partition *D*_*TR*_^(*i*)^ . The quality of the synthetic data was evaluated on the corresponding held-out test set *D*_*TE*_^(*i*)^ and performance metrics were averaged across the five folds to obtain a robust estimate of generalization. This cross-validation procedure also produced five membership inference datasets, one per fold, in which members correspond to training samples and non-members correspond to test samples.

### Differential Privacy (DP)

DP is a mathematical definition of privacy, which roughly indicates that including or removing a sample from the dataset does not significantly affect the outcome of an analysis ^1^. A method *M* is (*ϵ,δ*)-differentially private if, given two datasets of *D*_1_ and *D*_2_ that differ by at most one element, it holds: *P*[*M*(*D*_1_) ∈ *S*] ≤ *e*^*ϵ*^ *P*[*M*(*D*_2_) ∈ *S*] + *δ* where *S* is a subset of outputs generated by *M, ϵ* is a non-negative parameter that controls the “privacy budget” and *δ* is a non-negative parameter that indicates the level of privacy guarantee being violated. Determining *ϵ* and *δ* involves a trade-off between utility and privacy. For instance, smaller *ϵ* indicates stronger privacy, allowing more noise, and therefore might lead to less accurate results in utility. DP can be layered onto several paradigms (factorization, PGMs, VAEs/GANs), typically by perturbing gradients, statistics, or outputs. The benefit is formal privacy guarantees; the cost is reduced fidelity as ε shrinks.

### Generative Models

Here we describe the methods included in this benchmarking study in model family groups. Although different taxonomical approaches for generative models are used ^42^, here we group models according to their underlying generative mechanisms (statistical baselines, latent-variable autoencoders, adversarial models, etc.) and focus on the subset of approaches that were instantiated and evaluated in this benchmark.

Baselines were curated by the benchmark organizers to provide reference performance for each model family and were publicly available during the challenge. These include Multivariate Normal Distribution (MVN), Conditional Variational Autoencoder (CVAE) ^43,44^, and Conditional Tabular Generative Adversarial Network (CTGAN) ^45^, where CVAE and CTGAN were implemented with and without Differential Privacy (DP) ^1^. In addition, two post-challenge methods, CVAE with Gaussian Mixture Model (CVAE-GMM) ^46^ and Wasserstein GAN with Gradient Penalty (WGAN-GP) ^47, 48^, were included to increase the diversity of evaluated methods.

Submissions were provided by external participants and reflected realistic generative modeling practices. These include a Diffusion Model with noise injection ^49^, Private-PGM (P-PGM) ^50^, and implementation of Non-negative Matrix Factorization (NMF) ^51^ with and without DP.

#### Statistical and sampling based

These models specify a parametric probability distribution and sample from it, assuming relatively simple dependence structures and provide interpretable baselines, highlight the performance of classical probabilistic assumptions, and serve as a reference point for more complex generative models. Here we detail the instances evaluated in this benchmark.

##### 1. Multivariate Normal (MVN) Model

The Multivariate Normal (MVN) model assumes a joint Gaussian distribution, where the mean vector and covariance matrix capture pairwise correlations among genes. In this benchmark, we adopted a class-conditional MVN, fitting separate models for each subtype or disease type to serve as a simple and interpretable statistical baseline.

##### Implementation details

For each subtype(/disease type) *s*, the empirical mean vector 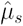 and covariance matrix 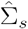 were computed from training data. To ensure numerical stability, the covariance was symmetrized and projected to a positive semi-definite matrix by clipping negative eigenvalues and adding a small diagonal jitter *ϵI*. Synthetic samples were then drawn as 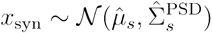 and perturbed with additive multivariate Gaussian noise

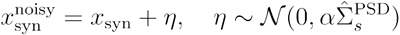 where *α* = 0.5 controls the noise scale. This additive noise simulates potential real-world measurement variability while preserving the underlying correlation structure of the data. We investigated noise levels of 0.5, 1, and 5 to systematically explore the effect of increased variance on downstream analyses, and reported the resulting impact on differential expression recovery in Supplementary Figure S18.

##### Limitations

The MVN assumption restricts the model to Gaussian features and linear correlations, which may not capture multimodality, over-dispersion, or nonlinear dependencies in gene expression.

##### 2. Non-Negative Matrix Factorization (NMF)

The NMF model represents data as low-rank latent factors^51^ and is widely used in bioinformatics to decompose datasets into interpretable structures, such as metagenes and sample clusters to provide biological insights ^52^. This model was submitted by a participant of the challenge.

##### Implementation

Normalized counts were filtered to obtain non-negative integer matrices. The input expression matrix 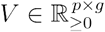 (patients × genes) is factorized into non-negative matrices 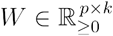 and 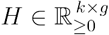 using MiniBatchNMF from Scikit-learn, which employs a coordinate descent solver and computes updates in batches for scalability:*V* ≈ *WH* .

The NMF *k* = 50 rank was determined via Frobenius-norm reconstruction error elbow criterion. Latent encodings in *W* capture metagene activity and are clustered via K-means to form biologically coherent subgroups. The number of clusters *k*_*c*_ was dynamically set equal to the number of classes in input data.

For each cluster *c*, per gene statistics were computed, mean (μ_*c*_), variance 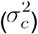, and zero-fraction (*z*_*c*_), to parameterize a Gaussian sampler that reconstructs synthetic count profiles while preserving realistic dispersion and sparsity patterns.

To preserve class identity, a Random Forest classifier (*n*_tree_ = 100) was trained on the real latent encodings *W*, using Gini impurity to measure the quality of a split. Synthetic labels were assigned probabilistically by sampling from the posterior class probabilities predicted by this classifier.

##### Limitations

The linear additive structure of NMF may miss complex, nonlinear dependencies between genes or samples.

##### 3. Differentially-Private Non-Negative Matrix Factorization (DP-NMF)

DP-NMF follows the same modeling framework as NMF approach but augments it with formal differential-privacy (DP) mechanisms to protect individuals contributing to the synthetic data. The method combines matrix factorization with calibrated noise injection such that the released synthetic gene gene-expression profiles satisfy an (*ϵ δ*)-DP guarantee while retaining the underlying biological patterns.

##### Implementation

The model first decomposes the non-negative matrix 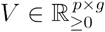 (patients *x* genes) into latent factors,*V* ≈ *WH* where 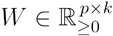 and 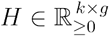 are estimated using MiniBatchNMF. Privacy protection is achieved through three-stage noise injection pipeline:

1. Output perturbation of the learned basis *H*: Gaussian noise is added to the learned components,
2. Privacy-aware clustering: K-means is applied to the latent encodings *W*, and cluster centroids are sanitized via Laplace noise to ensure DP over cluster assignments and
3. Sanitization of summary statistics: For each cluster *c*, the summary parameters used to generate synthetic samples,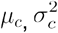, and *z*_*c*_, are released under DP using Laplace perturbation.

Privacy budgets were allocated across the pipeline as: *ϵ*_*NMF*_ = 0.5, *ϵ*_*clust*_ = 2.1, and *ϵ*._sum_ = 0.2. Under standard composition, the model therefore operates with a total privacy budget of *ϵ*_total_ = 2.8, for a fixed *γ*, which was held constant across all experiments. Synthetic samples are then drawn from the privatized summaries on a per-cluster, per-gene basis using the same sampling strategy as in the non-private NMF model.

##### Limitations

In addition to the inherent linearity constraints of NMF, DP noise introduces a fidelity– privacy trade-off: performance is sensitive to the choice of rank *k*, noise scales, and the allocation of *ϵ* across stages. The current implementation relies on manually chosen, scenario-specific privacy budgets; applying this method to new datasets would require justification of these settings and a unified privacy-accounting framework to ensure robustness and reproducibility.

### Probabilistic Graphical Models (PGM)

These methods learn an explicit probabilistic model over the variables using a dependency graph (e.g., Bayesian networks or Markov random fields). The joint distribution factorizes into local conditional or potential functions defined by the graph structure, and synthetic samples are generated by sampling from this fitted distribution ^53^. We evaluated Differentially Private PGM as an instance of this family, which was submitted by the participants.

#### 1. Differentially Private Probabilistic Graph Model (DP-PGM)

Private-PGM ^50^ (P-PGM) is a marginals-based differentially private synthetic data generator and uses the implementation provided in a public repository (https://github.com/ryan112358/mbi). The method operates by first selecting a collection of low-dimensional marginals over the dataset, releasing them under differential privacy, and then fitting a graphical model that is consistent with the noisy marginals.

##### Implementation

For a given dataset, (i) 1-way marginals for individual genes and (ii) 2-way marginals between each gene and class label were computed, yielding summary statistics that preserve feature-label dependencies. Each marginal *M*_*l*_ was perturbed using a DP mechanism to produce 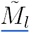, and a PGM was then estimated such that its implied marginals match { *M*_*l*_ }. The resulting joint distribution factorizes over model cliques, and synthetic samples are obtained by sampling from this distribution.

Continuous gene-expression values were discretized using quantile binning into four equiprobable bins prior to marginal estimation. Model parameters were optimized using the default mirror-descent routine provided in the public repository. Two privacy-budget settings were evaluated (ϵ = 7 and ϵ = 10); all other hyperparameters were kept fixed.

##### Limitations

Fidelity depends on the chosen marginal set and discretization scheme; higher-order (involving three or more genes) or nonlinear gene-gene dependencies not captured by the factorization may be under-represented. In high-dimensional continuous settings, scalability and structural misspecification can limit biological correlation recovery.

### Latent-variable Autoencoders

The variational autoencoder (VAE) is a latent-variable generative model that consists of encoder-decoder neural architecture with a parametric latent prior, such as isotropic Gaussian ^44^. The encoder maps each observation *x* to a distribution over latent variables *z*, while the decoder parametrizes the conditional likelihood *p* (*x*|*z*), enabling generation from the prior and decoding. VAEs are widely used to obtain low-dimensional, denoised representations that capture dominant sources of variation in RNA-seq data and related omics modalities. Following instances of the VAEs are evaluated in this benchmark:

#### 1. Conditional variational autoencoder (CVAE)

Conditional VAEs (CVAEs) augment the latent representation with sample-level covariates, allowing the generative process to respect experimental or biological attributes ^43^.

##### Implementation

The evaluated CVAE is implemented in Pytorch (2.4.0) ^54^ and employs a fully connected encoder-decoder architecture with three hidden layers per branch and ReLU activations. Conditioning variables (e.g., disease subtype/type) are incorporated either as one-hot vectors or via an embedding layer and are concatenated with both the encoder features and the latent representation in the decoder pathway. The latent space follows a Gaussian prior, and model training maximizes a \beta-weighted ELBO with mean-squared-error reconstruction loss and a KL-divergence regularizer, using the form *L* = MSE + *β*KL, with *β* = 0.001. The comparative results for one-hot encoding and embedding layer-based representation are included in the Supplementary Tables S4-S5.

##### Limitations

Like other autoencoder-based generators, CVAEs can exhibit reconstruction bias toward the training manifold, which may limit exploration of rare expression modes. Performance is sensitive to preprocessing choices (e.g. normalization) and to the relative weighting of the KL term, where excessive regularization may yield latent-space collapse while insufficient regularization can degrade sampling quality. Conditioning effectiveness depends on the representation of covariates (one-hot versus learned embeddings), and poorly tuned conditioning can lead to label leakage or reduced separation between biologically meaningful groups in the latent space.

#### 2. Conditional variational autoencoder with Gaussian Mixture Models (CVAE-GMM)

The CVAE–GMM extends the baseline CVAE by replacing the standard isotropic Gaussian prior with a Gaussian mixture model (GMM) ^55^ in latent space, following the intuition of ^46^. The objective is to better capture latent heterogeneity within a conditioning label, such as disease groups that may contain unobserved subtypes, by allowing multiple latent components rather than a single unimodal prior.

##### Implementation

The neural architecture and the training procedure are identical to the baseline CVAE. After training, the encoder is applied to the full (real) dataset to obtain posterior mean latent representations *z*. For each disease type or subtype label, relevant latent factors are subsetted and a GMM with *n*_components_ = 3 is fitted and stored for that group (in both benchmark datasets). During generation, latent samples are drawn from the corresponding fitted GMM rather than a standard normal distribution, and these samples are decoded using the trained CVAE decoder. Essentially, GMM acts as a dynamic data-adaptive mixture prior that replaces the single Gaussian prior used in the baseline CVAE.

##### Limitations

This approach introduces modeling complexity and requires careful selection of the number of mixture components, which may affect stability. Because GMM is fitted on the latent encodings derived from the original dataset, the method involves additional post-hoc pass over the data; which may increase the risk of latent overfitting and memorization, and therefore increase privacy vulnerabilities.

#### 3. Differentially Private Conditional Variational Autoencoder (DP-CVAE)

The DP-CVAE extends the baseline CVAE by incorporating a formal differential-privacy (DP) mechanism into training. The model architecture, encoder-decoder structure, and conditioning on the disease labels are identical to baseline CVAE, but per-sample gradients are clipped and perturbed with Gaussian noise to ensure privacy, providing (*ϵ, δ*) guarantees for the learned parameters.

##### Implementation

DP training was incorporated using Python package Opacus (v1.5.2) ^56^ which implements DP-SGD algorithm ^2^ in Pytorch. A privacy engine wraps the model, optimizer and data loader, and operates as follows:

1. Per-sample gradients are computed and clipped to a maximum *L*_2_ norm of 0.1.
2. Gaussian noise is added to the clipped gradients, with target privacy budget *ϵ*, target *δ* = 10^−5^, and batch size set to 64.
3. Privacy accounting is performed to track cumulative *ϵ* over all optimization steps.

To assess the privacy-utility trade-off we swept across *ϵ* = 1, 2.8, 7, 10, 20, 45 (Supplementary Figures S16-S17). Some of the *ϵ* values (*ϵ* = 2.8, 7, 10) were chosen to match privacy budgets submitted by challenge participants.

##### Limitation

In addition to the baseline CVAE limitations, DP-CVAE introduces an inherent tradeoff between privacy and fidelity: stricter privacy budgets reduce reconstruction quality, latent-space separation, and preservation of biologically meaningful variation, while looser budgets improve utility but relax privacy guarantees. Model performance is sensitive to the choice of gradient clipping norm, noise scale, and batch size.

### Generative Adversarial Networks (GANs)

The generative adversarial networks are composed of two components, a generative model *G* that captures the data distribution, and a discriminative model *D* that estimates the probability that a sample came from the training data rather than *G*, both of which are trained simultaneously ^57^. Training optimizes a min–max objective, effectively matching the generated distribution to the data distribution without requiring an explicit likelihood. Conditional variants allow generation to be guided by covariates (e.g., disease subtype), enabling targeted synthetic data creation. We evaluated the following instances in this benchmark.

#### 1. Wasserstein GAN with Gradient Penalty (WGAN-GP)

In Wasserstein GAN, the adversarial training follows the Wasserstein objective with gradient penalty to enforce a 1-Lipschitz constraint ^47^. WGAN-GP replaces the standard GAN objective with a Wasserstein loss and gradient penalty to improve stability and global distribution matching ^48^.

##### Implementation

A conditional WGAN-GP where both the generator and discriminator are fully connected networks conditioned on disease subtype is trained. The generator maps latent Gaussian noise concatenated with a disease-condition vector to the expression feature space, using a multilayer feed-forward architecture with batch normalization, ReLU activations, and dropout. Gradient penalty set equal to λ = 10 to enforce a 1-Lipschitz constraint, and for each generator update, the discriminator (critic) is updated *n*_critic_ = 5 times, computing the loss using random linear interpolations between real and synthetic samples. The generator is optimized to maximize the critic’s score on synthetic samples. Both networks are trained with Adam. Inputs are z-score normalized prior to training, and generated samples are inverse-transformed back to the original expression scale at synthesis time. Similar to CVAE, we investigated both one-hot disease encodings, and a learned embedding layer for compact condition representations (Supplementary Tables S4-S5).

##### Limitations

As with other GAN-based models, WGAN-GP may under-represent minority modes and is sensitive to critic-generator update balance and learning-rate tuning.

#### 2. Conditional Tabular GAN (CTGAN)

CTGAN is a tabular focused GAN model with conditional sampling which handles imbalanced categories and mixed data types ^45^.

##### Implementation

The model uses a fully connected generator with stacked residual blocks and batch normalization, which maps a latent Gaussian input concatenated with a conditional vector to the transformed feature space. The discriminator operates on PAC-grouped samples (pac = 10), enforcing a 1-Lipschitz constraint via a Wasserstein loss with gradient penalty (WGAN-GP). Conditioning is implemented by concatenating the one-hot-encoded label vector to both the generator and discriminator inputs, supporting label-guided sample synthesis.

Training follows the reference CTGAN implementation, using mode-specific data normalization. The generator is optimized with Adam, and the model is trained 10,000 iterations with a batch size of 64. Synthetic samples are generated by sampling latent noise and conditioning vectors. Synthetic samples are produced by sampling latent noise and conditioning vectors at generation time.

##### Limitations

CTGAN inherits known GAN trade-offs, including mode under-coverage though it attempts to solve this via the mode-specific normalisation. Performance is sensitive to the balance between discriminator and generator learning rates. Preprocessing may distort marginal RNA-seq feature structure, and adversarial optimization remains less stable than likelihood-based approaches.

#### 3. Differentially Private Conditional Tabular GAN (DP-CTGAN)

DP-CTGAN extends the baseline CTGAN implementation by incorporating differential privacy.

##### Implementation

DP training was implemented using the Python package Opacus (v1.5.2) which implements the DP-SGD algorithm in PyTorch and SmartNoise-Synth (https://docs.smartnoise.org/). Per-sample gradients are *l*2-clipped and perturbed with Gaussian noise; the generator is trained normally using the discriminator’s privatized outputs. Privacy budgets are accounted for using the moments accountant, with target *δ* = 10^−5^, batch size set to 64 and per-sample gradient clipping set to 0.1. A preprocessor epsilon is also applied to compute bounds for each feature, scaled by preprocessor-eps-multiplier = 0.1. For high-dimensional data (978 features), this preprocessor step consumes a significant portion of the privacy budget, which limits the minimum achievable epsilon. In our experiments, training with ϵ < 45 was unstable or failed, whereas *ϵ* = 45 successfully ran, reflecting the combined cost of generator training and preprocessing.

##### Limitations

In addition to baseline CTGAN limitations, differential privacy introduces a strict privacy-utility trade-off, where high-dimensional data amplifies privacy cost. The preprocessor’s contribution to epsilon reduces flexibility in achieving stricter privacy guarantees. As with most DP-GAN approaches, only the discriminator is privatized; the generator receives gradients through the privatized discriminator, so overall privacy guarantees rely on correct accounting of DP-SGD in the discriminator.

### Diffusion Models

A diffusion model consists of two key phases: a forward diffusion stage, where input data is gradually perturbed with Gaussian noise over multiple steps, and a reverse diffusion stage, in which the model systematically reconstructs the original data by reversing this process. Originally introduced in 2015 ^58^, and later popularized through denoising diffusion probabilistic models ^59^, and score-based generative models ^60^, diffusion models have shown strong performance across high-dimensional and complex data, including images, text, and tabular data ^42^. In this benchmark, we evaluated the following instance of a diffusion model:

#### 1. Embedded Noisy Diffusion

##### Implementation

The core strategy of the Embedded Noisy Diffusion is to use a conditional diffusion model with DP-SGD ^49^. Timesteps are encoded via sinusoidal positional embeddings and class labels via neural embeddings, combined into a single conditioning vector. The network comprises ResLinear residual blocks with Group Normalisation (8 groups), SiLU activations, dropout (0.2), and a progressive dimensionality scheme for embedding bottlenecks. Differential privacy is applied through DP-SGD with gradient clipping (max-grad-norm = 1.0) and a minimal noise multiplier (dp-noise-multiplier = 0.00001), chosen to preserve accuracy while providing privacy guidance. SMOTE oversampling ^61^ is adopted to balance class representation, and features are scaled with a QuantileTransformer and inversely transformed to maintain biological interpretability. The denoising objective is optimized with MSE loss, AdamW, a OneCycleLR scheduler, and early stopping. The model was trained on a single NVIDIA H100 GPU producing ∼25M parameters.

##### Limitations

While DP-SGD is applied, the extremely low noise multiplier (0.00001) means the model cannot provide a formal privacy guarantee, therefore this model is not considered as a DP model in this benchmark. Additionally, the model requires careful tuning of ResLinear blocks and normalization for stability, and training remains computationally intensive despite inference being efficient.

### Evaluation Framework

All models were evaluated under identical data splits, random seed, and preprocessing pipelines to ensure comparability. All synthetic datasets were generated with identical sample size and feature dimensionality as the training dataset (per each fold).

We should note that some of the methods in the benchmark incorporate Differential Privacy (DP) and direct cross-model comparison at a single ϵ was not always feasible, while submitted models were included in the benchmark as provided by external participants and their privacy budgets were determined by the submitters and could not be harmonized. Instead, we (i) evaluate each model at the ϵ provided by the submitters or required for stable training, and (ii) where possible, include matched-ϵ comparisons across models. Therefore, we interpret results primarily in terms of privacy–utility trade-off profiles rather than absolute rankings.

### Evaluation Metrics

We evaluate models along four complementary dimensions: (i) statistical fidelity to the real data distribution, (ii) downstream task utility, (iii) biological plausibility of generated features, and (iv) privacy risk. Supplementary Table 1 presents the complete set of metrics that are used in evaluation.

### Fidelity

Fidelity evaluates how closely the synthetic data matches the statistical properties of the real data. We quantify distributional similarity using Maximum Mean Discrepancy (MMD) and gene-wise Kullback– Leibler (KL) divergence.

MMD was computed using an RBF kernel with bandwidth parameter *γ* = 1/*d* where *d* denotes the number of features, and is evaluated between synthetic samples and the real training set (MMD-train), as well as between synthetic samples and the real test set (MMD-test). Lower MMD (close to 0) indicates closer alignment between real and synthetic distributions.

KL divergence was computed across marginal (gene-wise) feature distributions in the direction KL (*P*_real_ ‖*P*_syn_), treating the real data distribution as reference. Marginal densities were estimated using fixed-width histogram binning with shared bin edges and ϵ-smoothing (10^−10^) to avoid numerical instability. KL values were averaged across genes. We report synthetic-vs-real-train (authenticity) and synthetic-vs-real-test (generalizability) values. Lower KL divergence (close to 0) reflects better preservation of feature-level statistical structure. To provide better comparison across models in visual summaries (Fig.1), MMD and KL metrics were displayed after inverted transformation (e.g. Inverted KL = 1 / (1 + KL) . This way, the methods with the smallest values, and thus better similarity to real data are indicated with higher bars.

We compute the distance-to-closest record statistic between synthetic and real (train) samples as a measure of local coverage of the real data and to identify potential oversampling or sample memorization. Smaller distances indicate closer alignment between synthetic and real samples. The distance-to-closest metric was reported relative to the base value computed as the distance-to-closest within real train and real test data. While distance-to-closest is informative for fidelity, it is also commonly interpreted in the context of privacy; we therefore revisit this metric in the privacy evaluation section.

Finally, to assess whether synthetic samples are statistically distinguishable from real data, we compute a discriminative score using a logistic regression classifier trained to separate real training samples from synthetic samples. The combined dataset (real train + synthetic) is randomly split into training and test partitions. After training, evaluation is performed on an extended test set that includes (i) the held-out portion of the mixed real/synthetic data and (ii) independent real test samples not used during classifier training. Performance is quantified using the F1 score and referred to as the *discriminative score*, where higher values indicate easier separation between real and synthetic samples. We define inverted discriminative score as 1 − discriminative score, such that higher values correspond to greater similarity between real and synthetic data and thus higher fidelity.

### Utility

We evaluate downstream task utility by measuring how well models trained on synthetic data generalize to real data in relevant predictive settings. For TCGA-BRCA, the downstream task is molecular subtype prediction; for TCGA-COMBINED, the downstream task is multi-class cancer type prediction across multiple tissues. Throughout the evaluation, “downstream applications” refer specifically to these tasks, which also serve as proxies for biological utility at different levels of granularity.

We train a one-vs-rest Logistic Regression classifier on the synthetic training set and evaluate its performance on the held-out real test set (train-on-synthetic, test-on-real, TSTR). As a reference, we train the same classifier on the real training set and evaluate it on the same real test set (train-on-real, test-on-real, TRTR). Evaluation is performed under five-fold cross-validation, and mean performance across folds is reported. To account for class imbalance, we report multiple metrics, including accuracy, macro- and class-weighted precision-recall AUC, F1 score, and AUROC. Alignment between synthetic-trained and real-trained performance indicates preservation of task-relevant structure. For interpretability, we additionally report relative scores (e.g., AUROC relative, F1 relative), defined as the performance of the models trained on synthetic data normalized by the performance achieved when trained on real data (perf_synthetic_/perf_real_). Relative values closer to one indicate stronger downstream utility of the synthetic data.

To assess whether the synthetic data preserves feature-level signals, we additionally compute the count and percentage (%) of overlapping predictive genes identified in the synthetic-trained model that are also among the top predictive genes in the real-trained model. For each class, we extract the top-10 features corresponding to the largest positive Logistic Regression coefficients from the one-vs-rest classifiers, aggregate these per-class feature sets, and compare their overlap between synthetic-trained and real-trained models. Higher overlap indicates stronger retention of biologically and task-relevant structure in the synthetic data.

### Biological Plausibility

Unlike other domains where synthetic data generation is applied, healthcare data, especially high-dimensional genomics data, requires additional evaluation metrics to ensure validity and reliability. For bulk RNA-seq datasets, assessing biological plausibility is critical, including the ability of synthetic data to reproduce meaningful differential expression patterns (e.g., between molecular subtypes or cancer types) and realistic gene co-expression structures. We followed a biological utility analysis workflow similar to^17^, using their provided codebase (https://github.com/MarieOestreich/PRO-GENE-GEN/).

#### Gene co-expression

We employed the hCoCena (horizontal construction of co-expression networks and analysis), an R-package for network-based joint co-expression analysis^62^, to quantify co-expression preservation between the synthetic and real datasets. In these networks, nodes represent genes and edges represent co-expression relationships weighted by Pearson correlation coefficient (*r*). Edges below a user-defined correlation cutoff can be discarded, allowing focus on co-expression relationships across varying strengths. Analyses were performed for each fold, with correlation thresholds of *r* = 0.3 and *r* = 0.5 to evaluate network recovery across datasets. We then quantified co-expression recovery using three complementary metrics. First, the True Positive Rate (TPR) measures the fraction of real co-expression edges correctly recovered in synthetic network: 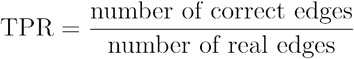. Second, the false edge rate measured the number of spurious edges (false correlations) introduced by the synthetic network relative to the real network: 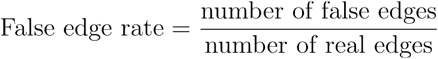. Finally, we defined a co-expression specificity score to summarize network fidelity 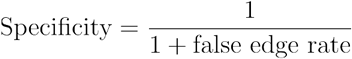 where higher values indicate fewer spurious edges per true edges. Together, higher TPR and higher specificity indicate that the synthetic dataset preserves the real co-expression structure while minimizing falsely introduced correlations.

#### Differential expression

Differential expression (DE) between molecular subtypes (TCGA-BRCA) or cancer types (TCGA-COMBINED) was evaluated for all pairwise class comparisons using the nonparametric Wilcoxon test implemented in the scran R package ^63^. This yielded 10 pairwise molecular subtype comparisons in TCGA-BRCA (e.g. HER2-LumA) and 66 pairwise cancer type comparisons in TCGA COMBINED (e.g. Skin-Colorectal) dataset. DE analysis was performed in the same VST-transformed space for both real and synthetic datasets.

Up- and down-regulated genes were analyzed separately for each comparison. Genes were considered differentially expressed (DE) if they met both a statistical significance threshold (adjusted p ≤ 0.05, Benjamini–Hochberg FDR correction with (α = 0.05) and a minimum effect-size threshold (ΔVST), implemented via the lfc argument in pairwise Wilcoxon function in scran R-package on VST-transformed expression values. The effect-size threshold corresponds to the minimum difference in median VST expression between groups required for a gene to be called DE, with the thresholds of 0.0 and 0.5 for BRCA, and 0.0, 0.5, 1.0 for COMBINED, respectively, used to assess both sensitivity and moderate shifts.

True Positive Rate (TPR) was defined as the proportion of real DE genes correctly identified in the synthetic data, while False Positive Rate (FPR) was the proportion of non-DE genes incorrectly called as DE. Metrics were computed separately for upregulated and downregulated genes. For DE preservation per direction (up and down), a maximum FPR threshold of 0.05 was fixed, and the corresponding TPR was taken as the DE preservation score used in correlation analyses across metrics.

## Privacy

### Membership Inference Attacks

For privacy evaluation, we adopted a suite of complementary membership inference attack (MIA) methods to assess whether the generative models exhibit memorization or disproportionate influence from individual training samples. We implemented these attacks using the DOMIAS package (https://github.com/holarissun/DOMIAS), extending it where necessary for our setting.

All attacks were performed in a black-box, offline setting, where the adversary has access only to the released synthetic data and a set of candidate records containing both members (training samples) and non-members. In other words, the attacker does not observe generative model parameters, gradients or latent representations and operates only on the synthetic outputs. For some attacks, we additionally provide the adversary with a disjoint reference cohort drawn from the same population, representing auxiliary population-level information rather than additional training data. We used the whole gene expression dataset (training + test dataset) as our membership inference test dataset.

For the TCGA-BRCA dataset, we applied two distance-based memorization attacks: GAN-leaks ^26^ and Monte Carlo (MC) ^64^. Both methods operate in the data space and quantify whether a candidate sample lies unusually close to the synthetic samples generated by the model. Gan-leaks scores each sample (record) based on its minimum distance to the synthetic dataset, whereas MC counts the fraction of synthetic samples within an adaptive distance threshold derived from the empirical nearest neighbor distribution.

For the TCGA-COMBINED dataset, the larger cohort size enabled the use of reference-anchored attacks that require auxiliary population information. We withheld 824 samples as a reference dataset that was never used in the training or evaluation of the generative models. Using this reference set, we evaluated GAN-leaks calibrated ^26^, LOGAN-D1 ^65^, and DOMIAS-KDE ^66^. LOGAN-D1 and calibrated GAN-leaks contrast candidate records jointly against the synthetic and the reference set distributions, while DOMIAS-KDE estimates a density ratio between the two using PCA-reduced features (n=150) and kernel density estimation to detect regions where the generator oversamples training-like signals.

In addition to distance and density-based attacks, we also included confidence-based downstream attacks ^67^. For these we trained classifiers on the synthetic data (Logistic Regression and Random Forest) and used the model’s maximum predictive confidence on each candidate as a membership score. This attack probes whether models trained on synthetic data retain higher confidence on records resembling training members relative to the non-members.

### Experiment Setup

The real dataset served as the membership query set and was evaluated against the corresponding synthetic data generated by each model. Membership inference risk was assessed separately for each fold under five-fold cross-validation. Since 80% of the data are used for training in each fold, the positive (member) class prevalence is 0.8.

### Metrics

Each attack returns a continuous membership probability where higher means more likely a member. To assess vulnerability, we use multiple metrics, including AUC-ROC and AUC-PR, which are standard measures of attack success. However, because the membership test set includes the entire dataset and only 20% of samples are treated as non-members under the 5-fold cross-validation setup, the resulting class imbalance can bias these metrics.

Another important consideration is that both AUC and AUPR provide average-case estimates of privacy risk, which may obscure high-risk scenarios where certain individuals are particularly exposed. To capture worst-case behaviour more effectively, we also report metrics where the true positive rate (TPR) is computed at fixed false positive rates (FPR), such as 0.1 and 0.01. These measurements offer a more practical sense of privacy risk; for example, the attacker’s success rate in correctly identifying members when only 10% of non-members are misclassified.

In addition to attack-based metrics, similarity-based measures are often used to evaluate potential privacy risks. For example, the distance to the nearest real neighbor computes the average distance from each synthetic data point to its closest real counterpart. While GAN-leaks is also based on a nearest-neighbor distance, it differs in that it is computed per candidate record in the membership test set, producing a membership score rather than a dataset-level average.

### Correlation analyses across evaluation metrics

We assessed pairwise correlations between evaluation metrics using model-level summary results. For each generative method, performance was averaged across five cross-validation folds prior to analysis, yielding one aggregate value per metric per method. Consequently, the effective sample size for correlation analysis was *n* = 11, corresponding to the number of evaluated methods.

Spearman’s rank correlation (ρ) was used to quantify monotonic associations between metrics. Statistical significance was evaluated using two-sided tests, and p-values were adjusted for multiple comparisons using the Benjamini–Hochberg procedure to control the false discovery rate (FDR) at α = 0.05. Adjusted p-values (q-values) are reported, with significance thresholds denoted as, * = *q* < 0.05, * * = *q* < 0.01, and * * * = *q* < 0.001.

Three correlation panels were computed: (i) Fidelity and downstream utility metrics (12 metrics total), presented in Supplementary Figures S1-S2, (ii) privacy-focused analysis, including seven metrics comprising privacy measures together with selected fidelity and utility metrics (Supplementary Figures S11-S12), (iii) cross evaluation-axis integration, including twelve representative metrics spanning all four evaluation axes (fidelity, utility, biological plausibility, and privacy), shown in Supplementary Figures S13-S14.

## Supporting information

Supplementary File 1_ Challenge details and Tables

## Supplementary information

Additional file 1: Documentation of the challenge structure and supplementary tables. Additional file 2: Supplementary figures.

## Acknowledgements

We would like to thank Dimosthenis Karatzas for his helpful feedback during the design stage of the Health Privacy Challenge; Sergi Robles for his support in setting up the challenge tasks at the ELSA Benchmark Platform; Paweł Łabaj and David P. Kreil for facilitating the smooth integration of the challenge with CAMDA; Katharina Mikulik for her thoughtful feedback on the challenge GitHub repository use cases; Danai Vagiaki for kindly sharing her experience with challenge tasks involving the second track; and Kevin Domanegg for his helpful feedback on the challenge communication.

H.Ö. acknowledges EMBL HPC, A.W. acknowledges DKFZ HPC and Karol Nowicki-Osuch (DKFZ), J.K. et al. acknowledges the Institute for Bioinformatics and Medical Informatics (IBMI) of the University of Tübingen, S.M., S.P et al., acknowledges National Artificial Intelligence Research Resources (NAIRR) for the provided computational resources and infrastructure.

## Authors’ contributions

H.Ö, T.A, J.J., R.B., M.F., O.S and A.H. conceived and designed the community challenge. H.Ö and T.A implemented the pre- and post-challenge generative methods. H.Ö. designed and implemented the evaluation framework, conducted the experiments, and performed the analyses. J.S.R., P.R.M. and S.L. provided conceptual feedback on the challenge design. P.R.M. contributed to code review and supported scalable implementation. S.L. contributed to code review. H.Ö. drafted the manuscript. All other co-authors developed the generative methods participating in the challenge, provided CAMDA abstracts and synthetic data: A.W. contributed NMF and DP-NMF; J.K. et al. contributed Embedded Diffusion; S.M. and S.P. et al., contributed P-PGM. All authors provided critical feedback, contributed to revisions, and approved the final version of the manuscript.

## CAMDA 2025 Health Privacy Challenge

### Team Differentially-Private NMF

Andrew Wicks

### Team Methods in Medical Informatics

Jules Kreuer, Sofiane Ouaari, Nico Pfeifer

### Team University of Washington Tacoma and Sage Bionetworks

Shane Menzies, Sikha Pentyala, Daniil Filienko, Steven Golob, Patrick McKeever, Jineta Banerjee, Luca Foschini, Martine De Cock

### Organizers

Hakime Öztürk, Tejumade Afonja, Joonas Jälkö, Ruta Binkyte, Pablo Rodriguez-Mier, Sebastian Lobentanzer, Julio Saez-Rodriguez, Mario Fritz, Oliver Stegle, Antti Honkela

## Data availability

We used open access bulk gene-expression data from The Cancer Genome Atlas (TCGA), accessed via the Genomic Data Commons (GDC) portal (https://portal.gdc.cancer.gov) (Methods). The pre-processed datasets generated, analysed and supporting the conclusions of this article are publicly available at the ELSA Benchmark platform (https://benchmarks.elsa-ai.eu/?ch=4) upon completion of a data access agreement. The baseline and post-challenge methods as well as all evaluation metrics are available in the challenge Github repository (https://github.com/PMBio/Health-Privacy-Challenge). The code is written for Python 3.10 and R 4. on Linux/Unix.

The participant submitted methods are available in the corresponding repos, compliant with the challenge format. Embedded Diffusion: https://github.com/not-a-feature/CAMDA25_NoisyDiffusion, NMF/DP-NMF: https://github.com/AndrewJWicks/SingleCellNMFGenerator, P-PGM: https://github.com/sikhapentyala/Health-Privacy-Challenge/tree/main/submission The results from the 2025 challenge are publicly accessible at the ELSA benchmark platform (https://benchmarks.elsa-ai.eu/?ch=4). The CAMDA 2026 edition of Health Privacy Challenge (https://benchmarks.elsa-ai.eu/?ch=8) remains as an open, ongoing challenge, serving as a living benchmark for method evaluation within a standardized framework.

## Declarations

### Ethics approval and consent to participate

Not applicable.

### Consent for publication

Not applicable.

