## Supplementary File 1_ Challenge details and Tables for "Towards Useful and Private Synthetic Omics: Community Benchmarking of Generative Models for Transcriptomics Data"

**[1]European Molecular Biology Laboratory (EMBL), Genome Biology Unit, Heidelberg, Germany**

**[2]Division of Computational Genomics and Systems Genetics, German Cancer Research Center (DKFZ), Heidelberg, Germany**

**[3]CISPA Helmholtz Center for Information Security, Saarbrücken, Germany**

**[4]University of Helsinki, Finland**

**[5]Heidelberg University, Germany**

**[6]Helmholtz Munich, Germany**

**[7]Division of Tumorigenesis and Molecular Cancer Prevention, German Cancer Research Center (DKFZ), Heidelberg, Germany**

**[8]DKFZ Hector Cancer Institute at the University Medical Center Mannheim, Germany**

**[9]Eberhard Karls Universität Tübingen, Germany**

**[10]University of Washington Tacoma, USA**

**[11]Sage Bionetworks, Seattle, USA**

**[12]Ghent University, Ghent, Belgium**

**[13]European Bioinformatics Institute (EMBL-EBI), UK**

### Supplementary Materials

#### Challenge infrastructure

The Health Privacy Challenge was launched in conjunction with the annual international Critical Assessment of Massive Data Analysis (CAMDA) 2025 conference

(https://bipress.boku.ac.at/camda2025/), which runs together with the Intelligent Systems for Molecular Biology (ISMB) conference series.
The challenge comprised two tracks: Track-1 focused on synthetic and private bulk RNA-seq generation, re-distributing two datasets obtained from the GDC portal (https://portal.gdc.cancer.gov) containing The Cancer Genome Atlas (TCGA) [1] cohorts. Track 2, on the other hand, focused on synthetic single-cell RNA generation and re-distributed raw counts of OneK1K single-cell RNA-seq dataset (https://onek1k.org/), a cohort containing 1.26 million peripheral blood mononuclear cells (PBMCs) of 14 cell types from 981 donors[2]. The right for the distribution was kindly granted by the authors. These datasets are available for download in the benchmark platform upon signing a data download agreement.

### Track 1 (Bulk RNA-seq) - Leaderboard format

Track 1 was structured as a "Blue Team vs. Red Team" challenge, and implemented in two sequential phases to ensure that Red Teams had a sufficiently large and diverse set of Blue Team submissions to engage with.

#### Blue Team setup

During the first phase (Phase I), Blue Teams developed generative models aimed at improving upon the provided baseline approaches and advancing methodological insight into privacy preservation for biological datasets.
The participants were provided with the re-distributed TCGA bulk RNA-seq datasets, denoted as $D$. Each team trained a generative model $f(D)$ to produce synthetic data derived from the original dataset. To prevent information leakage and enforce methodological independence:

- Each Blue Team selected a private, team-specific random seed, disclosed only to the organizers at submission,
- The seed determined the train-test split of $D$.
- The same split was used to train the generative model $f(D)$.

**Cross-validation protocol:** A five-fold cross-validation setting was utilized, where shape-matched synthetic datasets $D_{SYN}^{(i)}$ are generated from the model $f(D)$ trained on the training set $D_{TR}^{(i)}$ of each fold $i \in \{1, 2, 3, 4, 5\}$. The quality of the synthetic datasets were evaluated on the corresponding held-out test set $D_{TE}^{(i)}$. Performance metrics were averaged across folds to obtain a stable and generalizable estimate of the model performance.
**Membership Inference Setup:** For each cross-validation fold, a dedicated membership inference test dataset was constructed for subsequent use by Red Teams. Across folds:

- Members corresponded to samples included in the training partition.
- Non-members corresponded to samples in the held-out test partition.

- Membership composition varied by fold.

As five-fold cross-validation was employed, a total of five membership inference datasets were generated per Blue Team submission. By design, members constituted 80% of each inference dataset, reflecting 80/20 train test split.

Because each Blue Team used a distinct random seed, membership labels differed across teams. This avoided a fixed global split and introduced variability essential for robust Red Team evaluation.

**Timeline:** Blue Teams were only permitted to submit their methods during the Phase I of Track 1, which ran from January 14th 2025 to March 15th 2025.

#### Red Team Setup

Track 1 incorporated the Red Team component in two sequential phases aligned with the Blue Team workflow.

**Phase I - Preparation and baseline attacks:** During the first phase of Track 1, Red Teams were invited to launch membership inference attacks (MIA) against synthetic datasets generated using the baseline generative models provided by the organizers. This phase served two primary purposes: (i) to identify Red Teams intending to participate in Phase II, and (ii) to allow participants to develop, calibrate and validate their MIA pipelines prior to evaluating Blue Team submissions.

**Phase II - Evaluation of Blue Team submissions:** In the second phase, Red Teams launched MIA on the synthetic datasets and corresponding white-box generative models submitted by the Blue Team participants.

Here, white-box access indicates that Red Teams were provided with full visibility to model architecture, training procedure, and hyperparameters. The only withheld elements were the random seed used to generate the train–test partition and the stochastic seed governing model initialization and training randomness. In some cases, these seeds may have coincided; however, neither was disclosed to the Red Teams.

- Four Blue Team submissions were selected based on leaderboard performance on the benchmark platform, and shared with the Red Teams.
- Each Red Team was required to launch their MIA algorithm against at least one Blue Team model.
- If a Red Team did not explicitly attack a given Blue Team model, that model was evaluated using a predefined baseline MIA algorithm based on a Monte Carlo (MC) strategy [3].
- If a team participated in both roles (Blue and Red), and their own Blue Team model was among the selected submissions, baseline MC performance was used in place of a self-attack to avoid conflicts of interest.
- Final Red Team attack performance was computed as the average performance across all evaluated Blue Team models, combining participant-submitted MIAs where provided and baseline MC evaluations where attacks were not conducted.

**Timeline:** Phase II ran from March 22, 2025 to May 12, 2025.

##### Evaluation

For Track 1, the submission platform executed a Docker container in the background, prepared and maintained by the organizers, to ensure a standardized and reproducible evaluation. Upon submission, the container automatically evaluated provided synthetic datasets using the predefined metrics specified in the official challenge Github repository. Each submission was required to include:

- A short methodological description,
- Python source files implementing the method,
- A configuration file specifying the hyperparameters used for training and generation, and
- An environment specification file to ensure computational reproducibility.

The resulting performance metrics were then published on the public leaderboard, making them visible to participants in both the Blue Team and Red Team tracks, as well as external observers interested in the challenge outcomes. This setup ensured transparency and consistency across submissions.

#### Track 2 (Single-cell RNA-seq) - Open research format

Track 2 was structured as a single phase challenge, running from January 27th, 2025 to May 12th, 2025. Given the substantial size and storage footprint of the generated single-cell RNA-seq datasets, synthetic datasets were not centrally collected, and no public leaderboard was implemented on the submission platform. Instead, the focus was placed on methodological transparency and reproducibility. Participants were required to submit:

- A short methodological description,
- Python source files implementing their methods
- Configuration file specifying hyperparameters, and
- An environment specification file to ensure computational reproducibility.

Rather than operating as a leaderboard-driven benchmark, Track 2 was positioned as an open research problem. Submitted approaches were evaluated and discussed through CAMDA extended abstract submissions.

#### Logistics

**CAMDA extended abstract:** Participants of the both tracks were required to submit an accompanying CAMDA extended abstract by May 15, 2025 through the submission platform provided by the ISMB/ECCB 2025 conference (https://www.iscb.org/ismbeccb2025/home). Submissions were assessed using task-specific, multi-metric evaluation protocols designed to prevent over-optimization toward a single benchmark metric. The extended abstract was required to provide a detailed description of the proposed method, experimental design, and analytical findings, thereby enabling rigorous qualitative and quantitative assessment beyond leaderboard scores.

For Track 2 in particular, these abstracts carried increased weight due to the open-research format and the absence of a centralized leaderboard. As synthetic datasets were not collected centrally, methodological clarity, reproducibility documentation, and analytical justification presented in the abstract became central to the evaluation process.

**Github Starter Kit:** The participants were provided with a Github Starter Kit repository (https://github.com/PMBio/Health-Privacy-Challenge) that contained baseline methods, evaluation metrics, and guidelines to prepare their submissions for respective tracks. They were asked to follow the template provided in the repository based Python to ensure reproducibility of their submission.

**Submission Platform:** The submissions were collected through the ELSA Benchmarking Platform (https://benchmarks.elsa-ai.eu/?ch=4), where participants submitted the required files and thus, officially participated in the challenge.

### TABLES

**Table S1: Complete list of evaluation metrics used to assess synthetic dataset quality.** Metrics are grouped into categories (e.g., utility, fidelity), with primary metrics highlighted for interpretation in the main manuscript and intermediate metrics providing additional diagnostic information reported in the supplementary material. Arrows indicate the desired optimization direction: upward (↑) for higher-is-better scores and downward (↓) for lower-is-better scores. Some of the metrics are computed on both training and held-out test partitions to evaluate generalization and detect potential overfitting. The public repository includes additional scripts for exploratory visualization and diagnostic analyses (e.g., PCA), which were not used in the reported results but are available for reproducibility and further exploration.

| **Category** | **Method Detail** | **Description** | **Role** | **Direction** |
|---|---|---|---|---|
| Utility | Accuracy (Synthetic) | Downstream task performance for Train on Synthetic, Test on Real (TSTR) | Intermediate | ↑ |
| Utility | Accuracy (Real) | Downstream task performance for Train on Real, Test on Real (TRTR) | Intermediate | ↑ |
| Utility | Accuracy Relative | $\text{Accuracy}_{TSTR}/\text{Accuracy}_{TRTR}$ | Primary | ↑ |
| Utility | AUROC (Synthetic) | TSTR | Intermediate | ↑ |
| Utility | AUROC (Real) | TRTR | Intermediate | ↑ |
| Utility | AUROC Relative | $\text{AUROC}_{TSTR}/\text{AUROC}_{TRTR}$ | Primary | ↑ |
| Utility | PR Macro (Synthetic) | TSTR | Intermediate | ↑ |

| | | | | |
|---|---|---|---|---|
| Utility | PR Macro (Real) | TRTR | Intermediate | ↑ |
| Utility | PR Macro Relative | $\text{PR Macro}_{TSTR}/\text{PR Macro}_{TRTR}$ | Primary | ↑ |
| Utility | F1 (Synthetic) | TSTR | Intermediate | ↑ |
| Utility | F1 (Real) | TRTR | Intermediate | ↑ |
| Utility | F1 Relative | $\text{F1}_{TSTR}/\text{F1}_{TRTR}$ | Primary | ↑ |
| Utility | Overlapping features % | Proportion of shared important features identified by TSTR and TRTR models | Primary | ↑ |
| Fidelity | Maximum Mean Discrepancy (MMD) (test) | Kernel-based distance between synthetic data and held-out real test data distributions | Intermediate | ↓ |
| Fidelity | Maximum Mean Discrepancy (MMD) (train) | Kernel-based distance between synthetic data and real training data distributions | Intermediate | ↓ |
| Fidelity | Inverted MMD (test) | Transformed MMD score to align with “higher-is-better” interpretation: $1/(1 + \text{MMD (test)})$ | Primary | ↑ |
| Fidelity | Kullback-Leibler (KL) divergence (test) | Divergence between synthetic and held-out real test data distributions | Intermediate | ↓ |
| Fidelity | Kullback-Leibler (KL) divergence (train) | Divergence between synthetic and real training data distributions | Intermediate | ↓ |
| Fidelity | Inverted KL-divergence (test) | Transformed KL divergence score $1/(1 + \text{KL (test)})$ | Primary | ↑ |
| Fidelity | Discriminative score F1 | F1 score for distinguishing synthetic and real dataset | Intermediate | ↓ |
| Fidelity | Inverted Discriminative score | Transformed discriminative score, where higher values indicate high indistinguishability: $1 - \text{discriminative score}$ | Primary | ↑ |
| Fidelity | Distance-to-Closest | Average distance of synthetic dataset to the nearest real test data point | Intermediate | ↓ |
| Fidelity | Distance-to-Closest (baseline) | Average distance within real train dataset to the nearest test data point | Intermediate | ↓ |
| Fidelity | Distance-to-Closest Relative | Ratio of synthetic-to-real distances relative to the baseline, capturing fidelity normalized to intrinsic dataset structure, ideal value ≈ 1 | Primary | ↓ (~1) |
| Privacy | Area Under ROC Curve | The predictive performance of the membership inference attack (MIA) method | Primary | ↓ |

| | | | | |
|---|---|---|---|---|
| Privacy | Area Under Precision Recall Curve | The predictive performance of the membership inference attack (MIA) method | Intermediate | ↓ |
| Privacy | TPR at FPR=0.1 | The ability of the attack to identify correct members when FPR is fixed at 10% | Primary | ↓ |
| Privacy | TPR at FPR=0.1 | The ability of the attack to identify correct members when FPR is fixed at 1% | Intermediate | ↓ |
| Biological Plausibility | Differential Expression (DE) True positive rate (TPR) | Agreement between synthetic and real datasets in detecting DE genes (e.g. between two subtypes) using Wilcoxon rank-sum test | Primary | ↑ |
| Biological Plausibility | DE False positive rate (FPR) | Rate of falsely detected DE genes in synthetic data compared to real data | Primary | ↓ |
| Biological Plausibility | Co-expression TPR | Agreement between synthetic and real datasets in detecting co-expression network edges using hCoCena package | Primary | ↑ |
| Biological Plausibility | Co-expression False edge rate | Number of falsely detected co-expression edges in synthetic data normalized by the number of edges in real network | Primary | ↓ |
| Biological Plausibility | Co-expression Specificity | Transformed false edge rate for better readability, $1/(1+\text{false edge rate})$ | Primary | ↑ |

**Table S2: Fidelity and utility metrics for TCGA-BRCA dataset.** Models labeled "+" use an embedding layer for input encoding, while "−" indicates one-hot encoding.

| **Method** | **Fidelity** | | | | **Utility** | | | |
|---|---|---|---|---|---|---|---|---|
| | **MMD (test data) ↓** | **Avg. KL (test data) ↓** | **Discrim. Score ↓** | **Distance to closest ↓** | **Feature Overlap (%) ↑** | **AU-PR Relative ↑** | **AUROC Relative ↑** | **F1 Relative ↑** |
| **Real data** | 0.00 | - | - | | - | 1.000 | 1.000 | 1.000 |
| **Baseline selected methods** | | | | | | | | |
| **MVN** | 0.018 | **0.215** | **0.528** | **28.534** | **34.1** | **0.978** | **0.988** | **1.000** |
| **CVAE +** | 0.024 | 0.649 | 0.787 | 15.441 | 27.2 | 0.959 | 0.985 | 0.981 |
| **CTGAN -** | 0.092 | 5.761 | 0.996 | | 0.056 | | | |
| **DP-CVAE ($\epsilon = 10$) +** | 0.146 | 0.500 | 1.000 | 43.559 | 3.6 | 0.514 | 0.781 | 0.780 |
| **DP-CTGAN ($\epsilon = 45$)** | 1.113 | 19.925 | 1.000 | 91.417 | 5.4 | 0.274 | 0.463 | 0.318 |
| **Submissions** | | | | | | | | |
| **NMF** | **0.002** | 0.286 | 0.683 | 27.738 | 7.2 | 0.665 | 0.872 | 0.829 |
| **DP-NMF** $\epsilon = 2.8$ | 0.005 | 0.284 | 0.755 | 28.758 | 4.4 | 0.583 | 0.797 | 0.746 |
| **Embedded Diffusion** | 0.013 | 0.406 | 0.632 | 21.483 | 0.302 | 0.904 | 0.984 | 0.948 |
| **P-PGM ($\epsilon = 10$)** | 0.007 | 18.495 | 0.765 | 27.077 | 0.056 | 0.873 | 0.970 | 0.927 |
| **Post challenge methods** | | | | | | | | |
| **CVAE-GMM +** | 0.052 | 1.397 | 0.693 | 16.773 | 0.318 | 0.938 | 0.982 | 0.975 |
| **WGAN-GP+** | 0.032 | 0.830 | 0.705 | 19.361 | 0.137 | 0.918 | 0.967 | 0.938 |

**Table S3: Fidelity and utility metrics for TCGA-COMBINED dataset.** Models labeled "+" use an embedding layer for input encoding, while "−" indicates one-hot encoding.

| **Method** | **Fidelity** | | | | **Utility** | | | |
|---|---|---|---|---|---|---|---|---|
| | **MMD (test data) ↓** | **KL (test data) ↓** | **Discrm. Score ↓** | **Distance to closest ↓** | **Feature Overlap (%) ↑** | **AU-PR Relative ↑** | **AUROC Relative ↑** | **F1 Relative ↑** |
| **Real data** | -0.000 | 0.052 | - | 23.272 | - | 1.000 | 1.000 | 1.000 |
| **Baseline selected methods** | | | | | | | | |
| **Multivariate** | 0.009 | **0.074** | **0.577** | 28.877 | **0.471** | **0.999** | **0.999** | **0.999** |
| **CVAE +** | 0.025 | 0.343 | 0.787 | 16.550 | 0.405 | 0.994 | 0.998 | 0.993 |
| **CTGAN -** | 0.087 | 5.530 | 0.996 | 26.323 | 0.147 | 0.113 | 0.513 | 0.085 |
| **DP-CVAE ($\epsilon = 10$)** | 0.095 | 0.316 | 0.999 | 58.130 | 0.151 | 0.622 | 0.947 | 0.893 |
| **DP-CTGAN ($\epsilon = 45$)** | 0.925 | 19.142 | 1.000 | 104.463 | 0.104 | 0.102 | 0.512 | 0.090 |
| **Submissions** | | | | | | | | |
| **NMF** | **0.002** | 0.076 | 0.640 | 28.744 | 0.164 | 0.820 | 0.970 | 0.853 |
| **DP-NMF ($\epsilon = 2.8$)** | 0.002 | 0.077 | 0.677 | 28.992 | 0.175 | 0.787 | 0.963 | 0.831 |
| **Embedded Diffusion** | 0.006 | 0.100 | 0.722 | 22.097 | 0.369 | 0.994 | 0.998 | 0.992 |
| **P-PGM ($\epsilon = 10$)** | 0.005 | 17.641 | 0.620 | 29.213 | 0.184 | 0.966 | 0.994 | 0.948 |
| **Post challenge methods** | | | | | | | | |
| **CVAE-GMM +** | 0.025 | 0.538 | 0.789 | **16.179** | 0.368 | 0.991 | 0.996 | 0.992 |
| **WGAN-GP+** | 0.014 | 0.158 | 0.953 | 20.130 | 0.240 | 0.987 | 0.997 | 0.983 |

**Table S4: Ablation analysis of input encoding strategies for the TCGA-BRCA dataset, comparing one-hot encoding and embedding-layer representations.** Fidelity and utility metrics are reported for both variants; "+" denotes embedding-based encoding and "−" denotes one-hot encoding.

| Method | Fidelity | | | | Utility | | | |
|---|---|---|---|---|---|---|---|---|
| | MMD (test data) ↓ | KL (test data) ↓ | Discrm. Score ↓ | Distance to closest ↓ | Feature Overlap (%) ↑ | AU-PR Relative ↑ | AUROC Relative ↑ | F1 Relative ↑ |
| **Real data** | 0.00 | 0.234 | - | 24.043 | - | 1.000 | 1.000 | 1.000 |
| **CVAE +** | 0.024 | 0.649 | 0.787 | 15.441 | 27.2 | 0.959 | 0.985 | 0.981 |
| **CVAE -** | 0.023 | 0.648 | 0.783 | 16.026 | 27.3 | 0.968 | 0.991 | 0.992 |
| **CVAE-GMM +** | 0.054 | 1.397 | 0.693 | 16.773 | 31.8 | 0.938 | 0.982 | 0.975 |
| **CVAE-GMM -** | 0.056 | 1.506 | 0.686 | 16.742 | 22.2 | 0.833 | 0.951 | 0.906 |
| **WGAN-GP+** | 0.033 | 0.830 | 0.705 | 19.361 | 13.7 | 0.918 | 0.967 | 0.938 |
| **WGAN-GP-** | 0.044 | 0.941 | 0.683 | 19.124 | 5.2 | 0.354 | 0.639 | 0.489 |

**Table S5: Ablation analysis of input encoding strategies for the TCGA-COMBINED dataset, comparing one-hot encoding and embedding-layer representations.** Fidelity and utility metrics are reported for both variants; "+" denotes embedding-based encoding and "−" denotes one-hot encoding.

| Method | Fidelity | | | | Utility | | | |
|---|---|---|---|---|---|---|---|---|
| | MMD (test data) ↓ | KL (test data) ↓ | Discrm. Score ↓ | Distance to closest ↓ | Feature Overlap (%) ↑ | AU-PR Relative ↑ | AUROC Relative ↑ | F1 Relative ↑ |
| **Real data** | -0.000 | 0.052 | - | 23.272 | - | 1.000 | 1.000 | 1.000 |
| **CVAE +** | 0.025 | 0.343 | 0.787 | 16.550 | 40.5 | 0.994 | 0.998 | 0.993 |
| **CVAE -** | 0.024 | 0.326 | 0.800 | 16.832 | 39.0 | 0.997 | 0.999 | 0.995 |
| **CVAE-GMM +** | 0.026 | 0.538 | 0.789 | 16.179 | 36.8 | 0.991 | 0.996 | 0.992 |
| **CVAE-GMM -** | 0.028 | 0.554 | 0.797 | 16.211 | 34.3 | 0.988 | 0.996 | 0.989 |
| **WGAN-GP+** | 0.014 | 0.158 | 0.953 | 20.130 | 24.0 | 0.987 | 0.997 | 0.983 |
| **WGAN-GP-** | 0.019 | 0.213 | 0.884 | 19.544 | 27.3 | 0.986 | 0.997 | 0.982 |
