## Supplementary File 2_ Figures for "Towards Useful and Private Synthetic Omics: Community Benchmarking of Generative Models for Transcriptomics Data"

**[1]European Molecular Biology Laboratory (EMBL), Genome Biology Unit, Heidelberg, Germany**

**[2]Division of Computational Genomics and Systems Genetics, German Cancer Research Center (DKFZ), Heidelberg, Germany**

**[3]CISPA Helmholtz Center for Information Security, Saarbrücken, Germany**

**[4]University of Helsinki, Finland**

**[5]Heidelberg University, Germany**

**[6]Helmholtz Munich, Germany**

**[7]Division of Tumorigenesis and Molecular Cancer Prevention, German Cancer Research Center (DKFZ), Heidelberg, Germany**

**[8]DKFZ Hector Cancer Institute at the University Medical Center Mannheim, Germany**

**[9]Eberhard Karls Universität Tübingen, Germany**

**[10]University of Washington Tacoma, USA**

**[11]Sage Bionetworks, Seattle, USA**

**[12]Ghent University, Ghent, Belgium**

**[13]European Bioinformatics Institute (EMBL-EBI), UK**

### SUPPLEMENTARY FIGURES

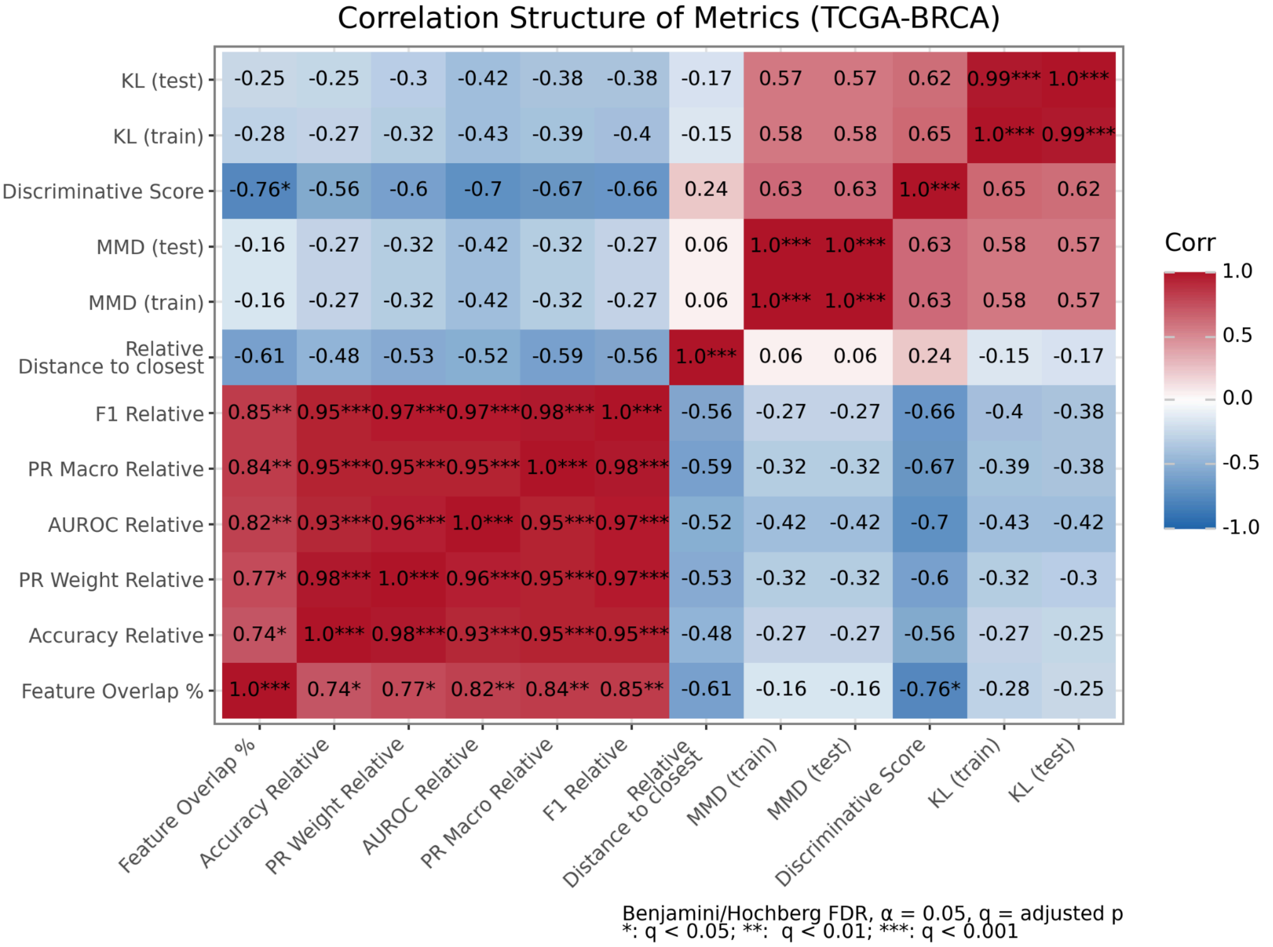


**Supp Figure S1: TGCA-BRCA Spearman correlation heatmap of complete fidelity and downstream utility metrics.** 12 metrics were included, and statistical significance was assessed using the Benjamini-Hochberg procedure controlling the false discovery rate (FDR) at α = 0.05. Significance levels are indicated as follows: *q < 0.05; ** q < 0.01; *** q < 0.001, where q represents the adjusted p-value.

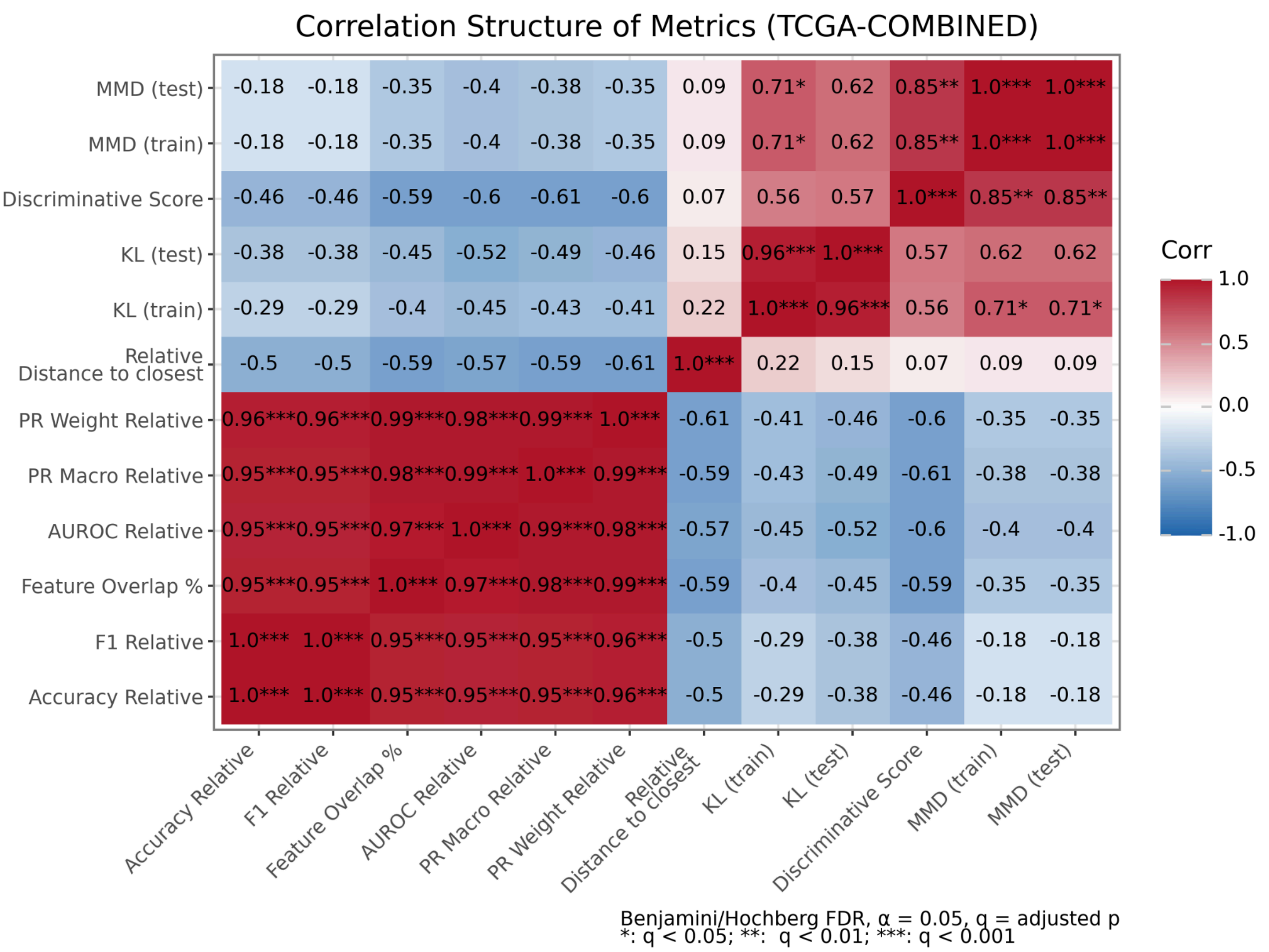


**Supp Figure S2: TGCA-COMBINED Spearman correlation heatmap of complete fidelity and downstream utility metrics.** 12 metrics were included, and statistical significance was assessed using the Benjamini-Hochberg procedure controlling the false discovery rate (FDR) at α = 0.05. Significance levels are indicated as follows: *q < 0.05; ** q < 0.01; *** q < 0.001, where q represents the adjusted p-value.

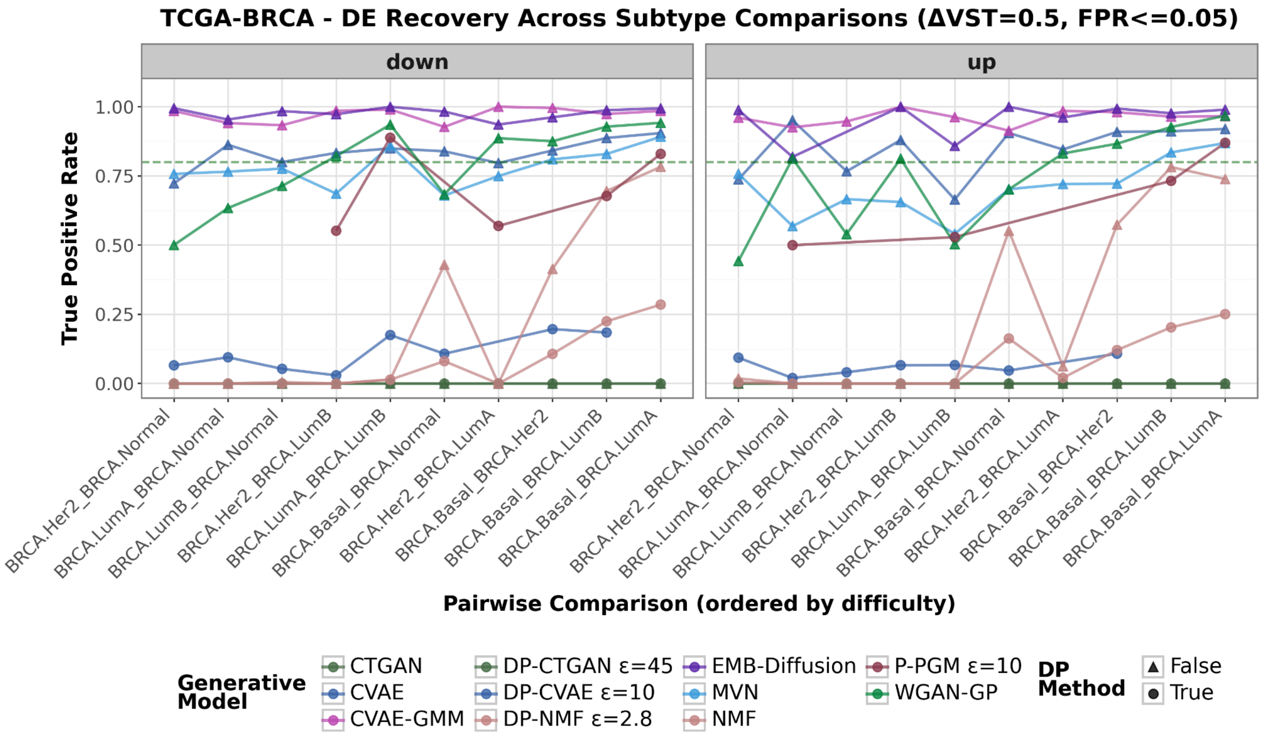


**Supp Figure S3: Differential expression (DE) recovery difficulty across pairs in BRCA dataset.** Differential expression (DE) recovery TPR across cancer type pairwise-comparisons with effect size ($\Delta\mathrm{VST}$) threshold set to 0.5 and FPR <=0.05. Each colored line represents a generative model. DP and non-DP methods are indicated with different symbols.

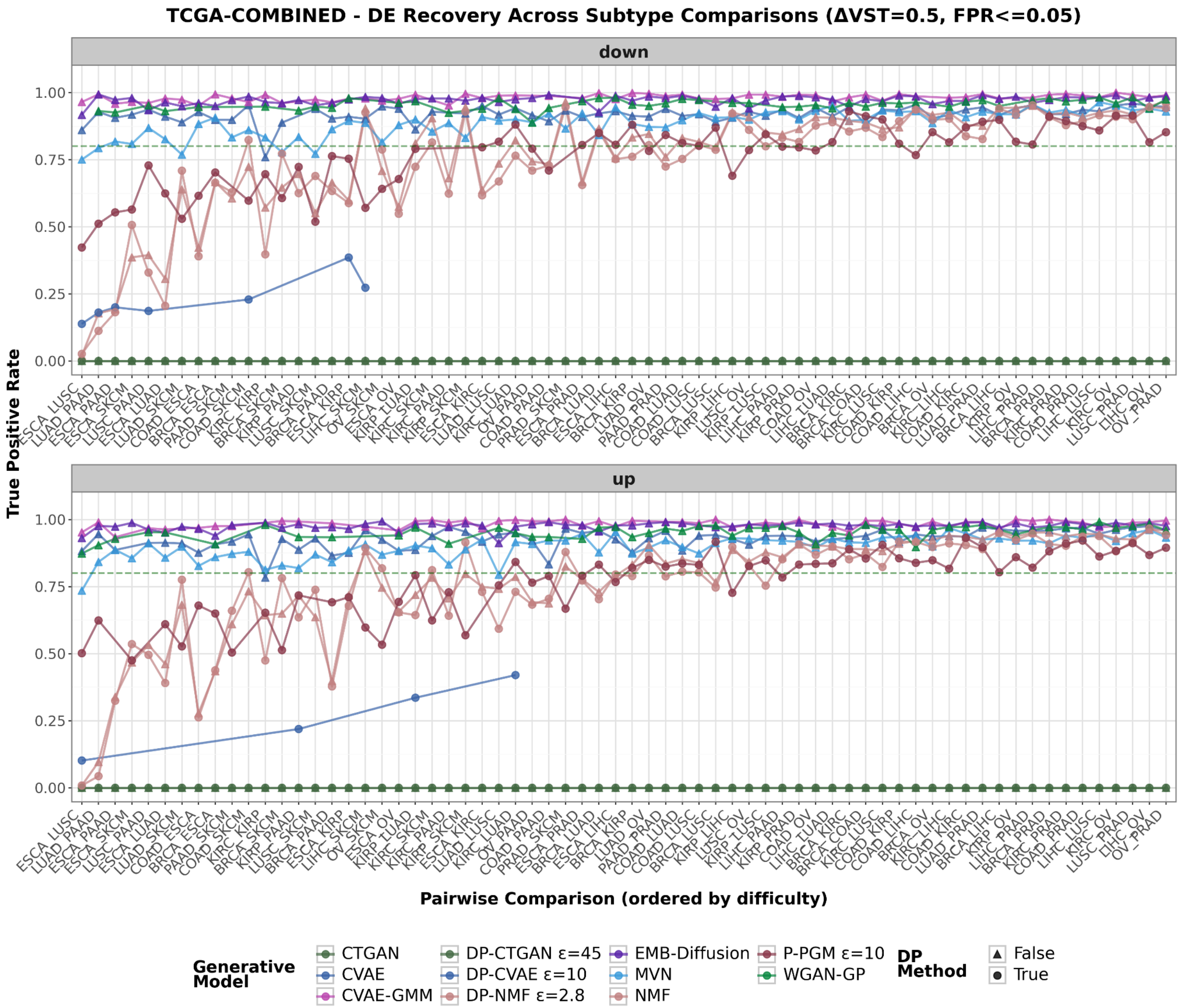


**Supp Figure S4: Differential expression (DE) recovery difficulty across pairs in COMBINED dataset.** TPR across cancer type pairwise-comparisons with effect size ($\Delta\mathrm{VST}$) threshold set to 0.5 and FPR <=0.05. Each colored line represents a generative model. DP and non-DP methods are indicated with different symbols.

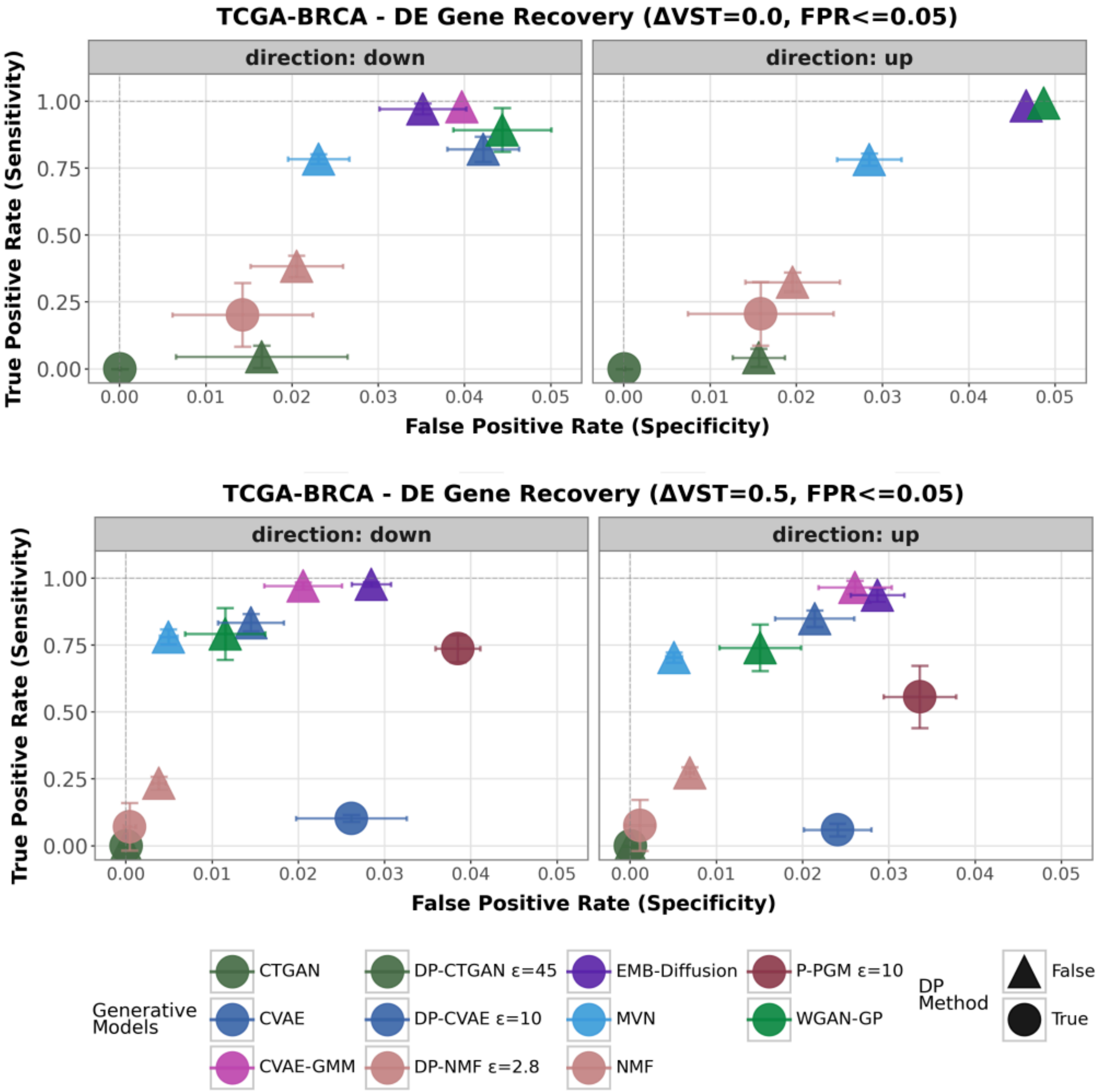


**Supp Figure S5: TCGA-BRCA ablation analysis of differential expression (DE) recovery under varying effect-size thresholds.** Performance is evaluated at thresholds of 0.0 and 0.5, illustrating the sensitivity of DE recovery to increasing effect-size stringency.

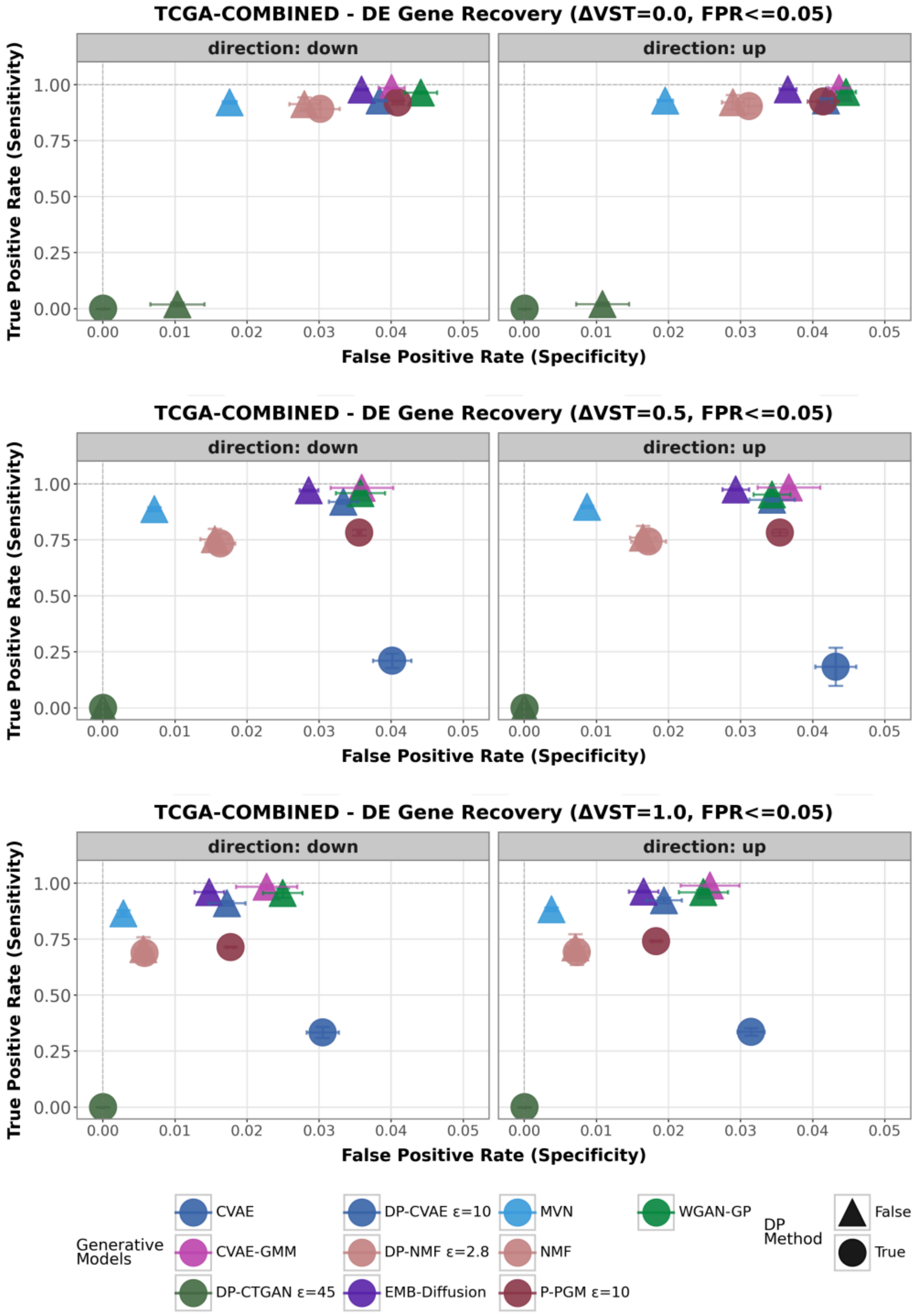


**Supp Figure S6: TCGA-COMBINED ablation analysis of differential expression (DE) recovery under varying effect-size thresholds.** Performance is evaluated at thresholds of 0.0, 0.5, and 1.0, illustrating the sensitivity of DE recovery to increasing effect-size stringency.

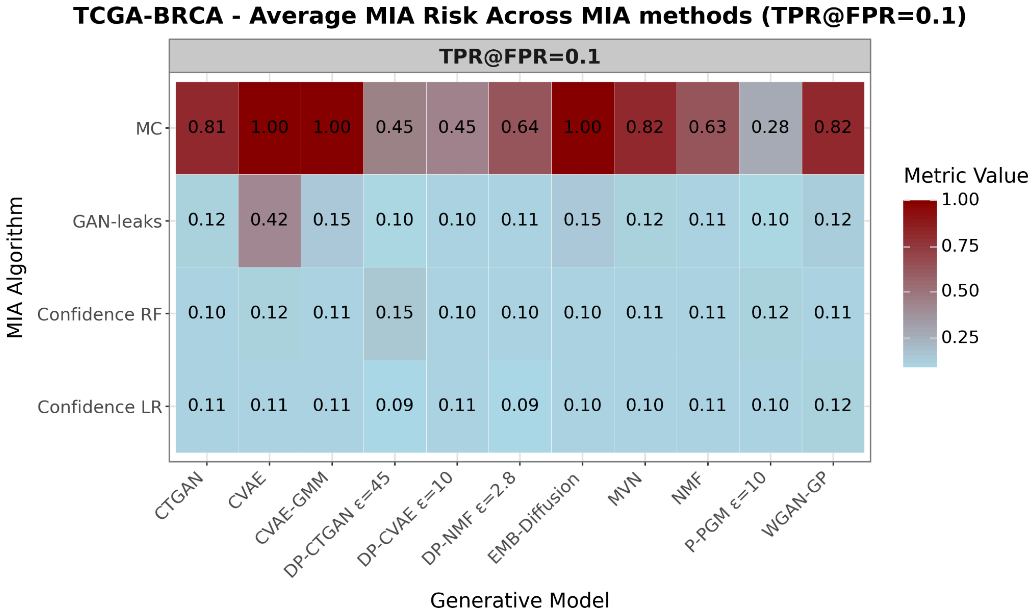


**Supp Figure S7: TCGA-BRCA MIA TPR fixed at TPR = 0.1 performance across four MIA methods.** Membership inference risk was computed at each fold under five-fold cross-validation and the average performance is reported. The baseline performance is equal to 0.1.

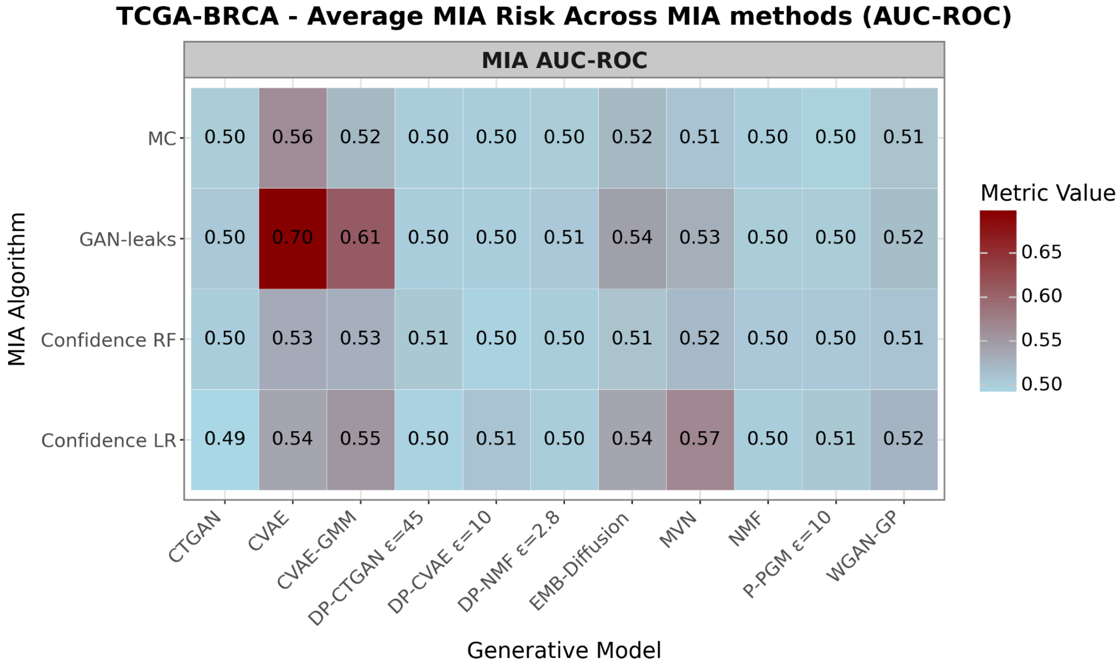


**Supp Figure S8: TCGA-BRCA MIA AUC-ROC performance across four MIA methods.** Membership inference risk was computed at each fold under five-fold cross-validation and the average performance is reported. The baseline AUC-ROC corresponding to random guessing equals to 0.5.

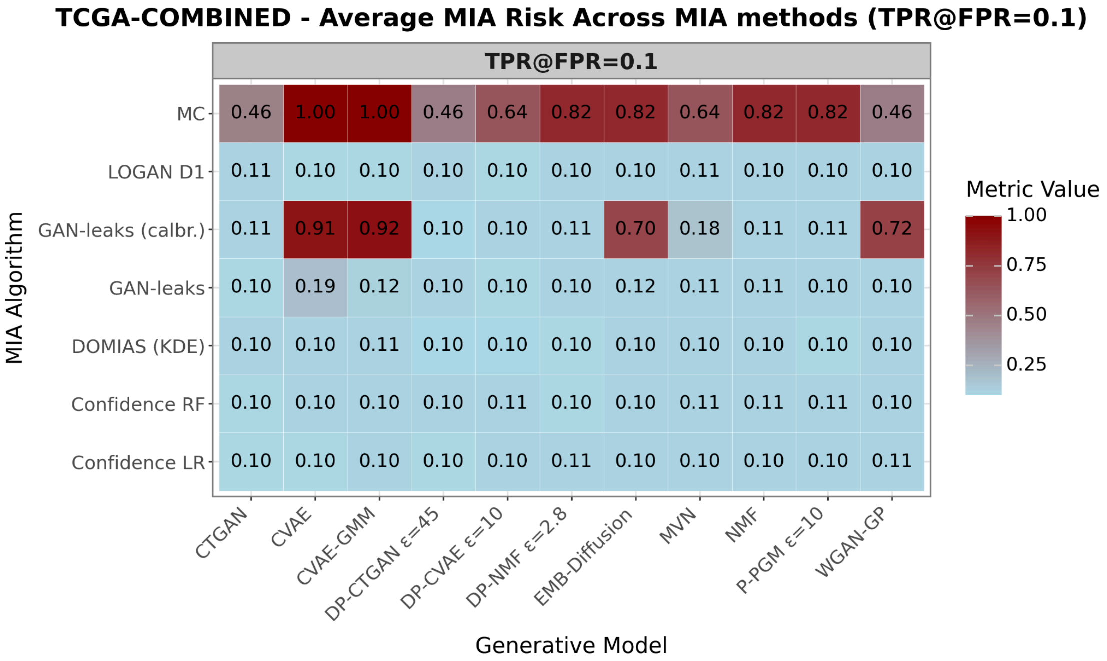


**Supp Figure S9: TCGA-COMBINED MIA TPR fixed at TPR = 0.1 performance across seven MIA methods.** Membership inference risk was computed at each fold under five-fold cross-validation and the average performance is reported. The baseline performance is equal to 0.1.

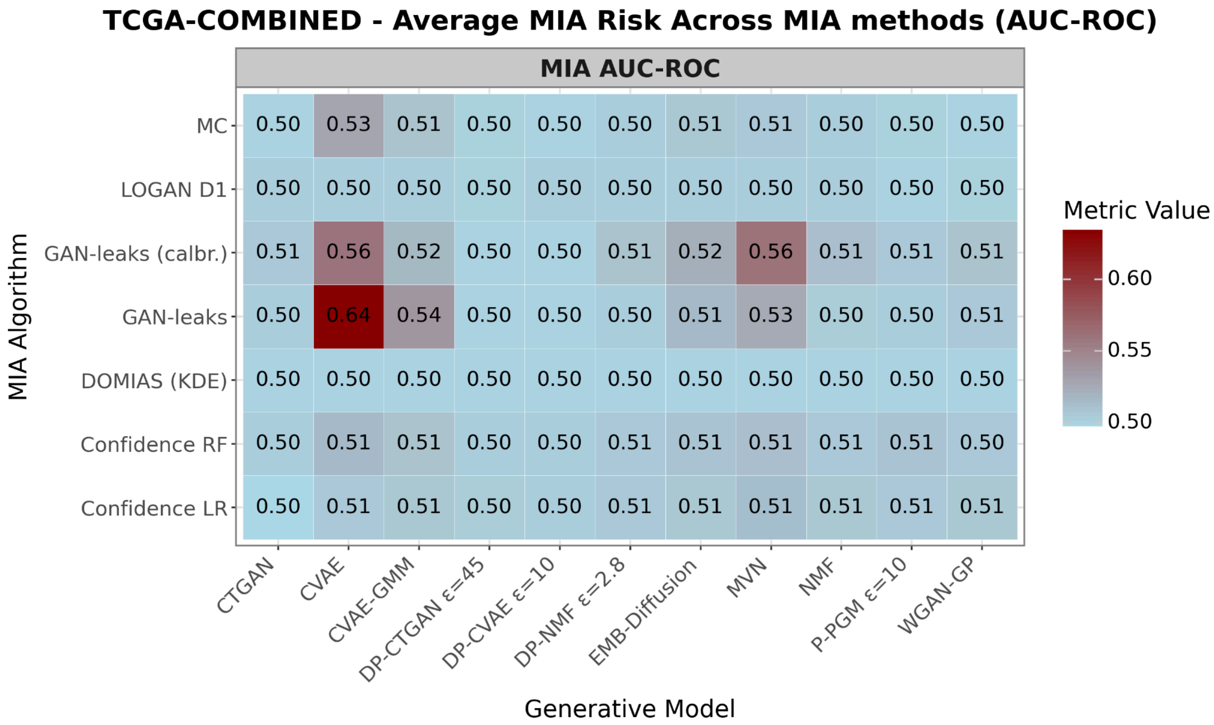


**Supp Figure S10: TCGA-COMBINED MIA AUC-ROC performance across four MIA methods.** Membership inference risk was computed at each fold under five-fold cross-validation and the average performance is reported. The baseline AUC-ROC corresponding to random guessing equals to 0.5.

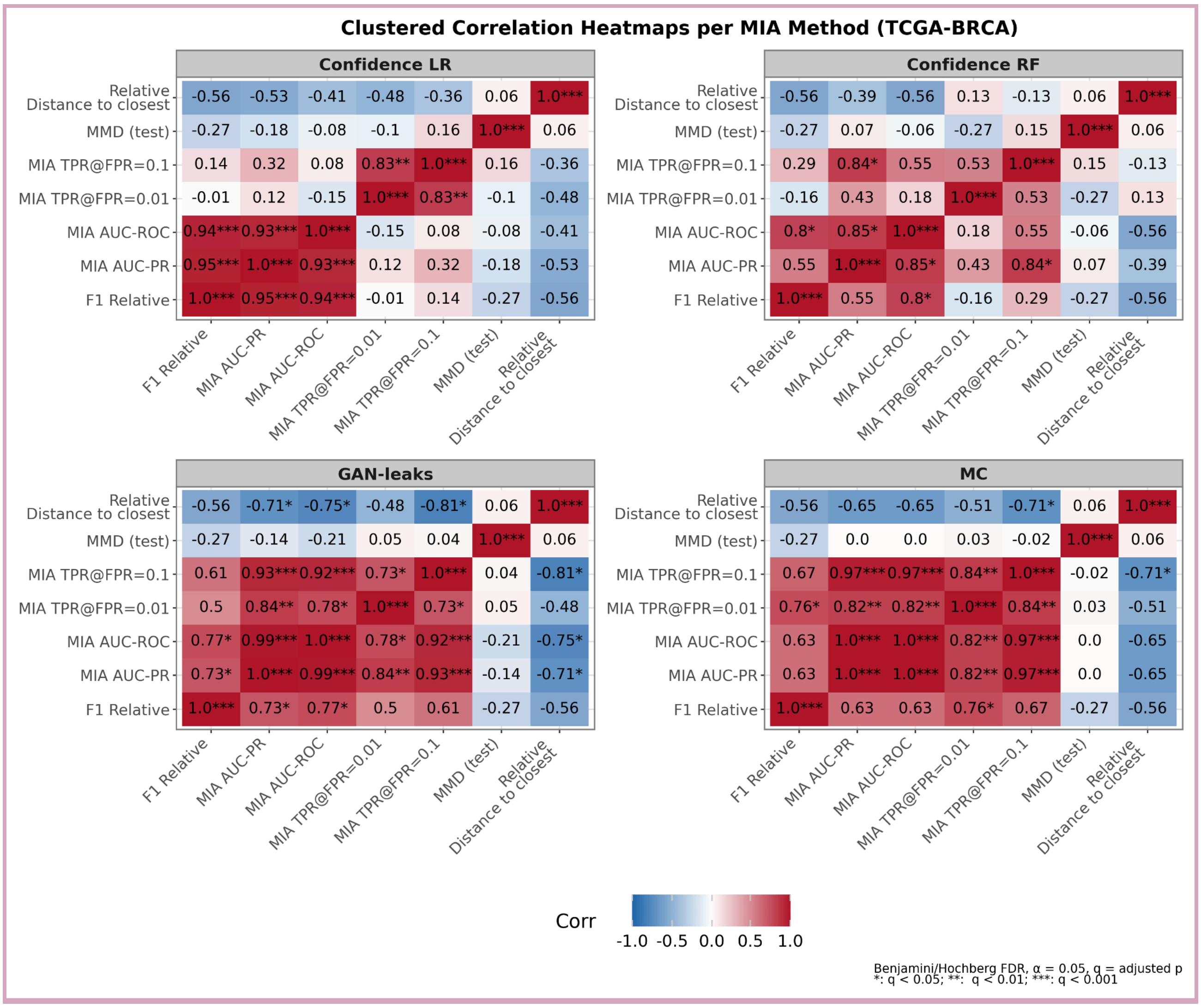


**Supp Figure S11: TGCA-BRCA Spearman correlation heatmap of MIA metrics with distance-to-closest, MMD (test) and F1 relative across four MIA methods.** Seven metrics were included, and statistical significance was assessed using the Benjamini–Hochberg procedure controlling the false discovery rate (FDR) at $\alpha = 0.05$. Significance levels are indicated as follows: *q < 0.05; ** q < 0.01; *** q < 0.001, where q represents the adjusted p-value.

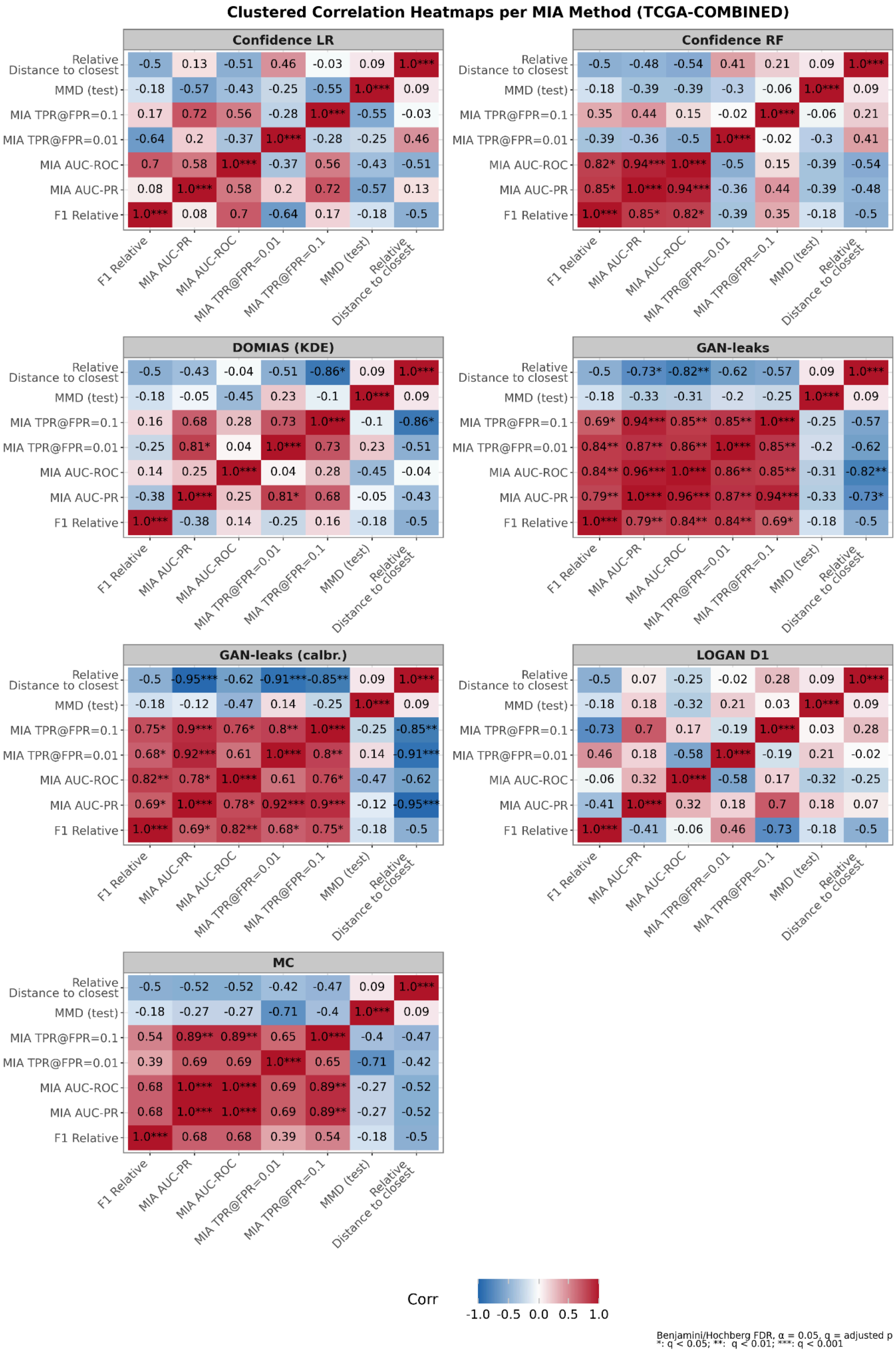
Clustered Correlation Heatmaps per MIA Method (TCGA-COMBINED)
Confidence LR
Relative Distance to closest -0.5 0.13 -0.51 0.46 -0.03 0.09 1.0***
MMD (test) -0.18 -0.57 -0.43 -0.25 -0.55 1.0*** 0.09
MIA TPR@FPR=0.1 0.17 0.72 0.56 -0.28 1.0*** -0.55 -0.03
MIA TPR@FPR=0.01 -0.64 0.2 -0.37 1.0*** -0.28 -0.25 0.46
MIA AUC-ROC 0.7 0.58 1.0*** -0.37 0.56 -0.43 -0.51
MIA AUC-PR 0.08 1.0*** 0.58 0.2 0.72 -0.57 0.13
F1 Relative 1.0*** 0.08 0.7 -0.64 0.17 -0.18 -0.5
F1 Relative, MIA AUC-PR, MIA AUC-ROC, MIA TPR@FPR=0.01, MIA TPR@FPR=0.1, MMD (test), Relative Distance to closest
Confidence RF
Relative Distance to closest -0.5 -0.48 -0.54 0.41 0.21 0.09 1.0***
MMD (test) -0.18 -0.39 -0.39 -0.3 -0.06 1.0*** 0.09
MIA TPR@FPR=0.1 0.35 0.44 0.15 -0.02 1.0*** -0.06 0.21
MIA TPR@FPR=0.01 -0.39 -0.36 -0.5 1.0*** -0.02 -0.3 0.41
MIA AUC-ROC 0.82* 0.94*** 1.0*** -0.5 0.15 -0.39 -0.54
MIA AUC-PR 0.85* 1.0*** 0.94*** -0.36 0.44 -0.39 -0.48
F1 Relative 1.0*** 0.85* 0.82* -0.39 0.35 -0.18 -0.5
F1 Relative, MIA AUC-PR, MIA AUC-ROC, MIA TPR@FPR=0.01, MIA TPR@FPR=0.1, MMD (test), Relative Distance to closest
DOMIAS (KDE)
Relative Distance to closest -0.5 -0.43 -0.04 -0.51 -0.86* 0.09 1.0***
MMD (test) -0.18 -0.05 -0.45 0.23 -0.1 1.0*** 0.09
MIA TPR@FPR=0.1 0.16 0.68 0.28 0.73 1.0*** -0.1 -0.86*
MIA TPR@FPR=0.01 -0.25 0.81* 0.04 1.0*** 0.73 0.23 -0.51
MIA AUC-ROC 0.14 0.25 1.0*** 0.04 0.28 -0.45 -0.04
MIA AUC-PR -0.38 1.0*** 0.25 0.81* 0.68 -0.05 -0.43
F1 Relative 1.0*** -0.38 0.14 -0.25 0.16 -0.18 -0.5
F1 Relative, MIA AUC-PR, MIA AUC-ROC, MIA TPR@FPR=0.01, MIA TPR@FPR=0.1, MMD (test), Relative Distance to closest
GAN-leaks
Relative Distance to closest -0.5 -0.73* -0.82** -0.62 -0.57 0.09 1.0***
MMD (test) -0.18 -0.33 -0.31 -0.2 -0.25 1.0*** 0.09
MIA TPR@FPR=0.1 0.69* 0.94*** 0.85** 0.85** 1.0*** -0.25 -0.57
MIA TPR@FPR=0.01 0.84** 0.87** 0.86** 1.0*** 0.85** -0.2 -0.62
MIA AUC-ROC 0.84** 0.96*** 1.0*** 0.86** 0.85** -0.31 -0.82**
MIA AUC-PR 0.79** 1.0*** 0.96*** 0.87** 0.94*** -0.33 -0.73*
F1 Relative 1.0*** 0.79** 0.84** 0.84** 0.69* -0.18 -0.5
F1 Relative, MIA AUC-PR, MIA AUC-ROC, MIA TPR@FPR=0.01, MIA TPR@FPR=0.1, MMD (test), Relative Distance to closest
GAN-leaks (calbr.)
Relative Distance to closest -0.5 -0.95*** -0.62 -0.91*** -0.85** 0.09 1.0***
MMD (test) -0.18 -0.12 -0.47 0.14 -0.25 1.0*** 0.09
MIA TPR@FPR=0.1 0.75* 0.9*** 0.76* 0.8** 1.0*** -0.25 -0.85**
MIA TPR@FPR=0.01 0.68* 0.92*** 0.61 1.0*** 0.8** 0.14 -0.91***
MIA AUC-ROC 0.82** 0.78* 1.0*** 0.61 0.76* -0.47 -0.62
MIA AUC-PR 0.69* 1.0*** 0.78* 0.92*** 0.9*** -0.12 -0.95***
F1 Relative 1.0*** 0.69* 0.82** 0.68* 0.75* -0.18 -0.5
F1 Relative, MIA AUC-PR, MIA AUC-ROC, MIA TPR@FPR=0.01, MIA TPR@FPR=0.1, MMD (test), Relative Distance to closest
LOGAN D1
Relative Distance to closest -0.5 0.07 -0.25 -0.02 0.28 0.09 1.0***
MMD (test) -0.18 0.18 -0.32 0.21 0.03 1.0*** 0.09
MIA TPR@FPR=0.1 -0.73 0.7 0.17 -0.19 1.0*** 0.03 0.28
MIA TPR@FPR=0.01 0.46 0.18 -0.58 1.0*** -0.19 0.21 -0.02
MIA AUC-ROC -0.06 0.32 1.0*** -0.58 0.17 -0.32 -0.25
MIA AUC-PR -0.41 1.0*** 0.32 0.18 0.7 0.18 0.07
F1 Relative 1.0*** -0.41 -0.06 0.46 -0.73 -0.18 -0.5
F1 Relative, MIA AUC-PR, MIA AUC-ROC, MIA TPR@FPR=0.01, MIA TPR@FPR=0.1, MMD (test), Relative Distance to closest
MC
Relative Distance to closest -0.5 -0.52 -0.52 -0.42 -0.47 0.09 1.0***
MMD (test) -0.18 -0.27 -0.27 -0.71 -0.4 1.0*** 0.09
MIA TPR@FPR=0.1 0.54 0.89** 0.89** 0.65 1.0*** -0.4 -0.47
MIA TPR@FPR=0.01 0.39 0.69 0.69 1.0*** 0.65 -0.71 -0.42
MIA AUC-ROC 0.68 1.0*** 1.0*** 0.69 0.89** -0.27 -0.52
MIA AUC-PR 0.68 1.0*** 1.0*** 0.69 0.89** -0.27 -0.52
F1 Relative 1.0*** 0.68 0.68 0.39 0.54 -0.18 -0.5
F1 Relative, MIA AUC-PR, MIA AUC-ROC, MIA TPR@FPR=0.01, MIA TPR@FPR=0.1, MMD (test), Relative Distance to closest
Corr
-1.0 -0.5 0.0 0.5 1.0
Benjamini/Hochberg FDR, α = 0.05, q = adjusted p
*: q < 0.05; **: q < 0.01; ***: q < 0.001

**Supp Figure S12: TGCA-COMBINED Spearman correlation heatmap of MIA metrics with distance-to-closest, MMD (test) and F1 relative across four MIA methods.** Seven metrics were included, and statistical significance was assessed using the Benjamini–Hochberg procedure controlling the false discovery rate (FDR) at $\alpha = 0.05$. Significance levels are indicated as follows: *q < 0.05; ** q < 0.01; *** q < 0.001, where q represents the adjusted p-value.

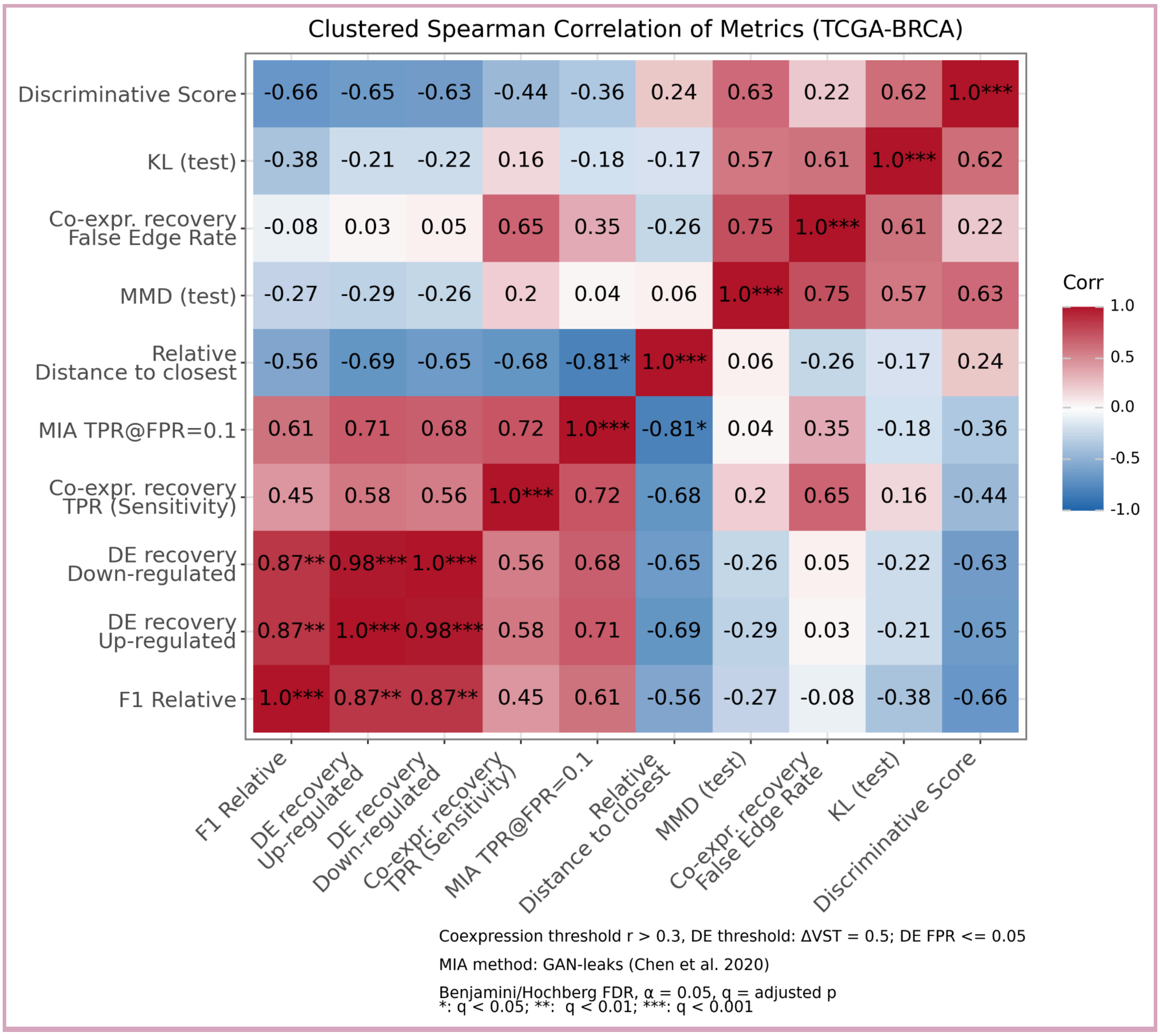


**Supp Figure S13: TGCA-BRCA Spearman correlation heatmap of selected metric from four evaluation axes: fidelity, downstream utility, biological plausibility and privacy.** Ten metrics were included, and statistical significance was assessed using the Benjamini–Hochberg procedure controlling the false discovery rate (FDR) at $\alpha = 0.05$. Significance levels are indicated as follows: *q < 0.05; ** q < 0.01; *** q < 0.001, where q represents the adjusted p-value.

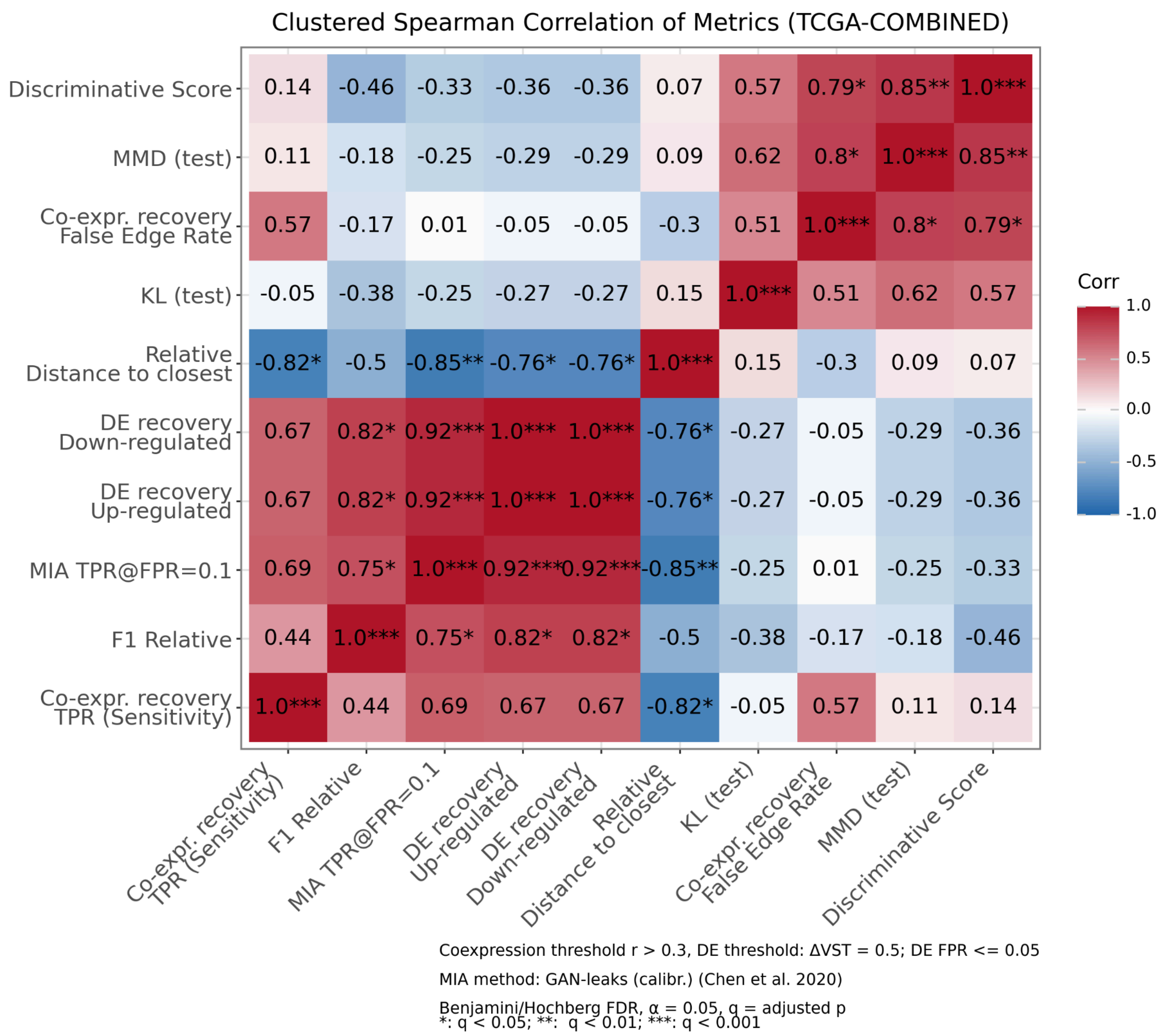


**Supp Figure S14: TGCA-COMBINED Spearman correlation heatmap of selected metric from four evaluation axes: fidelity, downstream utility, biological plausibility and privacy.** Ten metrics were included, and statistical significance was assessed using the Benjamini–Hochberg procedure controlling the false discovery rate (FDR) at $\alpha = 0.05$. Significance levels are indicated as follows: *q < 0.05; ** q < 0.01; *** q < 0.001, where q represents the adjusted p-value.

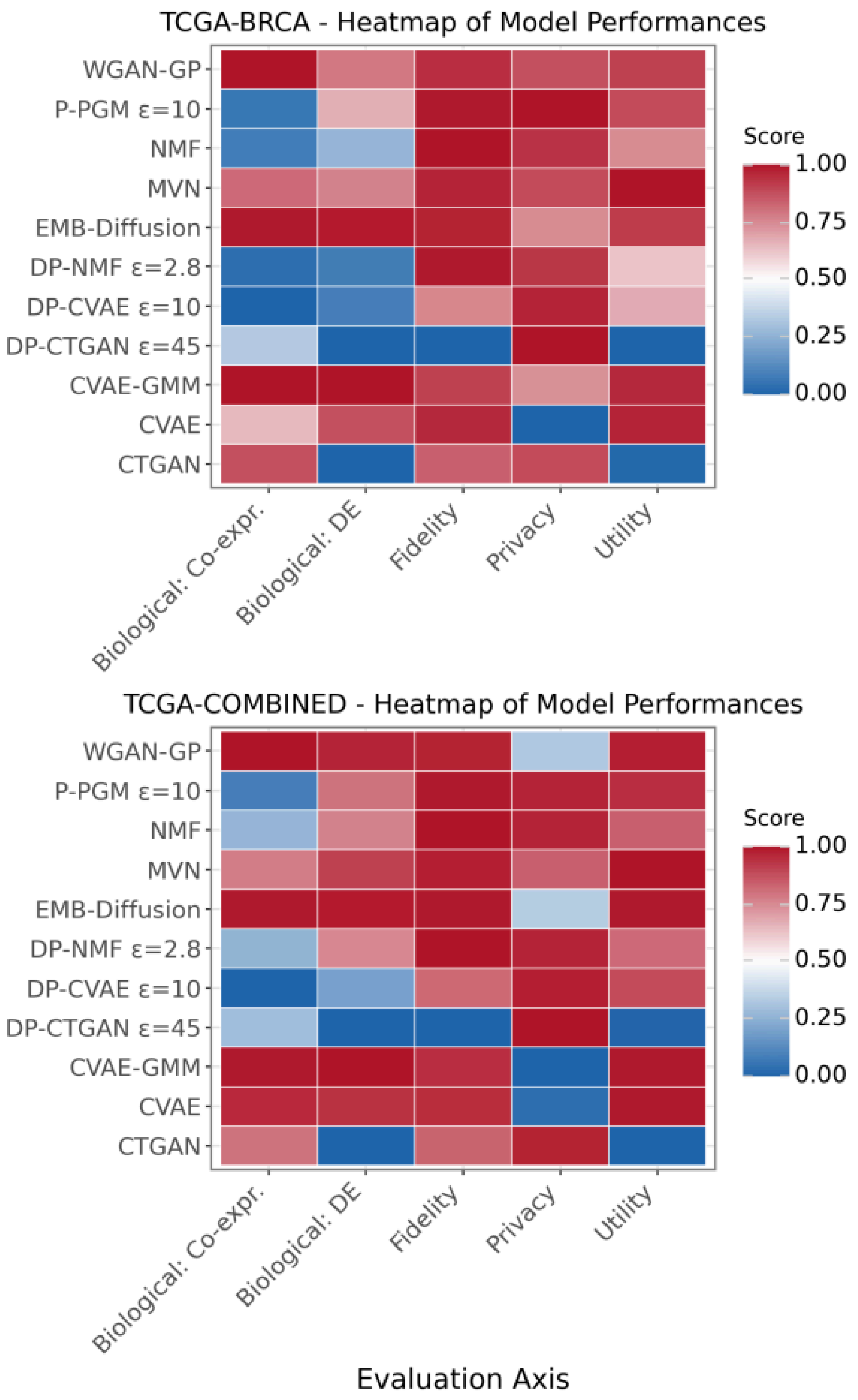


**Supp Figure S15: Model comparison heatmap for TCGA-BRCA and TCGA-COMBINED datasets**. Each row correspond to generative models (alphabetically ordered), and columns show selected metrics: **Fidelity** (inverted Maximum Mean Discrepancy on test data), **Utility** (relative F1 score), **Biological: DE** (average preservation TPR of differentially expressed genes), **Biological: Co-expression** (co-expression recall), and **Privacy** (logit-transformed TPR@FPR=0.1, scaled and inverted so higher values indicate better privacy protection). All metrics were min–max normalized to range 0-1, so higher values correspond to better performance. Tile colors range from blue (lower scores) through white (intermediate) to red (higher scores), summarizing model behavior across multiple performance dimensions.

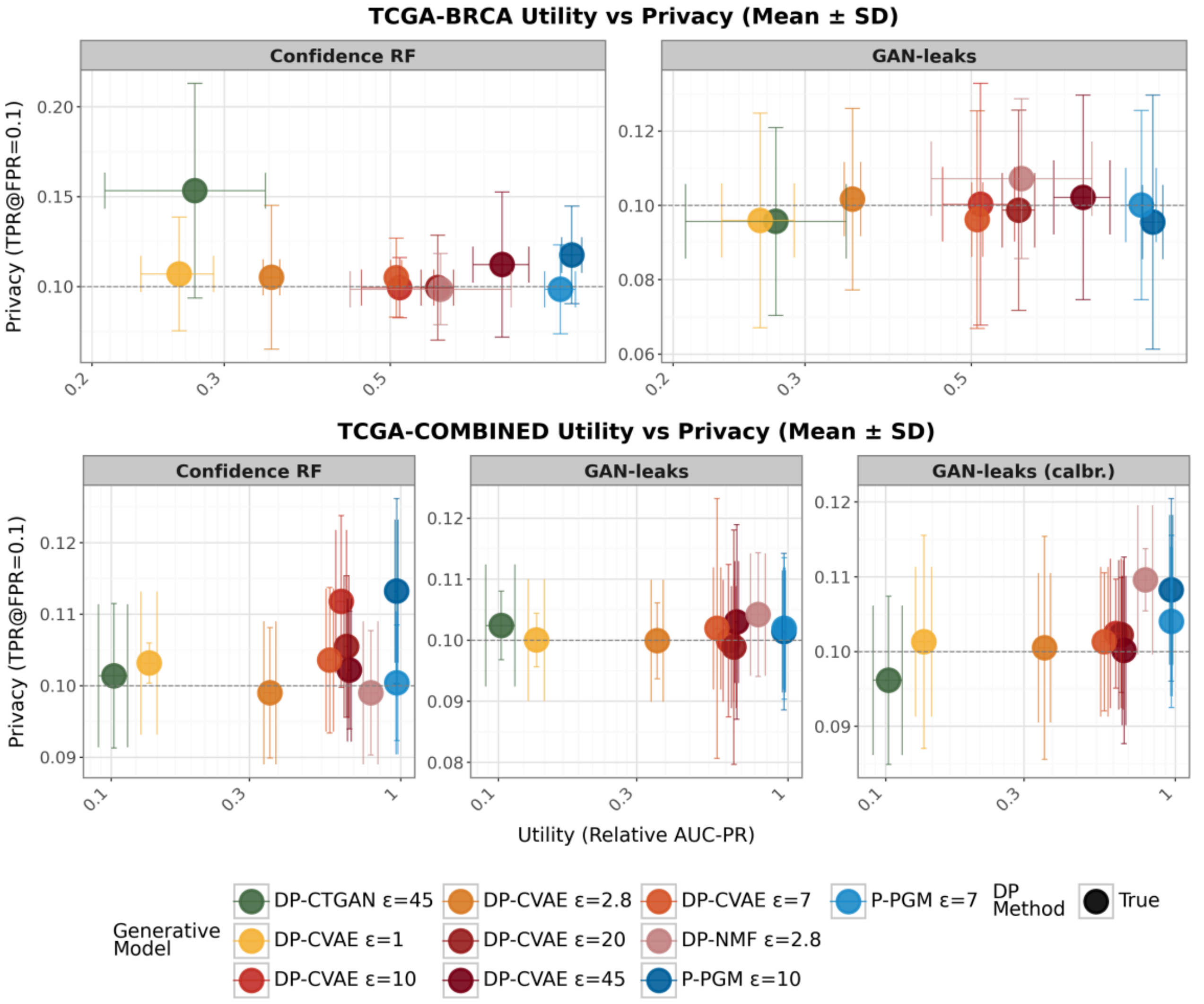


**Supp Figure S16:** Utility - privacy trade-off for DP-CVAE across privacy budgets ($\epsilon$), compared with other formally differentially private models with matched $\epsilon$ on TCGA-BRCA and TCGA-COMBINED datasets.

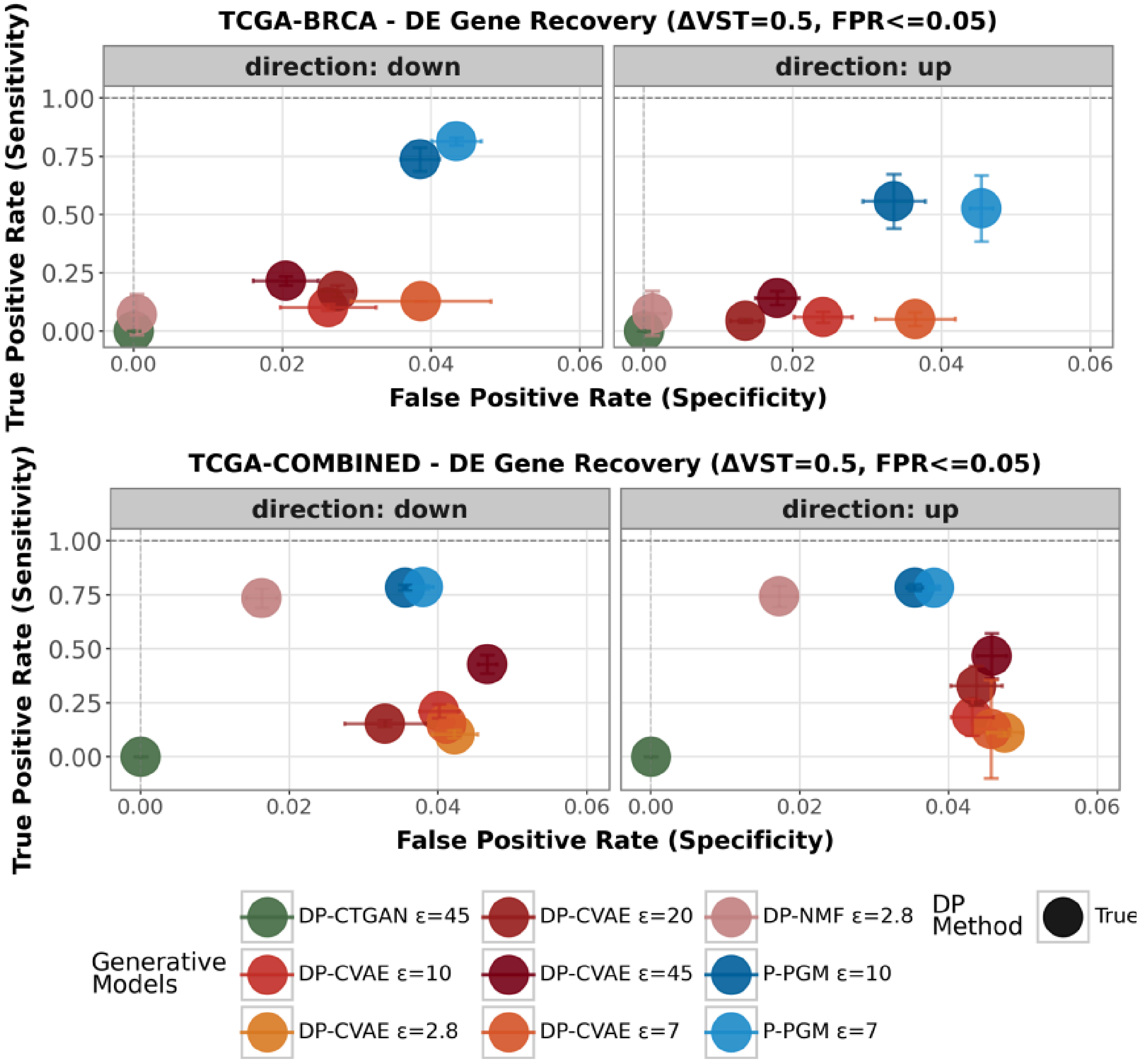


**Supp Figure S17:** Differential expression recovery for DP-CVAE across privacy budgets ($\epsilon$) compared with other formally DP models with matched $\epsilon$ on TCGA-BRCA and TCGA-COMBINED datasets.

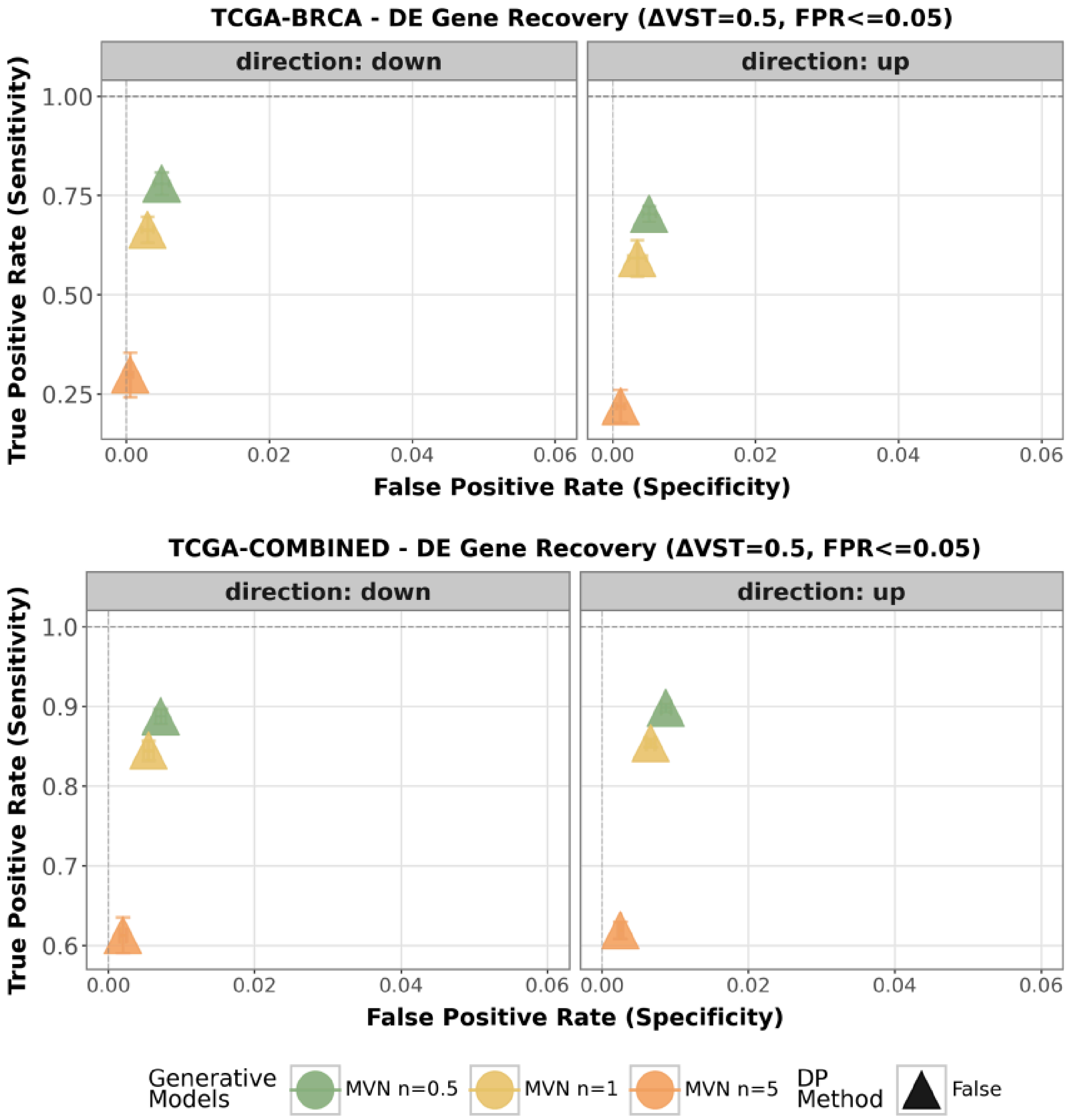


**Supp Figure S18:** Impact of Gaussian noise level on MVN model differential expression (DE) recovery foTCGA-BRCA and TCGA-COMBINED datasets.
